# Near Atomic Structure of an Atadenovirus Reveals a Conserved Capsid-Binding Motif and Intergenera Variations in Cementing Proteins

**DOI:** 10.1101/2020.07.24.220046

**Authors:** Roberto Marabini, Gabriela N. Condezo, Josué Gómez-Blanco, Carmen San Martín

## Abstract

Little is known about the basic biology of non-human adenoviruses, which could be alternative vectors free of issues posed by preexisting immunity to human adenoviruses. We present the cryo-EM structure of a lizard atadenovirus, LAdV-2, at 3.4 Å resolution. This is the first high resolution structure of an adenovirus with non-mammalian host, and of an adenovirus not belonging to the Mastadenovirus genus. Atadenovirus capsids contain genus specific proteins LH3, p32k, and LH2, and are more thermostable than the more studied human adenoviruses. We find a large conformational difference in the internal vertex protein IIIa between mast- and atadenoviruses, induced by the presence of an extended polypeptide in the region. This polypeptide, as well as α-helical clusters located beneath the icosahedral facet, likely correspond to proteins LH2 and p32k. The external genus specific protein LH3, with a trimeric β-helix fold typical of bacteriophage host attachment proteins, contacts the hexon shell surface via a triskelion structure identical to that used by protein IX in human AdV, revealing a conserved capsid-binding motif and a possible gene duplication event. Altogether, this work shows how the network of minor coat proteins differs between AdV genera and relates to virus evolution and capsid stability properties.

## Main

Adenoviruses (AdVs) are non-enveloped, dsDNA viruses with a ∼95 nm, *pseudo*T=25 icosahedral capsid. Because of their widespread use as experimental vectors, most of our current knowledge on AdVs comes from the study of only a few types, in particular human AdV type 5 (HAdV-C5). Each HAdV-C5 capsid facet has 12 trimers of the major coat protein, hexon. At each vertex, five penton base subunits form the penton base, from which the receptor binding, trimeric fibers project. Minor coat proteins IIIa, VI and VIII on the inner capsid surface, and IX on the outer surface, complete the intricate network of interactions required for capsid assembly and stabilization (Dai *et al.*, 2017; Liu *et al.*, 2010). Positively charged core proteins V, VII and μ are packed together with the 35 kbp, linear dsDNA genome within the capsid (Pérez-Berná *et al.*, 2015). AdV particle size and composition have represented a challenge for structural biology techniques; multiple proteolytic cleavages during maturation and lack of icosahedral ordering in a large part of the capsid components further complicate the situation (Mangel and San Martín, 2014; San Martín, 2012).

Recombinant human AdVs (HAdVs) are widely used as vehicles for gene transfer, oncolysis and vaccination. However, their successful use in the clinic requires surmounting hurdles such as pre-existing immunity in the population, or tropism control. A possible approach to overcome these problems is the use of non-human AdV (Lopez-Gordo *et al.*, 2014). There are currently five approved AdV genera: mastadenoviruses (infecting mammals); aviadenoviruses (birds); atadenoviruses (reptiles, ruminants and birds); siadenoviruses (amphibians, birds and reptiles); and ichtadenovirus (a single isolate from white sturgeon) (Harrach *et al.*, 2011). Structural characterization of non-human AdV is so far limited to medium resolution studies on one bat (BtAdV 250-A) (Hackenbrack *et al.*, 2016), one canine (CAdV-2) (Schoehn *et al.*, 2008), one ovine (OAdV-7) (Pantelic *et al.*, 2008), and one snake (SnAdV-1) AdV (Menéndez-Conejero *et al.*, 2017), as well as a 4 Å resolution report on incomplete bovine adenovirus (BAdV-3) capsids (Cheng *et al.*, 2014). All these are mastadenoviruses, except for OAdV-7 and SnAdV-1, which are atadenoviruses. There are no high resolution data on the complete virion structure for any AdV not belonging to the *Mastadenovirus* genus.

All AdV genera contain a common set of genes involved in DNA replication, DNA encapsidation, and viral particle formation (Davison *et al.*, 2003). Minor coat protein IX and core protein V are virion components unique to mastadenoviruses. In atadenoviruses, genus-specific proteins LH2, LH3 and p32K have also been found in the virion (Gorman *et al.*, 2005; Menéndez-Conejero *et al.*, 2017; Pantelic *et al.*, 2008; Pénzes *et al.*, 2014). Similarly to protein IX in mastadenoviruses, LH3 is located on the outer capsid surface, while p32k and LH2 have been tentatively assigned to positions on the inner capsid surface, and may be substituting for protein V (Gorman *et al.*, 2005; Menéndez-Conejero *et al.*, 2017; Pantelic *et al.*, 2008). LH3 was previously referred to as E1B 55K, due to limited sequence homology with the gene placed in a similar position in the HAdV genome (Vrati *et al.*, 1996). However, HAdV protein E1B 55K is not part of the virion, but is expressed in infected cells where it carries a large variety of functions, including promotion of genome replication and transcription, degradation of antiviral factors, or deregulation of the cell cycle (Hidalgo *et al.*, 2019).

Medium resolution cryo-EM combined with crystallography showed that, in SnAdV-1, LH3 has a trimeric β-helix fold typical of bacteriophage host attachment proteins, and indicated extensive contacts between LH3 and hexons surrounding the icosahedral and local 3-fold axes in the capsid, corroborating a role for LH3 in the high thermostability of atadenovirus capsids (Menéndez-Conejero *et al.*, 2017; Pantelic *et al.*, 2008). However, this study did not solve the full extent of the LH3 protein, as the form crystallized lacked ∼30 residues from the N-terminus. We now present the first near atomic resolution structure of an atadenovirus, LAdV-2, showing in detail how the network of minor coat proteins differs between genera.

## Results

### Structure determination of LAdV-2

The 3.4 Å resolution cryo-EM map of LAdV-2 (**Fig. 1A and S1A-B**) showed the expected particle morphology, with a *pseudo*T=25 icosahedral geometry, 940 Å diameter, 12 hexon trimers per facet, and outer protrusions corresponding to the genus specific protein LH3 (Menéndez-Conejero *et al.*, 2017). The penton base density was weaker than the hexon shell, indicating partial penton loss (**Fig. S1C**). Low occupancy reflected in lower resolution (∼4 Å, **Fig. S1D**) in the penton region, and higher B-factors in the refined model (**Fig. S1E**). Similarly to previous studies on mastadenoviruses, no indication of ordered structures or concentric shells was observed in the core (**Fig. S1C**). We traced over 13,400 residues in the icosahedral asymmetric unit (AU), corresponding to: four independent hexon trimers; one penton monomer; four independent copies of LH3; one copy of IIIa; and two independent copies of protein VIII. We observed density inside all hexons that we interpret as the N-terminal peptide of protein VI (nine copies per AU) and protein VII (three copies), based on the fitted chain length and on the latest HAdV-C5 structure (Dai *et al.*, 2017), although we were unable to unequivocally assign residue identities. Additionally, 255 residues were modelled as poly-alanine chains in density where it was not possible to assign a sequence (**Fig. 1B and Table S1**).

**Figure 1.**
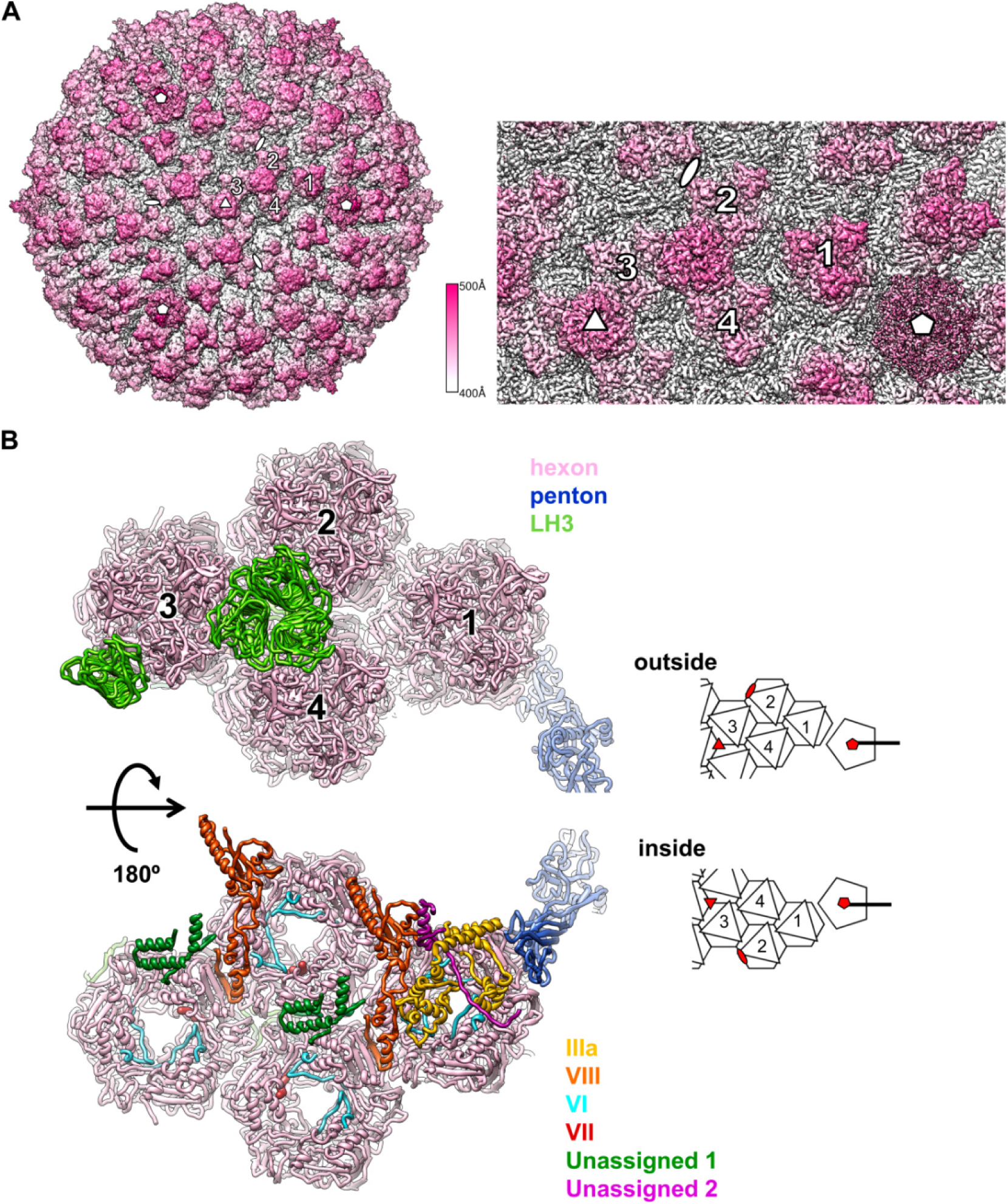
LAdV-2 cryo-EM map and molecular model. **(A)** Right: surface rendering of the LAdV-2 3D map coloured by radius from white to pink, as indicated by the colour scale. A zoom in on the area corresponding to the icosahedral AU is shown at the right. The four hexon trimers in an AU are numbered 1-4. White symbols indicate the icosahedral 5-fold (pentagon), 3-fold (triangle) and 2-fold (oval) symmetry axes. **(B)** Ribbon representation of the proteins traced in the AU coloured as indicated by the legend at the right. Two views are provided, as seen from outside (top) or inside (bottom) the capsid, with a cartoon at the right hand side for guidance. In the cartoon, icosahedral symmetry axes are indicated by red symbols.

### Main capsid proteins: hexon and penton

The LAdV-2 hexon protein is shorter than its HAdV-C5 counterpart (909 *vs.* 952 residues), and could be traced almost in its entirety for all 12 monomers in the AU (**Table S1, Fig. S2**). As expected from previous studies (Liu *et al.*, 2010; Xu *et al.*, 2007; Yu *et al.*, 2017), the overall structure of hexon, with a double jelly roll normal to the capsid surface, and extensive loops that imbricate to form the trimer towers, is conserved (**Fig. 2A**). The root mean square distance (RMSD) for all Cα atoms when comparing to HAdV-C5 hexon (PDB ID 6B1T) is ∼4 Å. The largest differences between mast- and atadenovirus hexons occur at the towers (**Fig 2B**). In HAdVs, mobile loops on the tower surface are formed by hypervariable regions (HVR) that contribute to define the virus serotype (Crawford-Miksza and Schnurr, 1996). In the most recent HAdV-C5 cryo-EM structure, the hexon model contains several gaps in these loops (Ala138-Gln164; Gln253-Leu258; Thr273-Asn279; Thr433-Asn437) where the chain could not be traced (Dai *et al.*, 2017). The loops in the LAdV-2 hexon tower are shorter than those in HAdV-C5, and could be modelled without gaps even if the density was in some regions slightly fragmented. In LAdV-2, the hexon towers adopt very similar conformations in all hexon monomers (RMSD < 2 Å for all Cα atoms) except for Ala374-Ala375, located at the valley formed by the trimer towers (**Fig. 2A, C**). In HAdV-C5, residues at this valley are involved in contacts with coagulation factors (Alba *et al.*, 2009). It is believed that the exposed HVRs have evolved in response to the evolutionary pressure of the host immune system. Simpler loops might correlate with a different immune system in reptiles (Zimmerman, 2016).

**Figure 2.**
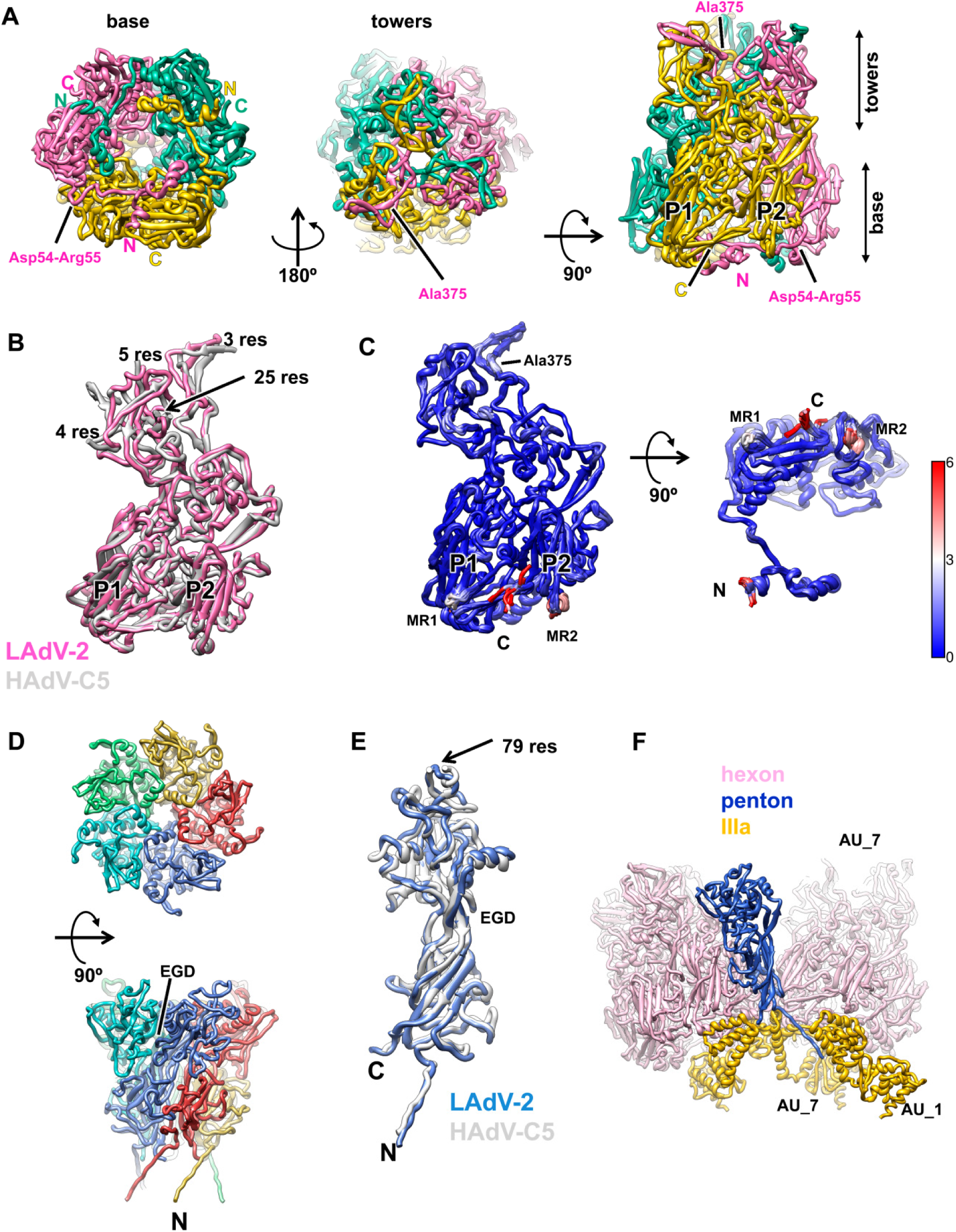
LAdV-2 main capsid proteins: hexon and penton. **(A)** Structure of the hexon trimer as seen from inside (left) or outside (center) the capsid, and in a side view (right). The capsomer base (formed by the double jelly roll domains) and towers are indicated, as well as the **N**- and **C**-termini of each monomer. The two β-barrels forming the double jelly roll are labelled **P1** and **P2. (B)** Superposition of the LAdV-2 and HAdV-C5 (PDB ID 6B1T) hexon monomers. The length in residues **(res)** of the HAdV-C5 untraced loops is indicated. **(C)** Superposition of the twelve hexon monomers in the LAdV-2 AU, coloured by RMSD according to the scale at the right hand side (in Å). **MR1** and **MR2** indicate the highest mobility regions, other than the **N**- and **C**-termini. **(D)** Structure of the penton base pentamer as seen from outside (top panel) the capsid, and in a side view (bottom). The **N**-terminus of one monomer and the location of the EGD sequence are indicated. **(E)** Superposition of the LAdV-2 and HAdV-C5 penton base monomers. The length in residues **(res)** of the HAdV-C5 penton RGD loop is indicated, as well as the location of the EGD sequence and the **N**- and **C**-termini of the LAdV-2 protein. **(F)** Interactions of one penton base monomer with other proteins. AU_1 and AU_7 indicate molecules in neighbouring AUs labelled as in **Fig. S3**.

All other major differences (RMSD > 2 Å) between hexon monomers in LAdV-2 are located at the trimer base, and comprise residues Met1-Glu2 (N-term), Thr299-Gln301 (mobile region 1, MR1), Val849-Ala852 (MR2), and Ser904-Ala908 (C-term) (**Fig. 2C**). In HAdV-C5, the N- and C-terminal regions were also observed to adopt different conformations in the different monomers, contributing to the quasi-equivalent interactions, but these regions were more extensive in the human virus (6 and 7 aminoacids respectively) (Liu *et al.*, 2010) (**Tables S2-S8**). The other two hexon mobile regions (MR1 and MR2) are located at alternate vertices of the pseudohexagonal base, one below each beta-jelly roll, and are involved in various hexon-hexon interactions, as well as hexon interactions with penton base or minor coat proteins IIIa and VIII (mostly MR2) (**Fig. 3A; Tables S2-S8**). Residues Asp54-Arg55 (**Fig. 2A**), also located on the inner hexon surface, can establish a pair of contiguous salt bridges at each of two local 2-fold symmetry axes at the facet edges: between hexon 1 (H1) in the AU and its 5-fold symmetry neighbour, and between H4 in the AU and H2 in the next facet (H2_AU6 in **Fig. S3** and **Table S3**, TT interactions). These residues are conserved in HAdV-C5 (Asp59-Arg60), suggesting that ionic interactions may be important for assembly of face plates onto the icosahedron. This interaction, however, is not present at the icosahedral 2-fold axis. The same two residues in a different monomer of H1 and H4 establish another possible pair of electrostatic interactions with the internal minor coat protein VIII (see below).

**Fig. 3.**
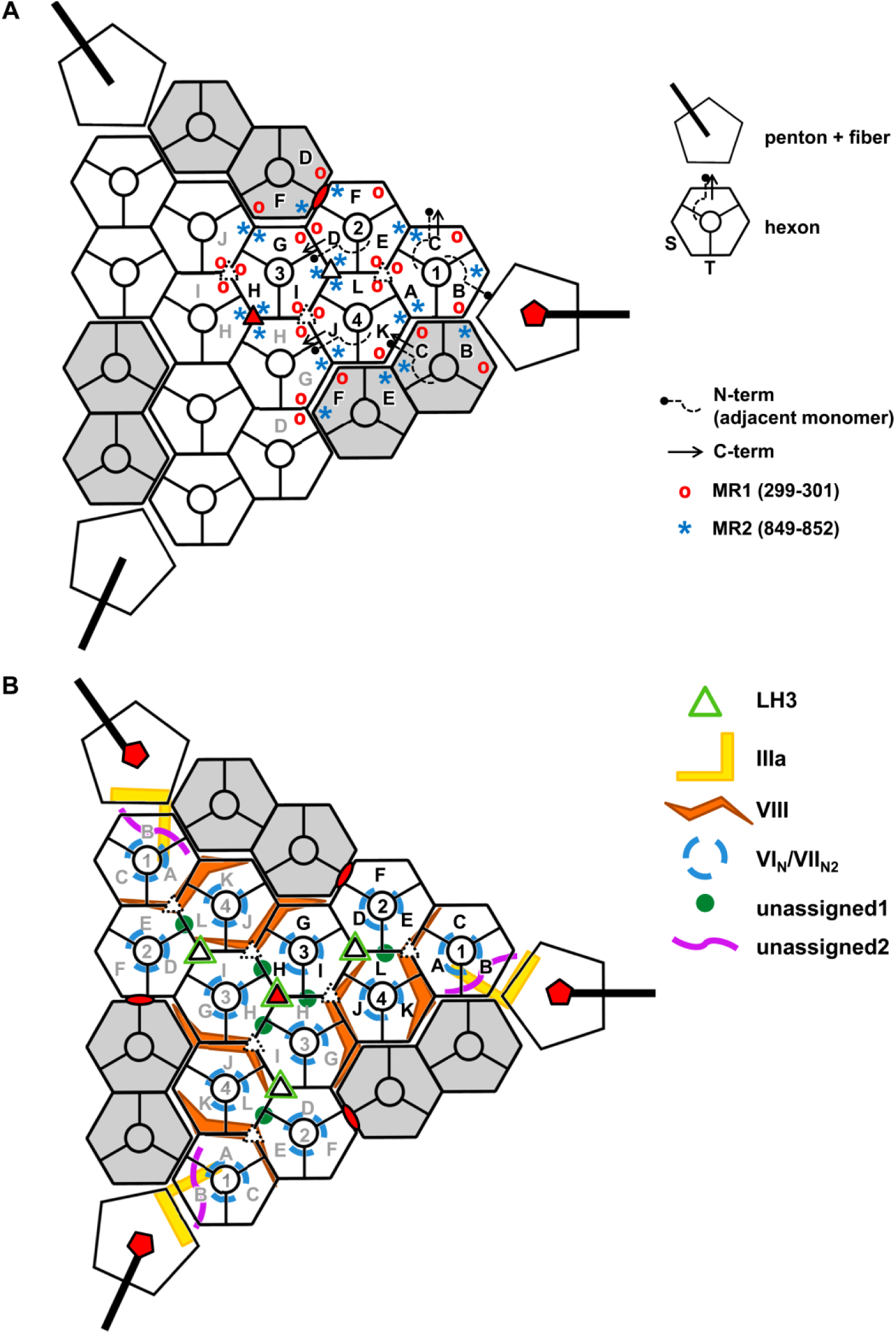
Schematics illustrating the LAdV-2 protein networks. Hexons 1-4 in one AU are depicted in white and labelled with black text; those in neighbouring AUs in the same icosahedral facet are in white with grey labels; and those in adjacent facets are in grey. Letters **A-L** identify different polypeptide chains as in **Tables S2-S9**. Red symbols indicate the icosahedral 5-fold (pentagon), 3-fold (triangle) and 2-fold (oval) symmetry axes. Closed and broken white triangles indicate two different local 3-fold symmetry axes. **(A)** Hexon conformational adaptability: location of mobile hexon regions in the capsid. Hexon mobile regions MR1 and MR2 in one AU and its immediate neighbours are indicated. A few hexon N- and C-termini are depicted as an example. In the hexon schematics legend, **S** designates the facet of the hexon pseudo-hexagonal base formed by the two β-barrels in a single monomer, and **T** indicates the facet of the hexon pseudo-hexagonal base formed by two β-barrels belonging to two adjacent hexon monomers (Liu *et al.*, 2010). **(B)** Location of the minor coat proteins. Only LH3 is located on the outer capsid surface.

In spite of the low occupancy, the penton base polypeptide chain could be traced in its entirety, including its N-terminal arm (Met1-Gly16) that stretches towards the viral core and corresponds to the same feature formed by residues 37-51 in the HAdV-C5 penton base (**Fig. 2D-E and S2**). Similarly to hexon, the penton base protein is shorter in LAdV-2 than in HAdV-C5 (451 vs. 571 residues), and the main differences reside in the outer surface loops, with the rest of the structure very close to that of HAdV-C5 (2.1 Å RMSD) (Liu *et al.*, 2010; Zubieta *et al.*, 2005). Most notably, the flexible, 80-residue long loop containing the RGD sequence motif in HAdV-C5 is not present in LAdV-2 (**Fig. 2E**). The closest sequence pattern in the LAdV-2 penton would be EGD at residues 133-135, but these are located at the interface between penton monomers, and therefore not available for interactions with cell receptors (**Fig. 2E**). Lack of the RGD integrin binding motif suggests an atadenovirus internalization mechanism different from the best characterized HAdVs.

Penton base monomers are arranged in an oblique fashion around the 5-fold symmetry axis, facilitating their bonding with multiple neighbouring molecules. Each penton base monomer interacts with the two neighbouring peripentonal hexon trimers (SP interactions, **Table S5**). Additionally, the N-terminal arm interacts with two different copies of protein IIIa, while the C-terminal residue is positioned within reach of interactions with a third IIIa molecule (**Table S6 and Fig. 2F**). In HAdV-5, the first 37 residues could not be traced, and were proposed to plunge into the viral core. Since here we can trace the penton base chain starting from residue 1, this penton-core interaction would not be present in LAdV-2.

The symmetry mismatch between the trimeric fibres and the pentameric penton base is worsened in LAdV-2 by the presence of two different fibre proteins, one of them attached in triplets to some of the vertices (Pénzes *et al.*, 2014). Consequently, fibres were not traced in this icosahedrally averaged map.

### External minor coat proteins: LH3

There are twelve copies of protein LH3 per icosahedral facet, organized in four trimers. One of these is located at the icosahedral 3-fold axis, while the other three occupy the local 3-fold axes between hexons 2, 3 and 4 in each AU (**Fig. 1B, 3B**). The structure of a stable fragment of recombinant SnAdV-1 LH3 (residues 28-373) was previously solved by crystallography, showing that it folds as a right-handed β-helix with three strands per turn, an architecture strikingly similar to bacteriophage tailspikes (Menéndez-Conejero *et al.*, 2017). LH3 in LAdV-2 is three residues shorter (370 *vs.* 373), and has a 62.2% sequence identity with the SnAdV-1 protein, according to the experimentally determined sequence (Menéndez-Conejero *et al.*, 2017). The cryo-EM map of LH3 in its native environment (the viral particle) allowed us to trace most of the chain in all four positions in the AU; only the first 3 residues lacked density in our LAdV-2 cryo-EM map (**Table S1**). The overall structure of LAdV-2 LH3 is very similar to that of SnAdV-1 LH3 in the β-helix domain (1.2 Å RMSD for 336 Cα atoms; **Fig. 4A**). A loop formed by residues Asp155-Ser162 in the outer trimer surface was absent in most of the SnAdV-1 LH3 crystal structures, indicating mobility, but could be unequivocally traced in LAdV-2, where it is fixed in place by interactions with the surrounding hexons (**Fig. 4A, Fig. S4A, Table S9**).

**Figure 4.**
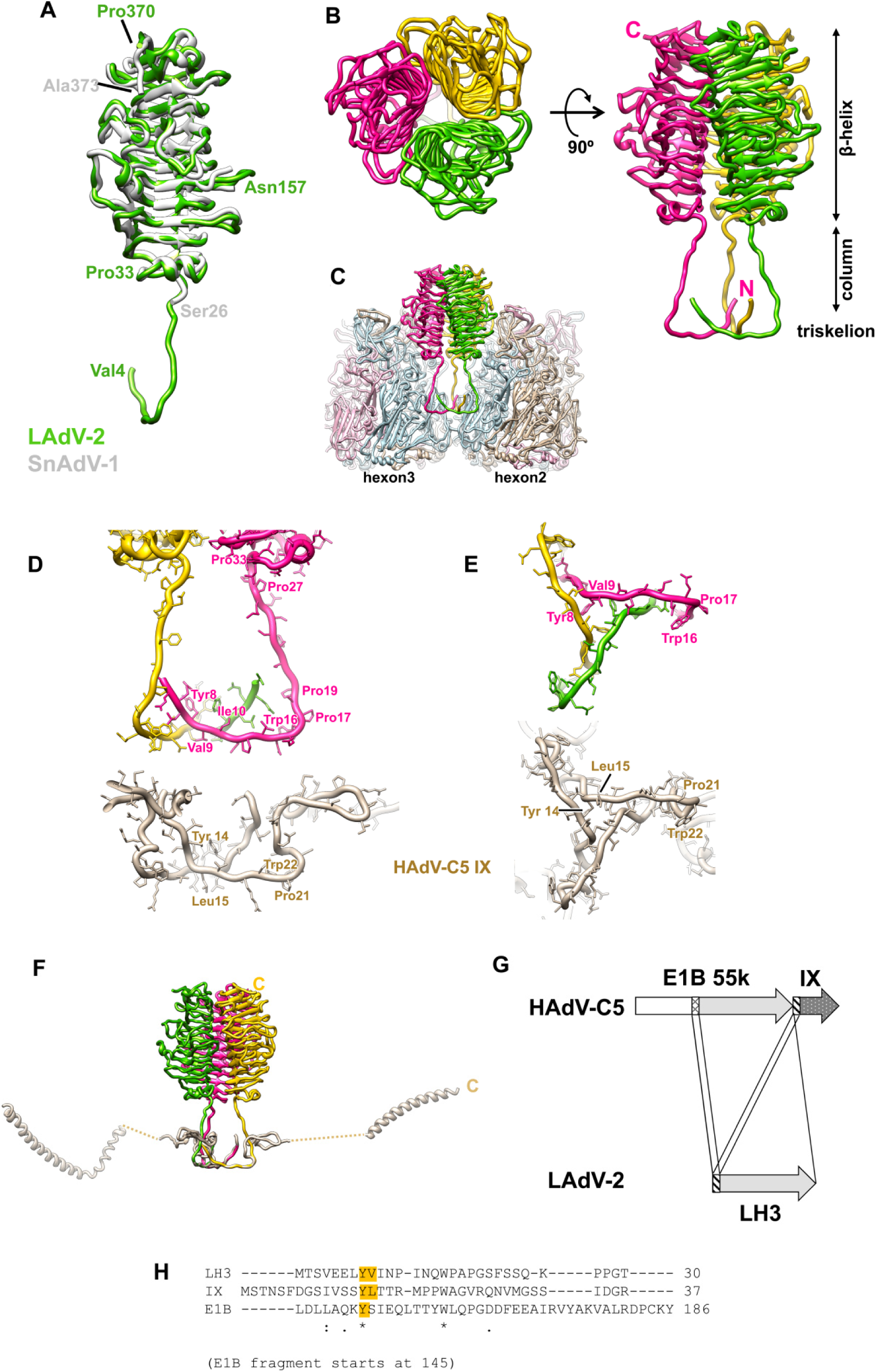
LAdV-2 external minor coat proteins: LH3. **(A)** Superposition of the LAdV-2 and SnAdV-1 (PDB ID 5G5O) LH3 structures. The first and last traced residues in each case are indicated, as well as the positions of a few other residues described in the main text. **(B)** Overall view of the LH3 trimer structure seen from outside the capsid (left) and in a side view (right). The **N-** and **C-**termini of the pink subunit are labelled, as well as the **β-helix, column** and **triskelion** domains. **(C)** Hexons 2 and 3 are represented to show the LH3 trimer in the capsid context. The monomers in each neighbouring hexon trimer are coloured light blue, light pink and tan. **(D)** Comparison between the triskelions formed by LAdV-2 LH3 and HAdV-C5 protein IX, in a side view similar to that in (C). Residues mentioned in the text are labelled for one monomer in each trimer. **(E)** As in (D), but in a view from the capsid surface. In this view, the LH3 column domains would travel away from the reader. **(F)** The LH3 trimer is overlapped with the HAdV-C5 protein IX trimer to show the large difference in the fold beyond the triskelion domain. The **C-** terminus for one monomer of each protein is labelled. **(G)** Cartoon schematizing the positions of the E1B-55K and IX genes in the HAdV-C5 genome, and the similarities between these proteins and LAdV-2 LH3. Each arrow represents one gene. Similarly shaded regions are connected by lines to indicate sequence similarity. **(H)** Sequence alignment of the LH3 and IX triskelion regions and a region of the E1B 55K protein sequence starting at residue 145. Hydrophobic residues at the triskelion core are highlighted in orange. Numbers at the right indicate the position of the last residue shown in the protein sequence.

We can now see that each LH3 monomer has an extended N-terminal domain that lays on the capsid surface at the valley formed by three surrounding hexon trimers, and after bending almost at a square angle goes up forming a column that reaches the β-helix domain (**Fig. 4A-C**). The column domain presents certain mobility, as shown by its weak density in the cryo-EM map (**Fig. S4B**). A region rich in proline and glycine residues (27-PPGTLLPG-34) allows the formation of a sharp bend when the flexible column gives way to the β-helix domain (**Fig. 4A and 4D**).

The three N-terminal domains (4-VEELYVINPINQWPAP-19) in each LH3 trimer join to form a triskelion-shaped joint on the surface of the valley between three hexon capsomers (**Fig. 4B-4C**).This structure is highly similar to the one formed by the N-terminus of polypeptide IX in HAdV-C5, which occupies the same positions in the capsid, stabilizing the nine central hexons in each capsid facet (**Fig. 3B and 4D-E**) (Liu *et al.*, 2010). LH3 residues Leu7, Tyr8, Val9 and Ile10 from each monomer form the triskelion hydrophobic core. A series of prolines (Pro12, Pro17, Pro19) facilitate the sharp bend from the triskelion to the column (**Fig. 4D**). A hydrophobic core (Tyr14-Leu15) also underpins the polypeptide IX triskelion in HAdV-C5 (Liu *et al.*, 2010). A conserved tryptophan (Trp16 in LH3, Trp22 in IX) is located at the outermost vertices of the triskelion, also with a nearby proline (Pro21) in IX (**Fig. 4D-E**). A core of two hydrophobic residues, as well as the tryptophan, are conserved in atadenoviruses (**Fig. S4C**). Sequence alignment of the complete LH3 and IX proteins does not detect the similarity at the triskelion region. However, the structure we present here reveals that such region is conserved (**Fig. 4D-E**).

Previous evidence has shown that the triskelion is critical for incorporation of both protein IX and LH3 to the capsid during assembly (Pantelic *et al.*, 2008; Rosa-Calatrava *et al.*, 2001; Vellinga *et al.*, 2005). Conversely, a large variability is tolerated for the rest of the “triskelion carrying” proteins: in HAdV-C5 IX, long, unstructured regions travel on the capsid surface all the way from the triskelions on the central plate of the facet to a C-terminal helix at the capsid edges **(Fig. 4F)** (Liu *et al.*, 2010). In non-human mastadenovirus IX proteins, shorter connecting regions climb away from the capsid surface and end in a helix directly on top of the triskelion (Cheng *et al.*, 2014; Hackenbrack *et al.*, 2016; Schoehn *et al.*, 2008). Atadenovirus LH3 has also a short connecting region that moves perpendicularly to the capsid surface (the column domain), but ends in a completely different fold (the β-helix) of the C-terminal domain (**Fig. 4F**).

It is now interesting to consider the genomic context for proteins IX in HAdV-C5 and LH3 in LAdV-2. When atadenoviruses were first characterized, LH3 was considered a homolog for the HAdV E1B 55K protein, because of its location at the left end of the genome and a limited similarity with mastadenovirus E1B 55K sequences (Vrati *et al.*, 1996). However, the discovery that LH3 was present in the virion, unlike E1B 55K which is a multifunctional protein involved in cell control and transformation, shed doubts on the homology (Gorman *et al.*, 2005; Hidalgo *et al.*, 2019). Alignment of the LAdV-2 LH3 and HAdV-C5 E1B 55K sequences shows a modest similarity (11% identity). E1B 55K is longer than LH3 (496 *vs*. 370 aminoacids); the similarity starts at E1B 55K residue 145 (**Fig. S4D**), and the BETA-WRAP server (Bradley *et al.*, 2001) predicts a β-helix fold for E1B 55K starting at residue 135 (score -18.22). The E1B 55K gene (nt 2019 to 3509) is followed by the IX gene (nt 3609 to 4031) in the HAdV-C5 genome (NCBI Reference Sequence: AC_000008.1). That is, in HAdV-C5 the triskelion region is located downstream from the putative beta-helix fold, while in LAdV-2 LH3 the positions are reversed, with the triskelion at the N-terminus of the protein (**Fig. 4G**). This rearrangement would suggest a swapping of gene parts. However, it is also possible that this apparent swapping is the result of gene duplication.

Sequence alignment of a 40 residue fragment of E1B 55K starting at residue 145 (where the low homology with LH3 starts) with the triskelion sequences of LH3 and IX shows that a “triskelion-like” YX_8_W motif is also present in E1B 55K (**Fig. 4H**). However, the second hydrophobic residue in the triskelion core is not present, and proline groups are less abundant in the region that in LH3 and IX. Therefore, it is possible that the mastadenovirus E1B 55K and IX genes arose from a duplication of an LH3-like gene. Addition of the large N-terminal extension, together with loss of a hydrophobic residue would have abrogated the ability to form a triskelion and binding to the capsid shell in E1B 55K, resulting in its acquiring a whole new, nonstructural functionality. Meanwhile, the second copy maintained the triskelion, while incorporating large genetic changes that would give rise to the α-helical domains in mastadenovirus IX proteins, possibly conferring new properties to the viral particle (e.g. tropism).

In our previous report on the structure of SnAdV-1, we fitted the crystal structure of recombinant LH3 (lacking the N-terminal domain) into a low resolution cryo-EM map of the viral particle, and estimated the LH3 regions interacting with hexons using a hexon homology model (Menéndez-Conejero *et al.*, 2017). This report indicated that each LH3 monomer could interact with three hexon monomers located in two different trimers. Now we have the high resolution structure of both the complete LH3 protein and the surrounding hexons, which provides a much clearer picture of the contacts between LH3 and its neighbours in their biological context. We now appreciate that the hexon-LH3 interaction network is even more complex than previously proposed (**Table S9, and Fig. S5A)**. On the one hand, the β-helix domain of each LH3 monomer interacts with the towers of four hexon monomers from two different trimers (**Fig. S5A, β-helix**: yellow LH3 monomer (chain S) interacts with monomers tan and blue in trimer 2, and monomers pink and tan in trimer 3). The Asp155-Ser162 loop which was mobile in the recombinant protein is now ordered by these interactions. On the other hand, the triskelion region interacts with the base of three hexon monomers from three different trimers (**Fig. S5A, triskelion**: yellow LH3 monomer (chain S) interacts with the blue monomer of the three surrounding hexons). That is, each LH3 monomer bridges the surrounding capsomers by contacts with six hexon monomers located in three different trimers. These extensive interactions between LH3 and hexon contrast with those established by protein IX in HAdV-C5, located at a more basal position between the hexons except for the C-terminal helix bundle (Liu et al., 2010). Together with the strong intramolecular interactions of both hexon and LH3 trimers (Menéndez-Conejero *et al.*, 2017; Rux and Burnett, 2000), this extensive interlacing probably accounts for much of the increased thermostability of the atadenovirus capsids (Menéndez-Conejero *et al.*, 2017). Hexon surface charge is predominantly negative, while the LH3 surface has alternating positive and negative regions, suggesting an electrostatic component in the capsid stabilizing interaction (**Fig. S5B**).

### Internal minor coat proteins: protein VIII

Polypeptide VIII is 50 residues longer in LAdV-2 than in HAdV-C5 (278 *vs.* 227 residues). Assuming the same sequence specificity for the mast- and atadenovirus maturation proteases, LAdV-2 protein VIII would be cleaved after residues 121 (LHGG-A), 172 (LRGG-S) and 203 (LQGS-G). That is, the central excised region is 82 residues long, 33 residues longer than in HAdV-C5, where it stretches from residues Gly110 to Arg159 (Mangel and San Martín, 2014).

There are two copies of protein VIII per AU, located on the inner capsid surface (**Fig. 1B**). In both of them we have been able to trace most of the chain for the N-terminal (residues 2-120) and C-terminal (207-273) fragments (**Table S1, Fig. 5A and S6A**), consistently with the predicted maturation cleavages. Similarly to HAdV-C5, in LAdV-2 the protein VIII fold can be described as forming three domains: body (residues 2-77 and 225-273), neck (78-89, and 207-224), and head (90-120) (**Fig. 5A**). However, the protein folds differ more than those of hexon and penton base (RMSD 7.75 Å for 173 C-alpha atoms). The largest differences are located at the neck, an extended, largely unstructured domain which in HAdV-C5 has a small, 2-stranded β-sheet. In LAdV-2 however, the only secondary structure element in the neck is a 3-turn α-helix. The neck domain also encompasses the gap left by the maturation cleavage, unlike in HAdV-C5 where the gap is in the head domain. These differences observed in the neck domain relate to differences in the interactions between protein VIII with IIIa, and to other elements of the AU which are present in LAdV-2 but not in HAdV-C5 (see below).

**Figure 5.**
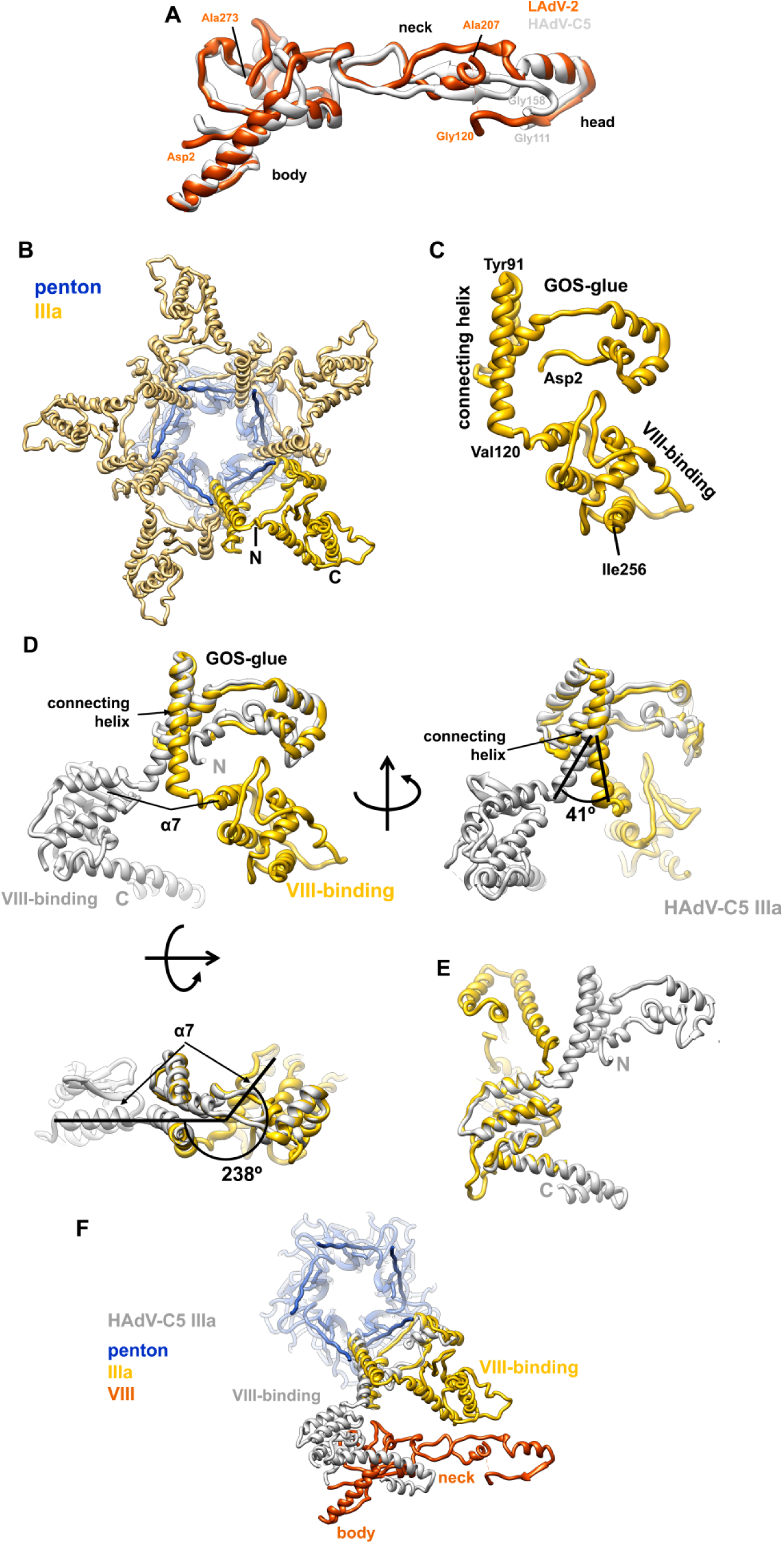
LAdV-2 internal minor coat proteins: VIII and IIIa. **(A)** Superposition of the LAdV-2 and HAdV-C5 protein VIII structures. The body, neck and head domains are indicated, as well as the positions of the N- and C-terminal residues in the structure and the residues flanking the central gap. **(B)** A view from inside the capsid along a 5-fold axis showing the ring of protein IIIa. One copy of IIIa is highlighted in vivid yellow, and the positions of the **N**- and **C**-terminal residues in the model are indicated. **(C)** Structure of the IIIa monomer. The **GOS-glue** and **VIII-binding** domains, as well as the **connecting helix**, the first and last residues traced, and those flanking the connecting helix are indicated. **(D)** Comparison between the LAdV-2 (yellow) and HAdV-C5 (grey) protein IIIa structures, presented in their original position in the capsid, in which the GOS-glue domains and part of the connecting helix overlap. Three points of view are shown, to highlight the large conformational change swinging the VIII-binding domain in the LAdV-2 protein away from its position in the human virus. The angles between the **connecting helices** and between the first helix in the VIII-binding domain of each protein (**α7**) are indicated, as well as the **N-** and **C-**termini of the HAdV-C5 protein. **(E)** Superposition of the two VIII-binding domains showing the fold similarity. **(F)** The LAdV-2 vertex proteins are depicted together with HAdV-C5 protein IIIa, to show the effect of the large conformational change on its interactions with protein VIII.

One copy of protein VIII (chain P) is located beneath the central plate of the facet, around the icosahedral 3-fold axis, while the other one (chain O) joins the central plate to the proteins in the peripentonal region (**Fig. 1B, 3B**). Each copy of protein VIII interacts with four different hexon trimers, one of them located in a different facet from the other three, therefore stabilizing each facet and riveting it to the next across the icosahedron edges (**Tables S7-S8, Fig. 1B and 3B**). In the head domain, protein VIII residues 108-113 form a β-strand that interacts with one of the jelly rolls in the neighbouring hexon trimer via a β-sheet augmentation (**Tables S7-S8, Fig. 1B and S6B**). Another β-sheet augmentation established between VIII and IIIa in HAdV-C5 (Liu *et al.*, 2010) is absent in LAdV-2 (see below).

Residues Asp54-Arg55 in H1 and H4 may establish electrostatic interactions with a pair of consecutive charged amino acids (Asp100-Lys101) located at an α-helix in the head domain of VIII (**Fig. S6C-D** and **Tables S7-S8**). Only one half of this interaction would be conserved across genera, since structure-guided sequence alignment indicates that in HAdV-C5 the same position corresponds to Glu98-Val99, lacking the basic residue in the pair (**Fig. S6D-E**).

### Internal minor coat proteins: protein IIIa

Five monomers of protein IIIa (one per AU) form a ring beneath each vertex, bridging the penton with the peripentonal hexons (**Fig. 5B, 3B**). Polypeptide IIIa is longer in LAdV-2 than in HAdV-C5 (609 *vs* 585 residues), but, as in HAdV-C5, only the N-terminal half appears to be ordered (**Table S1**). Similarly to HAdV-C5, in LAdV-2 the protein IIIa fold is predominantly α-helical, and can be described as forming two globular domains connected by a long α-helix (residues 93-121) (**Fig. 5C and S7**). By analogy with the HAdV-C5 structure (Liu *et al.*, 2010), the N-terminal domain (residues 2-92) is termed GOS-glue (GOS=Group of Six, consisting of the penton base pentamer and five surrounding peripentonal hexons), and the C-terminal domain (122-256) is designed as VIII-binding domain. When separately considered, these domains are quite similar to the corresponding ones in HAdV-C5, with RMSD values of 4.99 Å for the GOS-glue (residues 7-106 in HAdV-C5, 87 C-alpha pairs), 2.85 Å for the connecting helix (107-134 in HAdV-C5, 28 C-alpha pairs), and 1.8 Å for the VIII-binding domain (residues 135-269 in HAdV-C5, 124 C-alpha pairs). However, when the complete proteins are compared, a large difference is evident (**Fig. 5D, E**). The GOS-glue domain and start of the connecting helix occupy the same position in the capsid as their counterparts in HAdV-C5. However, at about half their length, the connecting helices start to become apart and end up separated by a 41 degree angle. Additionally, the helix is half a turn shorter in LAdV-2, resulting in the VIII-binding domain swinging away from its position in the human virus by 238 degrees. This large conformational change produces considerable differences in the network of contacts beneath the vertex.

Similarly to HAdV-C5 (Liu *et al.*, 2010), in LAdV-2 the IIIa GOS-glue domain and connecting helix establish an extensive set of interactions that join each IIIa monomer to two different peripentonal hexons; up to three penton base monomers; and the neighbouring IIIa molecule (**Table S6**). In spite of having a very similar fold, the VIII-binding domain is in a completely different location in the capsid and therefore establishes a completely different set of interactions. Instead of interacting exclusively with the body domain of the peripentonal copy of protein VIII, in LAdV-2 the VIII-binding domain of protein IIIa reaches outward to interact with hexon (**Table S6**), and establishes no contacts with the VIII body but with the VIII neck domain (**Fig. 5F**). The IIIa-VIII contacts are much fewer in LAdV-2: UCSF Chimera *findclash* estimates 84 possible contacts in HAdV-C5, *vs* only 22 in LAdV-2.

### Additional internal elements: proteins VI and VII, and unassigned density

The pseudo-hexagonal base of the hexon trimer encloses a central cavity that opens towards the interior of the viral particle. In HAdV-C5, weak density observed in this cavity allowed tracing of three copies per AU of the N-terminal peptide of protein VI cleaved by the protease during maturation (pVI_N_, residues 5-33), and one copy of a cleaved segment of core protein VII (pVII_N2_, residues 14-24) (Dai *et al.*, 2017). We observe patches of density associated with all 12 hexon monomers in the AU. However, the density did not have enough landmarks to unequivocally distinguish between proteins VI and VII. Following the HAdV-C5 model, we propose that the density patches in the cavity correspond to LAdV-2 peptides pVI_N_ (residues 2-25) or pVII_N2_ (residues 14-21) depending on the length of the peptide fragment that can be fitted in unfragmented density (**Table S1, Fig. 1B and Fig. S8**). Weak, fragmented density indicates variable occupancy, in agreement with the recently proposed model where proteins VI and VII compete for the same hexon binding site during assembly (Hernando-Pérez *et al.*, 2020).

Finally, we observe two additional groups of density fragments on the inner capsid surface for which we have been unable to unequivocally assign sequence identity, that we have termed “unassigned1” (U1) and “unassigned2” (U2) (**Fig. 1B**). The U1 density can hold three peptides of lengths 17, 36 and 41 residues with predominantly α-helical structure (**Fig. 6A and S9A**). It is present at two independent positions in the AU, near the gap left by maturation cleavages in protein VIII, and forms a wedge inserted at the local 3-fold axes surrounded by hexons 1, 2 and 4 in the AU, and hexons 3, 4 and 3 in the neighbouring AU (**Fig. 6B**). These are the 3-fold axes in the facet that do not hold an LH3 triskelion on the outer surface (**Fig. 3B**).

**Fig. 6.**
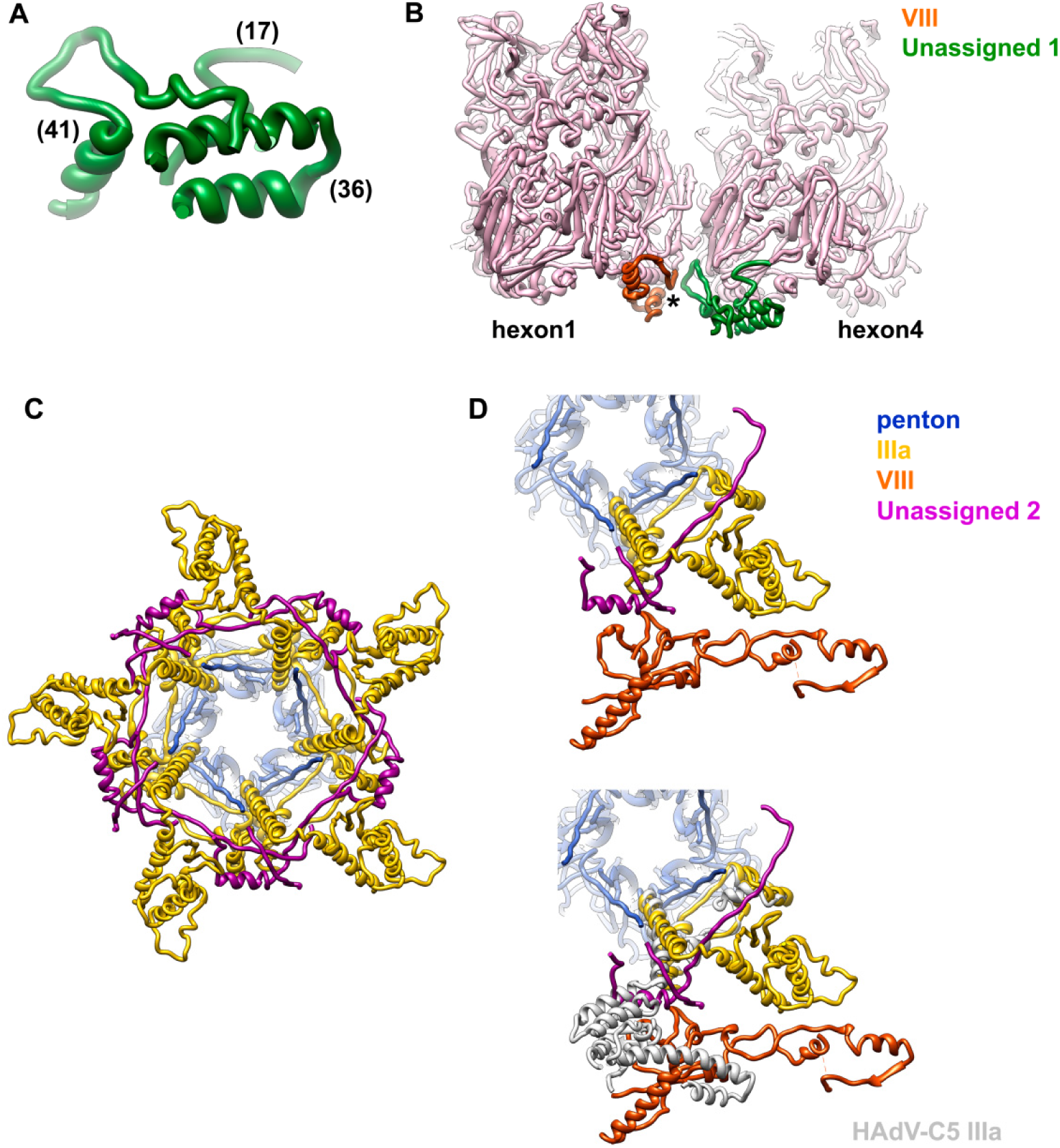
Unassigned elements on the LAdV-2 internal capsid surface. **(A)** Peptides modelled in the U1 density. Numbers in parentheses indicate the peptide lengths in amino acids. **(B)** View of the U1 peptides in the capsid context. The **star** indicates the location of the gap left by maturation cleavages in protein VIII. Hexon 2, which would be in front, has been removed for clarity. **(C)** Peptides modelled in the U2 density (magenta) form an interlaced ring with IIIa beneath the vertex. View along the 5-fold symmetry axis from inside the capsid. **(D)** The U2 peptides are intercalated between IIIa and the body domain of VIII (top). If IIIa adopted the same conformation as in HAdV-C5, it would badly clash with U2 (bottom).

The U2 density can hold two peptides of 50 and 16 amino acids, in a mostly extended conformation except for an α-helix (**Fig. S9B**). These peptides form an interlaced ring with protein IIIa beneath the vertex (**Fig. 6C-D**). A comparison with the structure of the vertex proteins in HAdV-C5 shows that the U2 peptides would clash with IIIa if this protein were not in a different conformation in the reptilian virus (**Fig. 6D**). U2 bridges IIIa with the body domain of VIII, compensating for the lost direct interactions. Although U2 has been modelled as a poly-alanine and therefore a proper interaction analysis cannot be carried out, it is notable that Chimera finds 24 possible contacts between U2 and proteins IIIa and VIII in the same AU, 52 with the neighbouring IIIa on one side and 11 more with the one on the other side (**Fig. S9C**).

## Discussion

AdVs have been found in most types of vertebrates, but few of them have been isolated and propagated in cell culture. Therefore, little is known about the basic biology of non-human AdV. We present here the first high resolution structure of an AdV infecting lower vertebrates, and not belonging to the mastadenovirus genus. The observation of the genus specific protein LH3 in the capsid context provides detailed insights on how this protein interlocks the surrounding hexons, contributing to make the atadenovirus capsid more stable than those of the HAdVs (Menéndez-Conejero *et al.*, 2017). Moreover, we show that the LH3 trimeric β-helix domain, folding as a bacteriophage tailspike, binds to the capsid surface using exactly the same structural motif (the triskelion) as its counterpart in mastadenoviruses, protein IX. The presence of a vestigial triskelion sequence in E1B 55kDa, a non-structural protein in HAdV-C5, suggests an ancient gene duplication. These observations indicate that the triskelion motif is critical for capsid binding, while the triskelion-carrying proteins are hotspots for AdV evolution, linked to capsid stabilization, interaction with host factors (because of their accessible location on the capsid surface) and consequently, tropism. Indeed, even within HAdVs, protein IX variations may be related to tropism determination, as recently shown by the structure of the enteric HAdV-F41 (Pérez-Illana *et al.*, 2020; Rafie *et al.*, 2020).

We show that even a protein critical for assembly and conserved throughout the AdV family, protein IIIa, can adopt very different conformations in two AdV genera. The large conformational change observed in IIIa is related to the presence of the extended, unidentified peptide U2 beneath the capsid vertex. As a result of the rotation of one of the IIIa domains, the interactions between IIIa and VIII are fewer than in the HAdVs, but U2 compensates this loss of contacts by reinforcing the interlacing of the internal vertex proteins. U2 in the vertex region, together with LH3 on the outer capsid surface, and U1 underpinning a second set of local 3-fold axes on the inside, appear as genus specific elements building a sturdier AdV capsid.

Possible virion components generating the U1 and U2 densities include maturation fragments or untraced regions of proteins IIIa, VI, or VIII (**Table S1**); as well as packaging proteins L1 52/55 kDa, IVa2, L4 33 kDa or L4 22 kDa (Condezo *et al.*, 2015; Guimet and Hearing, 2013; Gustin and Imperiale, 1998; Ostapchuk *et al.*, 2011; Wu *et al.*, 2013). Alternatively, U1 and U2 may correspond to the genus specific proteins LH2 and p32k. We favour this last possibility, since the other candidates are also present in mastadenoviruses, but no equivalent densities have been detected in the structures solved so far (Dai *et al.*, 2017; Liu *et al.*, 2010; Pérez-Illana *et al.*, 2020; Rafie *et al.*, 2020; Yu *et al.*, 2017). Both LH2 and p32k have positive charge (isoelectric point 11.02 for p32k, 8.59 for LH2) suggesting they interact with the genome, and are predicted to be predominantly α-helical. Protein p32k has also large regions predicted to be unstructured, similarly to the U2 peptides, and three consensus cleavage motifs for the maturation protease, again supporting an internal location (**Fig. S9D**) (Mangel and San Martín, 2014). Because of its proximity to the maturation gap in protein VIII, it is also possible that the excised peptides of VIII, which are 1.7 times longer in LAdV-2 than in HAdV-C5, form part of the U1 density (**Fig. 6B and S9D**). The two genus specific proteins and the longer central part of protein VIII would all be contributing to enhance the interactions stabilizing the atadenovirus capsid.

In summary, our work provides new information on the structure and evolution of AdVs, emphasizing the importance of minor coat proteins for determining specific physicochemical properties of the virions, and most likely their tropism. Additionally, knowing the structure of uncommon, non-human AdVs will facilitate their development as vectors, for example, by using the surface-exposed C-terminus of LH3 for display of peptides with biomedical interest (Matteson *et al.*, 2018).

## Methods

### Virus production

LAdV-2 (Pénzes *et al.*, 2014) was propagated in iguana heart epithelial cells (IgH-2, ATCC: CCL-108) (Clark *et al.*, 1970) at 37°C by amplification from one to 184 culture plates (10 cm diameter). The cells and supernatant from previous infections were used for the next infection round. Infected cells were frozen and thawed four times to release the viral particles before each infection step. Cell growth and virus propagation were carried out at 37°C. At 2 days post-infection in the last round, when the cells showed cytopathic effect, they were collected and viral particles purified by centrifugation in two consecutive CsCl gradients, as previously described (Condezo *et al.*, 2015).

### Cryo-electron microscopy

Purified LAdV-2 was dialyzed for 1 hour at 4°C against PBS (137 mM NaCl, 2.7 mM KCl, 10 mM Na_2_HPO_4_, 1.8 mM KH_2_PO_4_ pH 7.4) and concentrated by spinning in a Microcon YM-100 device for 6 min at 4°C, for a final estimated concentration of 7×10^12^ viral particles/ml. Samples were deposited in glow discharged, Quantifoil R2/4 300 mesh Cu/Rh grids and vitrified in liquid ethane after manual blotting in a Leica CPC device. Cryo-EM images (**Table S10**) were recorded using a 300 kV Titan Krios microscope equipped with a Falcon II detector (MRC-LMB, Cambridge, UK), with a total dose of 54 e-/Å^2^ distributed over 34 frames, at nominal pixel size 1.34 Å and defocus range between -1 and -3 µm.

### Image processing

All image processing and 3D reconstruction tasks were performed within the Scipion framework (**Table S10**) (de la Rosa-Trevin *et al.*, 2016). Frames 2-24 of each movie were aligned using whole-image motion correction implemented in Xmipp, followed by correction of local movements using Optical Flow (Abrishami *et al.*, 2015). The contrast transfer function (CTF) was estimated using CTFFIND4 (Rohou and Grigorieff, 2015). Particles were semi-automatically picked from micrographs corrected for the phase oscillations of the CTF (phase-flipped), extracted into 780×780 pixel boxes, normalized and downsampled by a factor of 2, using Xmipp (de la Rosa-Trevin *et al.*, 2013). All 2D and 3D classifications and refinements were performed using RELION (Scheres, 2012). 2D classification was used to discard low quality particles, and run for 25 iterations, with 50 classes, angular sampling 5 and regularization parameter T = 2. Classification in 3D was run for 40 iterations, with 3 classes, starting with an angular sampling of 3.7 degrees and sequentially decreasing to 0.5, and regularization parameter T = 4. Icosahedral symmetry was imposed throughout the refinement process. The initial reference for 3D classification was the SnAdV-1 cryo-EM map (Menéndez-Conejero *et al.*, 2017), low-pass filtered to 60 Å resolution. The class yielding the best resolution was individually refined using the original 780 px boxed particles and the map obtained during the 3D classification as a reference, producing a final map at 3.4 Å resolution, as estimated according to the gold-standard FSC = 0.143 criterion implemented in RELION auto-refine and postprocess routines (Chen *et al.*, 2013). A global B-factor was estimated after dividing the map Fourier transform by the modulation transfer function (MTF) of the Falcon II detector. The actual sampling for the map was estimated by comparison with the SnAdV-1 crystallographic and homology models (Menéndez-Conejero *et al.*, 2017) in UCSF Chimera (Pettersen *et al.*, 2004), yielding a value of 1.35 Å/px. Local resolution was calculated with ResMap (Kucukelbir *et al.*, 2014).

### Model building and analysis

The initial model for each polypeptide chain was predicted with Modeller (Webb and Sali, 2016), using as input template the structure of the respective homolog chain in HAdV-C5 (hexon: PDB ID 1P30; penton: PDB ID 1×9P; IIIa and VIII: PDB ID 3IYN) or SnAdV-1 (LH3: PDB ID 5G5O) (Liu *et al.*, 2010; Menéndez-Conejero *et al.*, 2017; Rux *et al.*, 2003; Zubieta *et al.*, 2005). UCSF Chimera (Pettersen *et al.*, 2004) was used to perform a rigid fitting of each chain initial model into the sharpened map. Next, the fitted model of each chain was refined using Coot (Emsley *et al.*, 2010), REFMAC (Brown *et al.*, 2015)and Phenix *real space refine* (Afonine *et al.*, 2018). Validation metrics to assess the quality of the atomic structure were computed with the Phenix *comprehensive validation (cryo-EM)* algorithm (**Table S10**). Once we generated the whole structure of the AU, the nearest neighbouring molecules were generated with Chimera (*sym #0 group i,222r contact 3*). Possible contacts between each molecule in the AU and all its neighbours were identified with Chimera *findclash*, listed and grouped using a protocol integrated in the Scipion molecular modelling workflow (Martinez *et al.*, 2020) (**Tables S2-S9**).

Sequence alignments were carried out with Clustal O 1.2.4 (Sievers *et al.*, 2011) and displayed as text or with JalView (Waterhouse *et al.*, 2009). Surface colouring by electrostatic potential was carried out with APBS and Chimera (Jurrus *et al.*, 2018; Pettersen *et al.*, 2004). Chimera *matchmaker* was used for RMSD calculation and structure guided alignment (followed by *match-align*). Secondary structure and disorder predictions were carried out with PsiPred and Disopred (Buchan and Jones, 2019; Jones, 1999; Jones and Cozzetto, 2015).

### Database deposition

The LAdV-2 cryoEM map and model are deposited at the Electron Microscopy Data Bank (EMDB, http://www.ebi.ac.uk/pdbe/emdb) and the Protein Data Bank (PDB, http://www.ebi.uk/pdbe) with accession numbers EMD-4551 and 6QI5, respectively.

## Acknowledgements

Work supported by the Spanish Agencia Estatal de Investigación and European Regional Development Fund (BFU2016-74868-P and PID2019-104098GB- I00/AEI/10.13039/501100011033); the Ministerio de Economía y Competitividad of Spain (BFU2013-41249-P and BIO2015-68990-REDT); and the Agencia Estatal CSIC (2019AEP045). The CNB-CSIC is further supported by a Severo Ochoa Excellence grant (SEV 2017-0712). We thank Shaoxia Chen, Sjors Scheres, Christos Savva, Greg McMullan (MRC-LMB, Cambridge, UK) and Felix de Haas (Thermo Fisher Scientific, formerly FEI, Eindhoven, NL) for cryo-EM data collection, as well as the INSTRUCT Image Processing Center (I2PC) for advice on data processing (INSTRUCT Access Project PID: 1395).

## Author contributions

G.N.C. prepared virus samples. G.N.C., R.M. and C.S.M. collected, processed and analysed cryoEM data. R.M., G.N.C., J.G.-B. and C.S.M. modelled, refined and analysed the structure. C.S.M. designed the study and wrote the paper with contributions from the rest of the authors.

## List of supplementary material

- Supplementary Tables S1 to S10
- Supplementary References
- Supplementary Figures S1 to S9
- Supplementary File S1

