## Supplementary material for "Near Atomic Structure of an Atadenovirus Reveals a Conserved Capsid-Binding Motif and Intergenera Variations in Cementing Proteins"

**Supplementary Tables**

**Table S1. Proteins in the LAdV-2 AU traced in this model.** Fragments that could not be unequivocally assigned to sequence are identified with the chain letter of the nearest identified molecule.

| Protein (UniProt ID) | Length (aminoacids) | Copy number in AU | Total aminoacids in AU | Chain ID | Residues traced |
|---|---|---|---|---|---|
| hexon (A0A076FYV7_9ADEN) | 909 | 12 | 10908 | A | 1-905 |
| | | | | B | 1-905 |
| | | | | C | 1-907 |
| | | | | D | 1-906 |
| | | | | E | 2-907 |
| | | | | F | 1-907 |
| | | | | G | 2-907 |
| | | | | H | 2-906 |
| | | | | I | 2-907 |
| | | | | J | 2-907 |
| | | | | K | 1-908 |
| | | | | L | 1-906 |
| penton base (A0A076FT28_9ADEN) | 451 | 1 | 451 | M | 1-451 |
| IIIa (A0A076FYV2_9ADEN) | 609 | 1 | 609 | N | 2-256 |
| VIII (A0A076FT36_9ADEN) | 278 | 2 | 556 | O | 2-120, 207-273 |
| | | | | P | 2-120, 207-273 |
| LH3 (A0A076FYU8_9ADEN) | 370 | 4 | 1480 | R | 4-370 |
| | | | | Q | 4-370 |
| | | | | S | 4-370 |
| | | | | T | 4-370 |
| VI (A0A076FTF8_9ADEN) | 226 | 9 | 2034 | near A | 2-25 |
| | | | | near B | 2-26 |
| | | | | near C | 2-25 |
| | | | | near D | 2-25 |
| | | | | near F | 2-25 |
| | | | | near H | 2-25 |
| | | | | near I | 2-25 |
| | | | | near K | 2-25 |
| | | | | near L | 2-25 |
| VII (A0A076FTB0_9ADEN) | 128 | 3 | 384 | near E | 14-21 |
| | | | | near G | 14-21 |
| | | | | near J | 14-21 |
| Unassigned densities. | Unassigned density beneath the central plate (U1) | 2 | ND | near O | fragments 36, 41, and 17 residues long |
| | | | | near P | fragments 36, 41, and 18 residues long |
| | Unassigned density beneath the vertex (U2) | 1 | ND | near N | fragments 50 and 16 residues long |

**Table S2.** Polypeptide regions involved in interactions between hexons in the same facet (ST interfaces)

| ST interactions (within the facet) | | | | | | | | | | | |
|---|---|---|---|---|---|---|---|---|---|---|---|
| | **H2** | | **H1** | | **H1** | | **H4** | | **H2** | | **H3** |
| E | Gln67-**Glu69**<br>Tyr74-Lys76 | A | **Arg62**-Ile66 | A | Glu69 | L | Thr558 | D | Gln67-**Glu69**<br>Tyr74-Lys76 | G | **Arg62**-Ile66 |
| | Arg78 | | Asn82 | | Arg78 | | Val64 | | | | |
| | Thr299-Gln301 | | Asn82-Arg87 | | Gln298-Leu302 | | Lys58-Ile66 | | Thr299-Leu302 | | Asn82-Asp85<br>Tyr571<br>Arg890 |
| | Leu306 | | Ser56 | | | | | | Leu306 | | Ser56 |
| | Asn71-Asn73 | C | Ala700-Glu701 | | | K | | | Asp70-Tyr74 | I | Glu701 |
| | Gly289-Ser290 | | Leu685-Trp689 | | Gly289-Ser290 | | Arg694<br>Gln698 | | Gly289-Ser290 | | Leu685-Asn688 |
| | | | | | Gln301-Leu302 | | Asp 682<br>Leu685 | | | | |
| | Asn309 | | Asn688 | | Leu306 | | Asn688 | | Asn309 | | Asn688 |
| | | | | | Tyr531 | | Ala700 | | | | |
| | Pro615-**Arg617** | | Asp616<br>**Asp673**-Ser675<br>Ser857 | | Pro615-Ser618 | | Val611-Asn613<br>Ser674 | | Pro615-**Arg617** | | Asp616<br>**Asp673**-Ser675<br>Ser857 |
| | Gly639 | | Trp689 | | | | | | Thr641 | | Asp699 |
| | Pro644 | | Gln698 | | | | | | | | |
| | Ala903-Ala907 | | Trp672-Val676<br>Asn848<br>His856 | | Ser904 | | Gln861 | | Phe902-Ser906 | | Ser675-Val676<br>**Met850** |
| | | | | B* | Glu2-Gln4 | | Ser675-Val676 | | | | |

(table continues in next page)

**Table S2 (continued)**

| ST interactions (within the facet) | | | | | | | | | | | | | |
|---|---|---|---|---|---|---|---|---|---|---|---|---|---|
| | **H4** | | **H2** | | | **H3** | | **H4** | | | **H3** | | **H3_AU4** |
| | Gln67-Glu69<br>Lys76 | | Val64-Ile66 | | | Gln67-Glu69<br>Lys76 | | Arg62-Ile66<br>Gln559 | | | Gln67-Glu69<br>Tyr74 | | Arg62-Ile66 |
| | | | | | | | | | | | Arg78 | | Asn82 |
| | Gln298-Leu302 | E | Ser56<br>Asn82-Arg87<br>Thr525<br>Tyr571<br>Arg890 | | | Thr299-Leu302 | J | Asn82-Arg87<br>Tyr571<br>Arg890 | | | Thr299-Leu302 | I | Asn82-Asp85<br>Tyr571<br>Arg890 |
| | Leu306 | | Ser56 | | | Leu306 | | Ser56 | | | Leu306 | | Ser56 |
| L | Asn73-Tyr74 | | Ala700-Glu701 | | I | Asn71-Tyr74 | | Ala700-Glu701 | | H | Asn71-Tyr74 | | Glu701 |
| | Gly289-Ser290 | | Leu685-Asn688 | | | Gly289-Ser290 | | Leu685-Asn688 | | | Gly289-Ser290 | | Leu685-Asn688 |
| | Asn309 | | Asn688 | | | Asn309 | | Asn688 | | | Asn309 | | Asn688 |
| | Pro615-Arg617 | D | Asp616<br>Asp673-Ser675<br>Ser857 | | | Pro615-Arg617 | L | Asp616<br>Asp673-Ser674<br>Ser857 | | | Pro615-Arg617 | H | Asp616<br>Asp673-Ser675<br>Ser857 |
| | Gly639 | | Trp689 | | | Gly639 | | Trp689 | | | Gly639<br>Thr641 | | Trp689<br>Asp699 |
| | Arg899-Ser906 | | Trp672-Val676<br>**Met850** | | | Arg899-Ala907 | | Ser675-Val676<br>**Asn848-Val849** | | | Ala903-Ser906 | | Ser675-Val676<br>**Met850** |

(table continues in next page)

**Table S2 (continued)**

| ST interactions (within the facet) | | | |
|---|---|---|---|
| **H4** | | **H3_AU2** | |
| J | Arg78 | H | Val64 |
| J | Thr299-Leu302 | H | Lys58<br>Arg62-Val64<br>Thr80-Asn82 |
| J | Ser290 | G | Gln698 |
| J | Leu306 | G | Asn688 |
| J | Tyr 531 | G | Ala700 |
| J | Asp616-Ser618 | G | Asn613<br>Ser674<br>Gln861 |
| J | Leu900-Gly905 | G | Ser675<br>Gln671-Ser674<br>Gln861 |
| K* | Gln4 | G | Ser675-Val676 |

Interacting amino acids were identified with UCSF Chimera *findclash* (Pettersen *et al.*, 2004). Following the nomenclature in Liu *et al.* (2010), **ST**, **TT** and **SS** indicate the three kinds of interfaces between hexons (**Fig. 3**). **S** refers to the sides of the hexon trimer pseudo-hexagonal base composed by the two β-barrels in a single monomer; **T** refers to the faces composed by two β-barrels coming from two different hexon monomers. **H1-H4** refer to the four hexon trimers in the icosahedral asymmetric unit; letters **A-L** denote the 12 hexon monomer chains in the icosahedral asymmetric unit. Suffixes _AU1 to _AU7 indicate neighbouring asymmetric units (**Fig. S3**). Red text and **blue text** highlight regions that include residues located at the hexon mobile regions 1 and 2, respectively (MR1 and MR2, **Fig. 2C**). **Purple text** indicates residues potentially involved in salt bridges. Cells shaded in light orange indicate N- and C-terminal flexible regions. Interacting amino acids are grouped by regions for clarity sake.

*Notice that some "S" interfaces, which are defined as involving a single hexon monomer on the basis of the hexagonal shape of the trimer, in fact may involve residues from two different monomers. This is due to the extensive interlacing of molecules in the hexon trimer, which results in the N-terminus of one hexon monomer reaching all the way to the center of the hexagon facet formed by the adjacent monomer (**Fig. 2A, 3A**).

**Table S3.** Polypeptide regions involved in interactions between hexons in different facets (TT interfaces)

| TT interactions (between facets) | | | | | | | | | | | |
|---|---|---|---|---|---|---|---|---|---|---|---|
| | | | | | | | | Icosahedral 2-fold axis | | | |
| | H1 | | H1_AU7 | | H4 | | H2_AU6 | | H2 | | H2_AU3 |
| C | Asp54-Arg55 | B | Asp54-Arg55-Ser56† | K | Asp54-Arg55-Ser56 | F | Asp54-Arg55† | D | Leu61-Ile66 | F | Gln671-Ser675<br>Asn859 |
| C | Arg55-Ser56 | A | Gly680-Asn681 | K | Ser56 | E | Asn681 | F | Gln671-Ser675<br>Asn859 | D | Leu61-Ile66 |
| C | Leu61 | A | Ser675 | K | Leu61-Val64 | E | Ser674-Ser675 | F | Thr609 | F | Ala700 |
| C | Asn82-Asp85-Arg87 | A | Asp673-Val676<br>Ser854-His856 | K | Asn82 | E | Asp673<br>His856 | F | Leu685-Asn688 | F | Leu685-Asn688 |
| C | | A | | K | Gln298 | E | Asp616 | F | Ala700 | F | Thr609<br>Gln861 |
| C | Arg890 | A | Ser855 | K | | E | | F | Gln861 | F | Ala700 |
| B | Asn681-Asp682 | A | Asp682<br>Asn688 | J | Asp682 | E | Gly680-Asn681 | | | | |
| B | Asp616 | B | Gln298 | J | | F | | | | | |
| B | Asp673-Val676 | B | Leu61-Ile66<br>Asn82 | J | Asp673-Val676 | F | Asn82-Arg87 | | | | |
| B | Asn681 | B | Ser56 | J | Asn681 | F | Arg55-Ser56 | | | | |
| B | Tyr851-His856 | B | Asn82-Asp85<br>Thr299<br>Thr525<br>Glu578<br>Arg890-Ser891 | J | Met850<br>His856 | F | Asp85-Asn86 | | | | |

Nomenclature and color codes as in the previous table.

†A possible double salt bridge ($Asp54_C$-$Arg55_B$/$Asp54_B$-$Arg55_C$) is observed at the TT interfaces at the local, but not at the icosahedral, 2-fold axes.

**Table S4.** Polypeptide regions involved in interactions between hexons in different facets (SS interfaces)

| SS interactions (between facets) | | | | | | | |
|---|---|---|---|---|---|---|---|
| | H4 | | H1_AU6 | | H3 | | H2_AU3 |
| K | Gln67 | C | Pro615 | G | Ile66-Gln67 | F | Asn613-Pro615 |
| | Asn73 | | Gly639 | | Tyr74 | | Asn309 |
| | Arg78 | | Pro615 | | | | |
| | Ser290 | | Leu308 | | Ser290 | | Asp307-Leu308 |
| | Thr292 | | Gly905 | | Thr292 | | Gly905 |
| | Gln298-Gln301 | | Asp616-Ser618<br>Asn853-Ser855 | | Gln298-Gln301 | | Asp616-Ser618<br>Asn853-Ser855 |
| | | | | | Leu306-Leu308 | | Ser906-Ala907 |
| | Asn309-Asp310 | | Gln301-Leu302 | | Asp310 | | Leu302 |
| | Asn599-Leu600 | | Gln301 | | | | |
| | Ile614-Arg617 | | Thr299-Gln300<br>Thr889 | | Pro615-Arg617 | | Gln298-Gln300<br>Thr889 |
| | Ala640-Gln642 | | Gly289-Ser290 | | Ala640-Gln642 | | Gly289-Ser290 |
| | Pro644-Asn645 | | Glu69<br>Tyr74 | | Pro644-Asn645 | | Glu69<br>Tyr74 |
| | | | | | Phe902 | | Gln300 |
| | Gly905-Ser906 | | Asn886<br>Ser894-Ala896 | | Ser904 | | Ser894 |
| | Ala908 | | Ala584 | | Ala907 | | Ala584 |
| | Gln301-Leu302 | A* | Met1 | | Gln300-Leu302 | D* | Met1-Arg5 |
| | Ser906-Ala907 | | Phe7 | | Ser906-Ala907 | | Phe7 |

Nomenclature and color codes as in the previous tables.

*Notice that some "S" interfaces, which are defined as involving a single hexon monomer on the basis of the hexagonal shape of the trimer, in fact may involve residues from two different monomers. This is due to the extensive interlacing of molecules in the hexon trimer, which results in the N-terminus of one hexon monomer reaching all the way to the center of the hexagon facet formed by the adjacent monomer (**Fig. 2A, 3A**).

**Table S5.** Polypeptide regions involved in interactions between hexon and penton base

| SP interaction | | | | | | | | |
|---|---|---|---|---|---|---|---|---|
| H1 | | P | | | H1 | | P_AU6 | |
| B | Glu69 | M | Asp335 | | B | Thr299-Gln301 | M | Pro24-Ala26 |
| | Arg78 | | Asn332 | | | | | |
| | Gly289-Ser290 | | Leu47<br>Lys51<br>Tyr57 | | | | | |
| | Ser296<br>Gln301 | | Asn331-Asn332 | | | | | |
| | Leu308 | | Thr73 | | | | | |
| | Asn309 | | Leu47 | | | | | |
| | Asn527-Tyr531 | | Asn332-Thr334 | | | | | |
| | Pro615-**Arg617** | | Asp65-Val66<br>**Asp70**<br>Gln74 | | | | | |
| | **Arg673**-Pro644 | | **Glu45**-Ile49 | | | | | |
| | Arg899-Gly905 | | Thr73-Thr75 | | | | | |
| C* | Met1 | | Arg81 | | | | | |

Nomenclature and color codes as in the previous tables. **SP** denotes the hexon-penton interface; **P** indicates the penton base monomer (with chain id M in the coordinate file).

*Notice that some "S" interfaces, which are defined as involving a single hexon monomer on the basis of the hexagonal shape of the trimer, in fact may involve residues from two different monomers. This is due to the extensive interlacing of molecules in the hexon trimer, which results in the N-terminus of one hexon monomer reaching all the way to the center of the hexagon facet formed by the adjacent monomer.

**Table S6.** Polypeptide regions involved in interactions between IIIa and other capsid components

| IIIa interactions with hexon | | | | | | | |
|---|---|---|---|---|---|---|---|
| **IIIa** | | **H1** | | **IIIa** | | **H1_AU6** | |
| *Domain* | *Amino acids* | *Amino acids* | *Chain* | *Domain* | *Amino acids* | *Amino acids* | *Chain* |
| GOS-glue | Pro3-Pro7 | Leu579-Thr585 | B | GOS-glue | Gly78-Asp83 | **Met850-Ala852** | B |
| | Leu36-Leu40 | **Leu842-Met850** | | | | | |
| | Lys52-Gln64 | Leu579-Thr585 | | | | | |
| | Pro74-Ile86 | Arg55<br>Asn86<br>Tyr571-Ser575<br>Glu578-Leu579<br>Thr889-Arg890 | | | | | |
| VIII-binding | Val139-Gly143 | Glu17-Gln28 | | | | | |
| GOS-glue | Pro3-Thr12 | Arg13-Asn22 | C | | | | |
| | Thr12 | Glu6 | | | | | |
| | Pro35-Asn38 | Val45-Pro47<br>Ala11-Gly12 | | | | | |
| | Thr41-Glu44 | Pro3-Arg5 | | | | | |
| | Pro50 | Met1 | | | | | |

(table continues in next page)

**Table S6 (continued)**

| IIIa interactions with penton | | | | | | | | | | | |
|---|---|---|---|---|---|---|---|---|---|---|---|
| **IIIa** | | **P** | | **IIIa** | | **P_AU6** | | **IIIa** | | **P_AU5** | |
| *Domain* | *Amino acids* | *Amino acids* | *Domain* | *Domain* | *Amino acids* | *Amino acids* | *Domain* | *Domain* | *Amino acids* | *Amino acids* | *Domain* |
| Conn. helix | Asn92-Thr94 | Gln451 | C-terminus | GOS-glue | Asp17-Ala21 | Met1-Val3 | N-terminal arm | Conn. helix | Asn95-Val102 | Glu2-Pro7 | N-terminal arm |
| | | | | | Gln49-Asp53 | Tyr4-Arg9 | | | Asp106 Ala109 | Met1 | |
| | | | | | His89-Asn92 | Val10-Glu15 | | | | | |

| IIIa interactions with other IIIa molecules | | | |
|---|---|---|---|
| **IIIa** | | **IIIa_AU6** | |
| *Domain* | *Amino acids* | *Amino acids* | *Domain* |
| GOS-glue | **Glu65**-Met69 | Ile16-**Arg23** | GOS-glue |
| | Met69-Ala71 | Val28-Arg32 | |
| | **Glu76**-**Arg87**-Val88 | **Lys39**-**Glu44**-**Asp45**-Val48 | |
| Connecting helix | Val96-Leu103 | Ala46-Gln49 | |
| | Asp106-Ile107 | Ile20-Ala21 | |

| IIIa interactions with protein VIII | | | |
|---|---|---|---|
| **IIIa** | | **VIII (chain O)** | |
| *Domain* | *Amino acids* | *Amino acids* | *Domain* |
| VIII-binding | Val139-Gln144 | Arg221-Ile222 | Neck |
| | Thr236-Asp238 | Tyr85-Pro87 | |
| | **Asp238** | **Lys211** | |

Nomenclature and color codes as in the previous tables. Notice that, because of our choice of AU convention and the torsion of penton base monomers around the 5-fold axis, the most extensive contacts between IIIa and penton turn out to be with the penton monomer belonging to the next AU.

**Table S7.** Polypeptide regions involved in interactions between protein VIII (chain O) and hexons in the peripentonal region. For interactions between VIII and IIIa, see **Table S6**.

| **Interactions for protein VIII, chain O (peripentonal region)** | | | | | | | |
|---|---|---|---|---|---|---|---|
| **VIII_O** | | **H1** | | **VIII_O** | | **H2** | |
| *Domain* | *Amino acids* | *Amino acids* | *Chain* | *Domain* | *Amino acids* | *Amino acids* | *Chain* |
| Body | Gln13<br>Ala20 | Asn853 | A | Head | Ser104-Glu108 | Leu898-Gly905 | E |
| | Gln15 | Ser618 | | | Glu108-Ser113 | Leu302-Leu308 | |
| Head | Pro89 | Glu576 | | | | Ser894-Leu898 (*β-strand sheet augmentation*) | |
| | Leu92 | Ala49 | | | Arg114-Pro115 | Pro888-Gly892 | |
| | Gly94-Thr97 | Asn570-Asp572 | | | Glu106-Glu108 | Gln4-Phe8 | F |
| | Arg99-**Asp100**-**Lys101**-Ser104 | **Asp54**-**Arg55** | | | | | |
| | Leu110-Gly112 | Arg55-Ser56 | | | | | |
| | Ser113 | Tyr571 | | | | | |
| | Leu118-***Gly120*** | Asp85<br>Glu578-Leu579<br>Arg890 | | | | | |
| Neck | Val212-Pro220 | Leu579-Thr585 | | | | | |
| Body | Leu260-Phe261 | **Val849-Met850** | | | | | |
| | Pro14-Gly17 | Pro3-Arg5 | B | | | | |
| Neck | Tyr85-Ala88 | Glu21-Gln25 | | | | | |
| | Leu215-**Arg221**-Pro223 | Arg13-**Glu17**-**Glu21**-Val24 | | | | | |
| Body | Glu226-Pro230 | Glu2-Glu6 | | | | | |
| Head | Ala105 | **Tyr851** | C | | | | |

(table continues in next page)

**Table S7 (continued)**

| Interactions for protein VIII, chain O (peripentonal region) | | | | | | | | |
|---|---|---|---|---|---|---|---|---|
| **VIII_O** | | **H4** | | | **VIII_O** | | **H1_AU6** | |
| *Domain* | *Amino acids* | *Amino acids* | *Chain* | | *Domain* | *Amino acids* | *Amino acids* | *Chain* |
| Body | Ile9-Gln11 | Asp616<br>Ala852-Ser855 | K | | Body | Trp32-Phe40 | Met1-Gln4<br>Phe7-Phe8 | A |
| | Gln11-Gln15 | Asp673-Val676 | | | | Pro245-Gln248 | Ser20-Asn22 | |
| | Asn25-Tyr26 | Asp616-Ser618 | | | | Tyr8 | Asn583 | C |
| | Gly27-Met39 | Ser894-Ala907 | | | | Arg23-Tyr26 | Leu579<br>Arg582-Ala584<br>Pro888-Thr889 | |
| | Asn31-Ser34 | Leu302-Leu306 | | | | Leu33 | Ala896 | |
| | Asn38-Met39 | Val884-Pro888 | | | | Pro245-Phe246<br>Phe253-Phe257 | Glu576-Thr585 | |
| | **Arg42** | Ala584<br>**Asp587** | | | | Leu260 | Asn86 | |
| | Val43 | Ala896 | | | | | | |
| | Pro230 | **Met850** | | | | | | |
| | Pro239 | Asn853 | | | | | | |
| | Ala28 | Gln4 | L | | | | | |
| | Ile30 | Phe8 | | | | | | |
| | Met39<br>Arg42 | Phe7 | | | | | | |
| | Ala49-Arg50<br>Ile53 | Pro3 | | | | | | |

Nomenclature and color codes as in the previous tables. Interacting amino acids are grouped by regions for clarity sake, but interesting residues (e.g. those in **purple**) may be explicitly named in the middle of some regions.

**Table S8.** Polypeptide regions involved in interactions between protein VIII (chain P) and hexons in the central plate region.

**Interactions for protein VIII, chain P (central plate region)**

| VIII_P | | H4 | |
|---|---|---|---|
| *Domain* | *Amino acids* | *Amino acids* | *Chain* |
| Body | Gln13<br>Ala20 | Asn853 | J |
| Head | Pro89 | Glu576 | J |
| Head | Leu92 | Ala49 | J |
| Head | Gly94-Thr97 | Asn570-Asp572 | J |
| Head | Arg99-**Asp100**-**Lys101**-Ser104 | **Asp54**-**Arg55** | J |
| Head | Leu110, Gly112 | Arg55-Ser56 | J |
| Head | Leu118-***Gly120*** | Asp85<br>Glu578-Leu579<br>Arg890 | J |
| Neck | Leu213-Pro220 | Leu579-Thr585 | J |
| Body | Leu260-Glu262 | **Val849-Met850**<br>Ser854 | J |
| Body | Pro14-Gly17 | Pro3-Arg5 | K |
| Neck | Tyr85-Ala88 | Glu21-Gln25 | K |
| Neck | Leu215-**Arg221**-Pro223 | Arg13-**Glu17**-**Glu21**-Val24 | K |
| Neck | **Arg224** | **Glu6** | K |
| Body | Gly227-Pro230 | Glu2-Pro3 | K |
| Head | Ala105 | **Tyr851** | L |

| VIII_P | | H3 | |
|---|---|---|---|
| *Domain* | *Amino acids* | *Amino acids* | *Chain* |
| Head | Ser104-Glu108 | Leu898-Ser906 | I |
| Head | Glu108-Ser113 | Gln300-Leu306 | I |
| Head | Glu108-Ser113 | Ile893-Leu898<br>(*β-strand sheet augmentation*) | I |
| Head | Arg114-Pro115 | Pro888-Gly892 | I |
| Head | Glu106-Glu108 | Gln4-Phe8 | G |

(table continues in next page)

**Table S8 (continued)**

**Interactions for protein VIII, chain P (central plate region)**

| VIII_P | | H3_AU2 | |
|---|---|---|---|
| *Domain* | *Amino acids* | *Amino acids* | *Chain* |
| Body | Thr6 | Ser906 | G |
| | Ile9-Gln11 | Asp616-Ser618<br>Asn853-Ser855 | |
| | Pro14-Gln15 | Asp673-Val676 | |
| | Asn25-Gly27 | Asp616-Ser618 | |
| | Gly27-Met39 | Ser894-Ser906 | |
| | Asn31-Ala35 | Leu302-Leu306 | |
| | Asn38-Met39 | Val884-Pro888 | |
| | **Arg42** | Ala584<br>**Asp587** | |
| | Val43 | Ala896 | |
| | Pro230 | **Met850** | |
| | Pro239 | Asn853 | |
| | Ile30 | Gln4<br>Phe8 | H |
| | Met39<br>Arg42<br>Val46 | Phe7 | |
| | Ala49-Arg50 | Pro3 | |
| | Ile53 | Glu2-Pro3 | |

| VIII_P | | H2_AU6 | |
|---|---|---|---|
| *Domain* | *Amino acids* | *Amino acids* | *Chain* |
| Body | Trp32-Phe40 | Glu2-Gln4<br>Phe7-Phe8 | D |
| | Pro245-Gln248 | Ser20-Asn22 | |
| | Tyr8 | Asn583 | F |
| | Arg23-Tyr26 | Leu579<br>Arg582<br>Pro888-Thr889 | |
| | Leu33 | Ala896 | |
| | Pro245-Phe246<br>Phe253-Phe257 | Glu576-Thr585 | |
| | Leu260 | Asn86 | |

Nomenclature and color codes as in the previous tables.

**Table S9.** Polypeptide regions involved in interactions between protein LH3 and hexons.

**At the local 3-fold symmetry axis**

| LH3 | | | H3 | |
|---|---|---|---|---|
| *Domain* | | *Amino acids* | *Amino acids* | |
| Triskelion | Q | Glu5-Ile10 | Thr609-Asn613<br>Gln861 | I |
| | | Ile10-Ile13 | Asn599-Tyr601 | |
| | | Ile13-Ala18 | Asn309<br>Arg637-Ala640<br>Pro644-Asn645 | |
| β-helix | | Gln54 | Gln771 | |
| | | His56-Lys58 | Val259<br>Ala437-Asn438 | |
| | | Ser78 | Val259 | |
| | | Thr80 | Val132 | |
| | | **Glu106** | **Arg236** | |
| | | Thr109-**Arg112**-Met113 | Tyr146-Ala147<br>**Glu155**<br>Gln159<br>Asn185 | G |
| | | Leu259 | Thr154-Glu155<br>Gln158 | |

| LH3 | | | H4 | |
|---|---|---|---|---|
| *Domain* | | *Amino acids* | *Amino acids* | |
| Triskelion | Q | Asn11 | Val611 | L |
| | | Asn14-Pro17 | Arg667-Gln671<br>Trp689<br>Gln861-Thr863 | |
| β-helix | | Arg90-Thr91 | Gln160-Ser162<br>Asp165 | J |
| | | **Asn157-Phe160** | Ile157-Gln160 | |
| | | | Glu384-Gln387 | K |

| LH3 | | | H2 | |
|---|---|---|---|---|
| *Domain* | | *Amino acids* | *Amino acids* | |
| Triskelion | Q | None detected | None detected | D |

(table continues in next page)

**Table S9 (continued)**

**At the local 3-fold symmetry axis**

| **LH3** | | | **H4** | |
|---|---|---|---|---|
| *Domain* | | *Amino acids* | *Amino acids* | |
| Triskelion | R | Glu5-Ile10 | Thr609-Val611 | L |
| | | Ile10-Gln15 | Asn599-Tyr601 Asn309 | |
| | | Ile13-Ala18 | Arg637-Ala640 Asn645 | |
| β-helix | | Gln54 | Gln771 | |
| | | His56-Lys58 | Ala437-Asn438 | |
| | | Gly79 | Val259 Arg263 | |
| | | Glu118 | Asn134 | |
| | | Thr109-**Arg112** | Tyr146-Ala147 **Glu155** Gln159 | J |
| | | Leu259 | Thr154-Glu155 Gln158 | |

| **LH3** | | | **H2** | |
|---|---|---|---|---|
| *Domain* | | *Amino acids* | *Amino acids* | |
| Triskelion | R | Asn11 | Val611-Asn613 | D |
| | | Pro12-Asn14 | Gln861-Thr863 | |
| | | Asn14-Pro17 | Arg667-Ser669 Trp689 | |
| | | Pro19 | Ser608 | |
| β-helix | | Arg90-**Lys93** | Gln160-**Asp165** | E |
| | | **Asn157-Phe160** | Ile157-Gln158 | |
| | | | Glu384-Gln387 | F |

| **LH3** | | | **H3** | |
|---|---|---|---|---|
| *Domain* | | *Amino acids* | *Amino acids* | |
| Triskelion | R | Glu6-Leu7 | Gln602 | I |

(table continues in next page)

**Table S9 (continued)**

At the local 3-fold symmetry axis

| LH3 | | | H2 | |
|---|---|---|---|---|
| *Domain* | | *Amino acids* | *Amino acids* | |
| Triskelion | S | Glu5-Ile10 | Thr609-Asn613 Gln861 | D |
| | | Ile10-Ile13 | Asn599-Tyr601 | |
| | | Ile13-Trp16 | Arg637-Ile638 Asn309 | |
| | | Pro17-Ala18 | Ala640 Pro644-Asn645 | |
| β-helix | | Gln54 | Gln771 | |
| | | His56-Lys58 | Ala437-Asn438 | |
| | | Ser78-Gly79 | Val259 | |
| | | Thr80 | Val132 | |
| | | Glu118 | Asn134 | |
| | | Thr109-Asn111 | Tyr146-Ala147 | E |
| | | **Arg112** | **Glu155** | |
| | | Met113 | Asn185 | |
| | | Leu259 | Thr154-Glu155 Gln158 | |

| LH3 | | | H3 | |
|---|---|---|---|---|
| *Domain* | | *Amino acids* | *Amino acids* | |
| Triskelion | S | Asn11 | Val611 | I |
| | | Asn14-Pro19 | Arg667-Ser669 Gln861-Thr863 Trp689 | |
| β-helix | | Arg90-Thr91 | Gln160-Ser162 | G |
| | | **Asn157-Phe160** | Ile157-Gln160 | |
| | | | Glu385-Gln387 | H |

| LH3 | | | H4 | |
|---|---|---|---|---|
| *Domain* | | *Amino acids* | *Amino acids* | |
| Triskelion | S | **Glu5** | **Arg637** | L |
| | | Glu6-Leu7 | Gln602 | |

(table continues in next page)

**Table S9 (continued)**

**At the icosahedral 3-fold symmetry axis**

| LH3 | | | H3 | |
|---|---|---|---|---|
| *Domain* | | *Amino acids* | *Amino acids* | |
| Triskelion | T | Leu7-Val9 | Val611 Gln861 | H |
| | | Ile10-Ile13 | Leu600-Gln602 Tyr872 | |
| | | Ile13-Ala18 | Arg637-Ile638 Ala640 Asn645 | |
| β-helix | | Gln54 | Gln771 | |
| | | His56-Lys58 | Ala437-Asn438 | |
| | | Ser78-Gly79 | Val259 | |
| | | Thr80 | Val132 Asp260 | |
| | | Thr109- Arg112- Met113 | Tyr146-Ala148 Glu155 | I |
| | | Leu259 | Thr154-Glu155 Gln158 | |

| LH3 | | | H3_AU4 | |
|---|---|---|---|---|
| *Domain* | | *Amino acids* | *Amino acids* | |
| Triskelion | T | Asn11 | Val611 | H |
| | | Pro12-Trp16 | Gln861-Thr863 | |
| | | Asn14-Pro17 | Arg667-Ser669 Trp689 | |
| β-helix | | Arg90-Lys93 | Gln160-Asp165 | I |
| | | Asn157-Phe160 | Ile157-Gln158 | |
| | | | Glu384-Gln387 | G |

| LH3 | | | H3_AU2 | |
|---|---|---|---|---|
| *Domain* | | *Amino acids* | *Amino acids* | |
| Triskelion | T | Glu5 | Arg637 | H |
| | | Glu5-Tyr8 | Gln602-Pro604 | |

**H1-H4** refer to the four hexon trimers in the icosahedral asymmetric unit; letters **A-L** denote the 12 hexon monomer chains, and **Q-T** the 4 LH3 chains in the AU. Suffixes _AU1 to _AU7 indicate neighboring asymmetric units (**Fig. S3**). **Purple text** indicates residues potentially involved in salt bridges. **Green text** refers to the surface loop disordered in the crystal structure but fixed in place by interactions with hexon in the capsid (Menéndez-Conejero *et al.*, 2017). Interacting amino acids are grouped by regions for clarity sake, but interesting residues (e.g. those in **purple**) may be explicitly named in the middle of some regions.

**Table S10.** Cryo-EM data collection, refinement, modeling and validation statistics.

| **Data collection** | |
|---|---|
| Microscope | Titan Krios |
| Camera | Falcon II |
| Voltage | 300 kV |
| Magnification | 59,000 |
| Nominal pixel size | 1.34 Å |
| Dose rate | 27 e/Å$^2$s |
| Cumulative electron dose | 54 e/Å$^2$ |
| Exposure time | 2 s |
| Number of frames | 34 |
| Defocus range | -1 to -3 μm |
| Micrographs collected | 2,618 |
| Acquisition software | EPU |
| **Image processing** | |
| Preprocessing software | XMIPP, Scipion |
| Frame alignment software | Xmipp optical flow , Scipion |
| CTF estimation software | CTFFIND4, Scipion |
| Particle picking software | XMIPP, Scipion |
| Micrographs used | 2,614 |
| Particles selected | 19,128 |
| **Reconstruction** | |
| Software | RELION, Scipion |
| Particles included | 16,071 |
| Symmetry imposed | Icosahedral I2 |
| Rotational accuracy | 0.05 degrees |
| Translational accuracy | 0.10 px |
| B-factor applied | -93.86 Å$^2$ |
| Final resolution (gold standard FSC=0.143) | 3.4 Å |
| **Model building, refinement and validation** | |
| Experimental pixel size | 1.35 Å |
| Number of modeled residues | 13,417 |
| Number of chains | 20 |
| Software | Coot, Phenix, REFMAC, UCSF Chimera |
| Model to map correlation coefficient | 0.826 |
| R.M.S. deviations (bond length) | 0.0069 Å |
| R.M.S. deviations (bond angle) | 1.29º |
| Ramachandran preferred | 91.13% |
| Ramachandran outliers | 0.21% |
| Rotamer outliers | 0.36% |
| C-beta outliers | 0 |
| Molprobity score | 1.83 |
| Clashscore | 5.81 |

**Supplementary Figures**

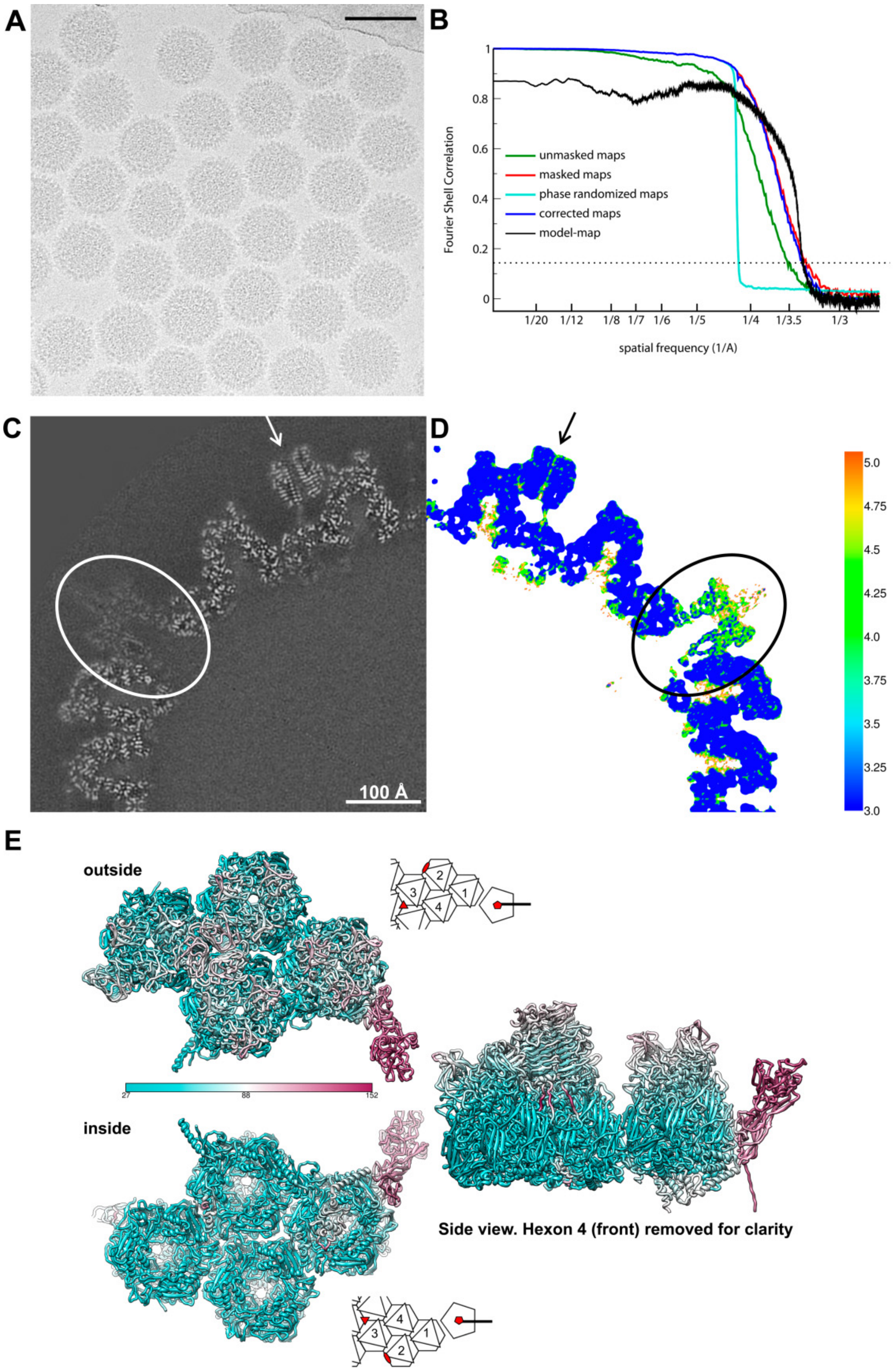


**Figure S1. LAdV-2 cryo-EM data and map resolution**. **(A)** Representative cryo-EM field. The bar represents 100 nm. **(B)** Resolution curves calculated with *relion postprocess* according to the gold standard procedures, and with Phenix *model-map*, as indicated. The FSC=0.143 threshold is indicated by a dotted line. **(C)** Detail of the cryo-EM map central slice. The penton region is indicated by a white oval and one LH3 trimer by a white arrow. Highest protein density is white. **(D)** The same central section coloured by local resolution (in Å) as calculated with *ResMap*. A black oval and arrow indicate penton base and LH3, as in (C). **(E)** The asymmetric unit is shown as seen from outside or inside the virion (compare with Fig. 1B), or in a side view as indicated. Colours represent the B-factor values provided by *Phenix* real space refinement, according to the scale bar (in $Å^2$).

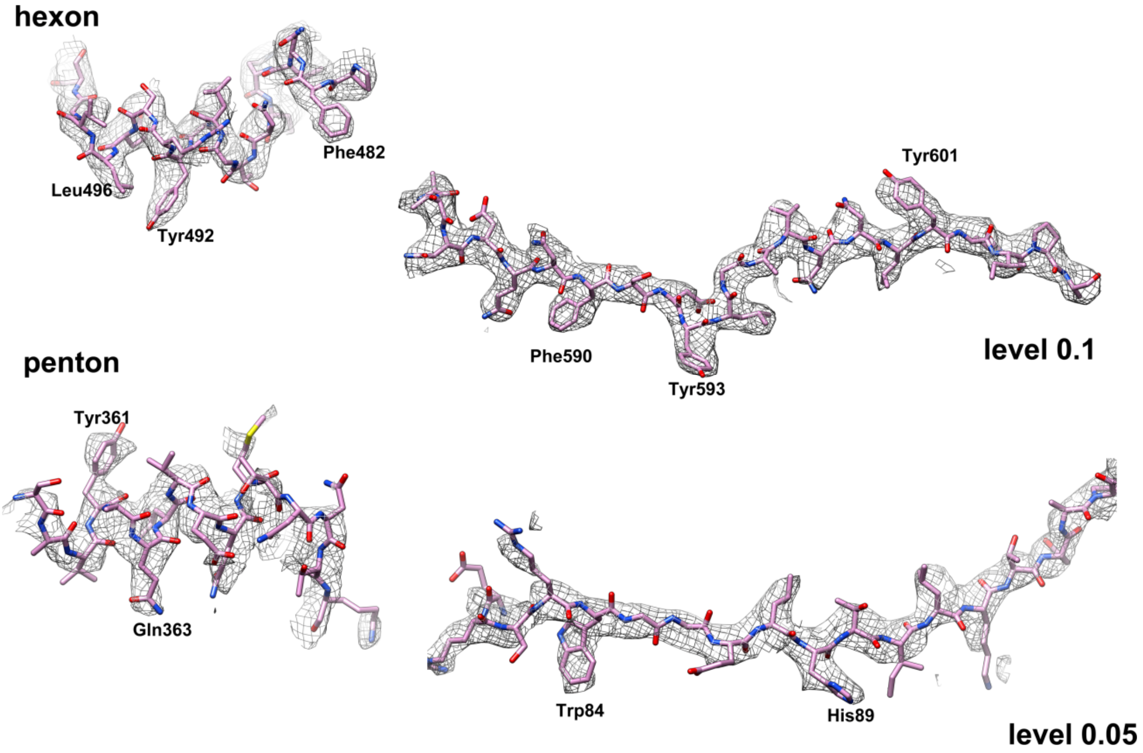


**Figure S2. LAdV-2 main capsid proteins.** Details of the molecular model fit on the density map for hexon and penton base. The map contour level for each protein is indicated at the right.

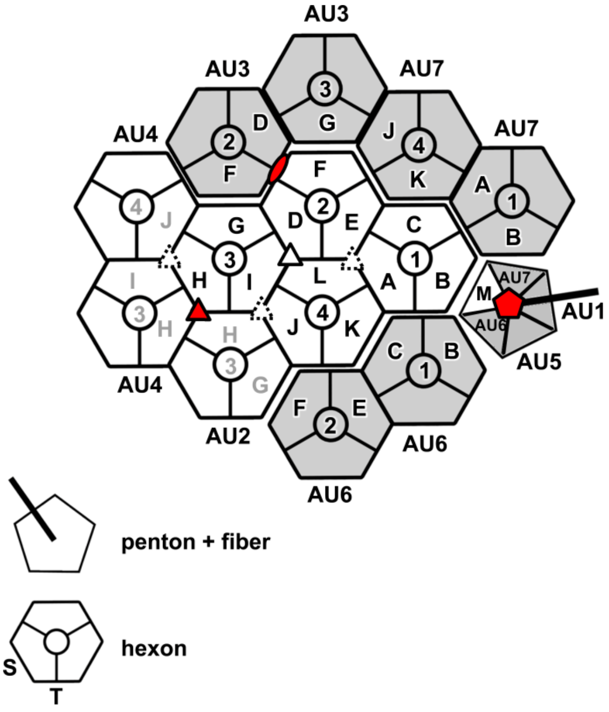


**Figure S3. Cartoon showing one AU and its immediate neighbours**. Hexons 1-4 in one AU are depicted in white and labelled with black text; those in neighbouring AUs in the same icosahedral facet are in white with grey labels; and those in adjacent facets are in grey. Letters **A-M** identify different polypeptide chains. Neighbouring AUs are numbered AU1-AU7 as in **Tables S2-S9.** In the hexon schematics legend, **S** designates the facet of the hexon pseudo-hexagonal base formed by the two β-barrels in a single monomer, and **T** indicates the facet of the hexon pseudo-hexagonal base formed by two β-barrels belonging to two adjacent hexon monomers (Liu *et al.*, 2010). Red symbols indicate the icosahedral 5-fold (pentagon), 3-fold (triangle) and 2-fold (oval) symmetry axes. Closed and broken white triangles indicate two different sets of local 3-fold symmetry axes.

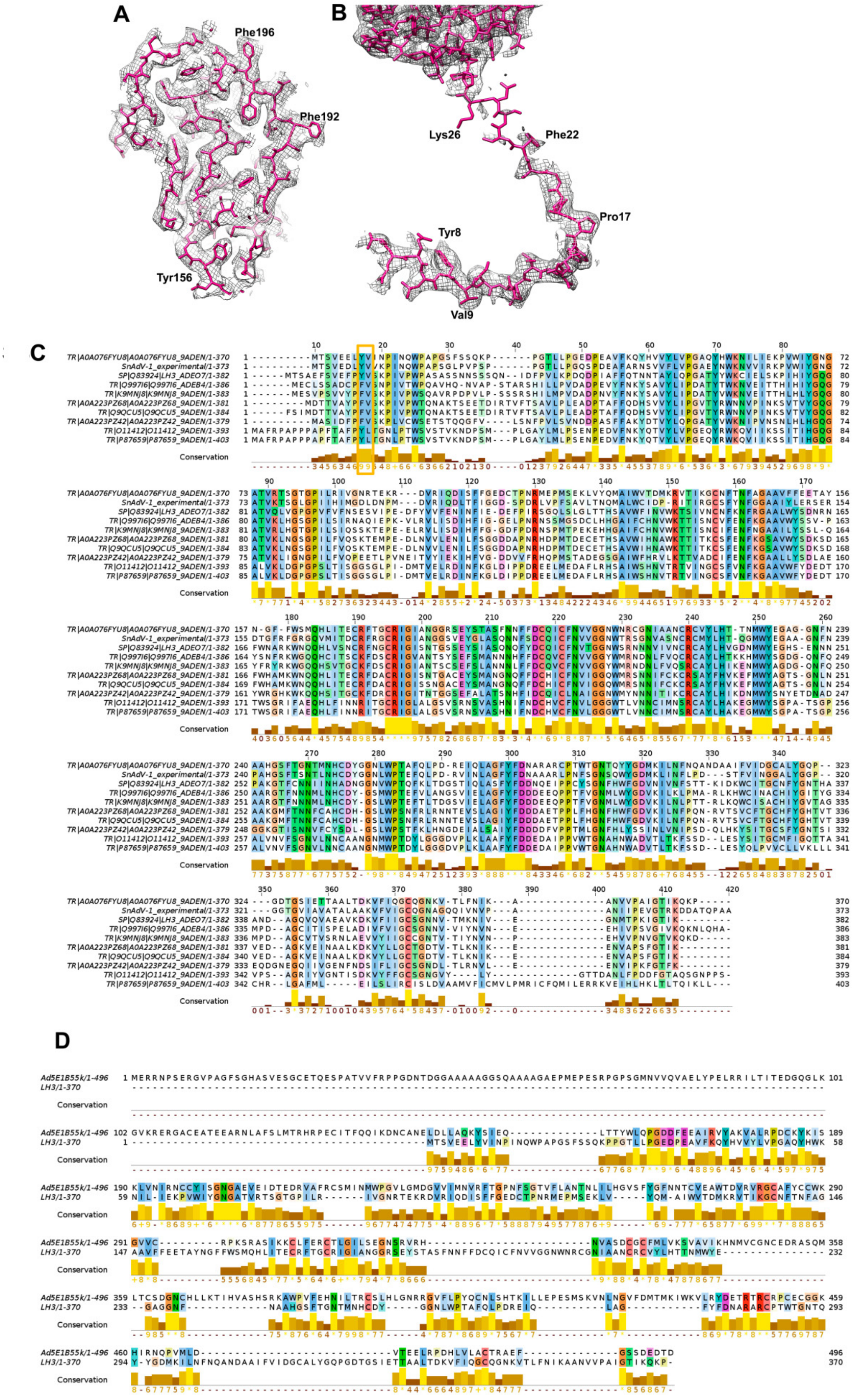


**Figure S4. LAdV-2 minor coat proteins on the outer capsid surface: LH3. (A)** Detail of the molecular model fit on the density of the β-helix domain. A region including the mobile loop Asp155-Ser162 discussed in the text is shown, as seen from outside the capsid. **(B)** Fit of the triskelion and column domains in the map. **(C)** Conservation of the LH3 protein sequence in atadenoviruses. The orange rectangle highlights the triskelion hydrophobic core. Sequence identifiers are as follows: TR|A0A076FYU8|A0A076FYU8_9ADEN: lizard adenovirus 2; SnAdV-1_experimental: snake adenovirus type 1, as determined in (Menéndez-Conejero *et al.*, 2017); SP|Q83924|LH3_ADEO7: ovine adenovirus D type 7 (isolate OAV287); TR|Q997I6|Q997I6_ADEB4: bovine adenovirus type 4 (BAdV-4); TR|K9MNJ8|K9MNJ8_9ADEN: bovine adenovirus type 6; TR|A0A223PZ68|A0A223PZ68_9ADEN, TR|Q9QCU5|Q9QCU5_9ADEN and TR|A0A223PZ42|A0A223PZ42_9ADEN: odocoileus adenovirus type 1; TR|O11412|O11412_9ADEN and TR|P87659|P87659_9ADEN: duck adenovirus 1. **(D)** Alignment of the HAdV-C5 E1B 55K and LAdV-2 LH3 protein sequences.

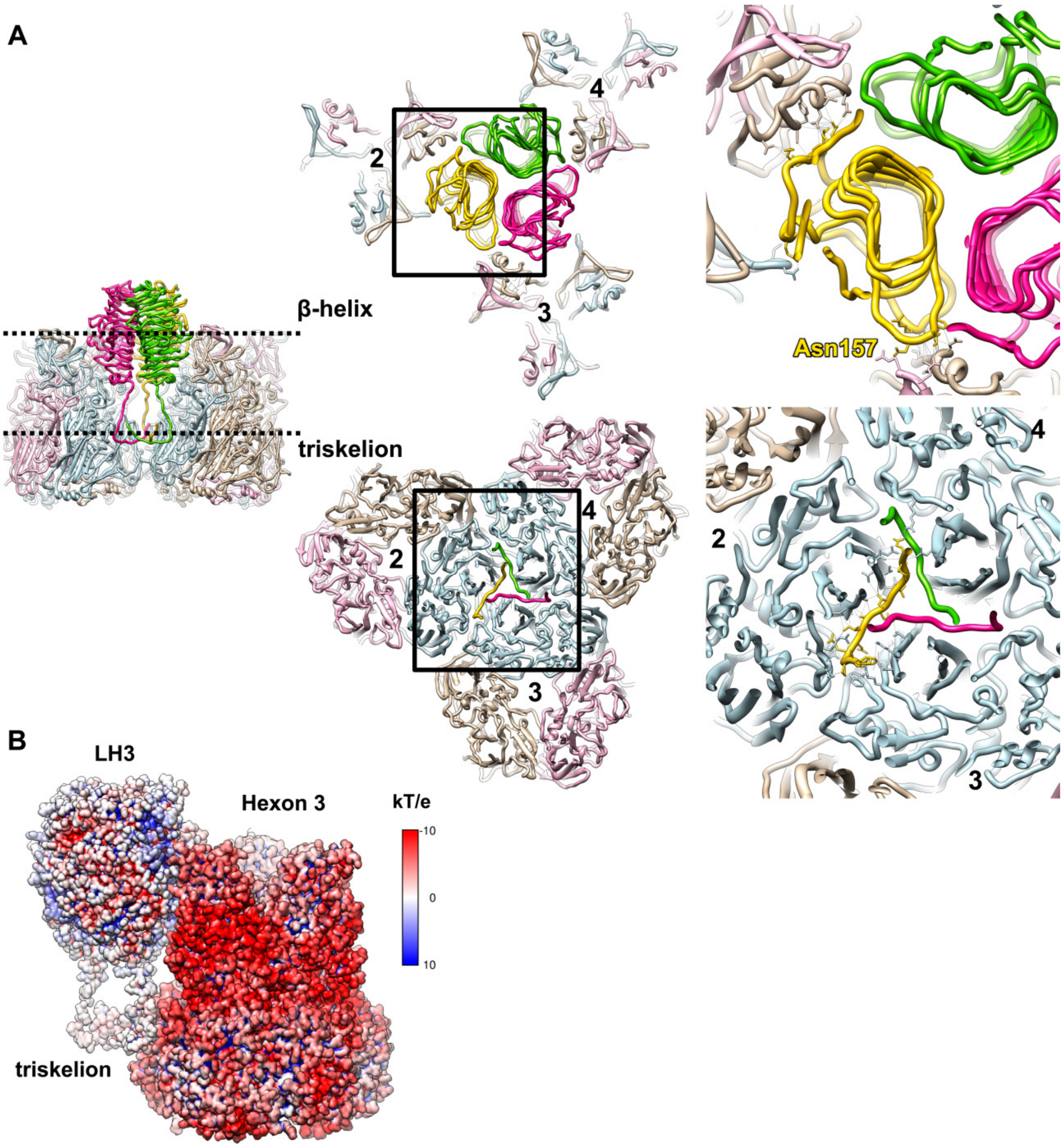


**Figure S5. Extended contact network between LH3 and the surrounding hexons. (A)** Left: inset showing the same side view as in Fig. 4C, with dotted black lines indicating the regions depicted in the rest of the panels, where the molecules are rendered in a view from outside the capsid. Slabs at the height of the LH3 **β-helix** or **triskelion** domains are shown, as indicated. The areas within the black rectangles are magnified at the right, with side chains rendered for those residues making contacts between the yellow LH3 monomer (chain S) and the surrounding hexon monomers. Numbers **2-4** identify the hexons in the AU. Residue **Asn157** is labelled to indicate the location of the LH3 mobile loop that was flexible in the crystal structure. **(B)** An LH3 trimer together with one of its hexon neighbours, represented as surfaces coloured by electrostatic potential, according to the scale at the right hand side.

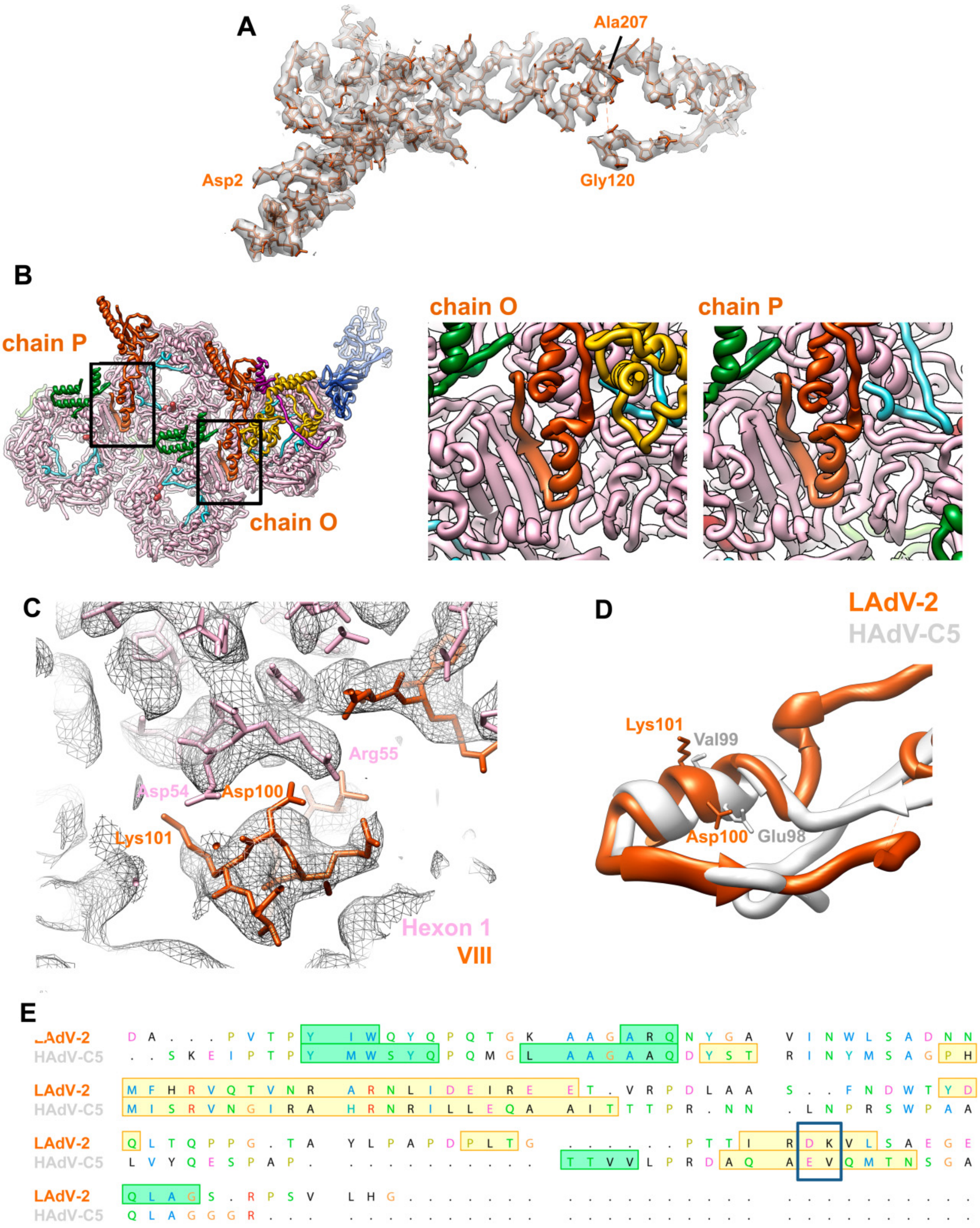


**Fig. S6. LAdV-2 internal minor coat proteins: polypeptide VIII. (A)** Fit of the complete protein VIII model on the cryo-EM map (semitransparent surface). **(B)** Protein VIII interaction with hexon by β-sheet augmentation. The AU is shown as in Fig. 1B, with the regions near the head domain of both protein VIII copies (black rectangles) magnified at the right. **(C)** Residues involved in possible electrostatic interactions between hexon and protein VIII, with the map rendered as a grey mesh. **(D)** Detail of the superimposed LAdV-2 and HAdV-C5 protein VIII structures, showing the location of the equivalent residues in the head domain. **(D)** Structureguided sequence alignment. Only the N-terminal fragment of protein VIII is shown. Green shading indicates β-sheets, yellow shading indicates α-helices. The blue rectangle highlights the charged residue pair in LAdV-2 and the equivalent region in HAdV-C5 protein VIII. Text colours follow the ClustalX scheme (Larkin *et al.*, 2007).

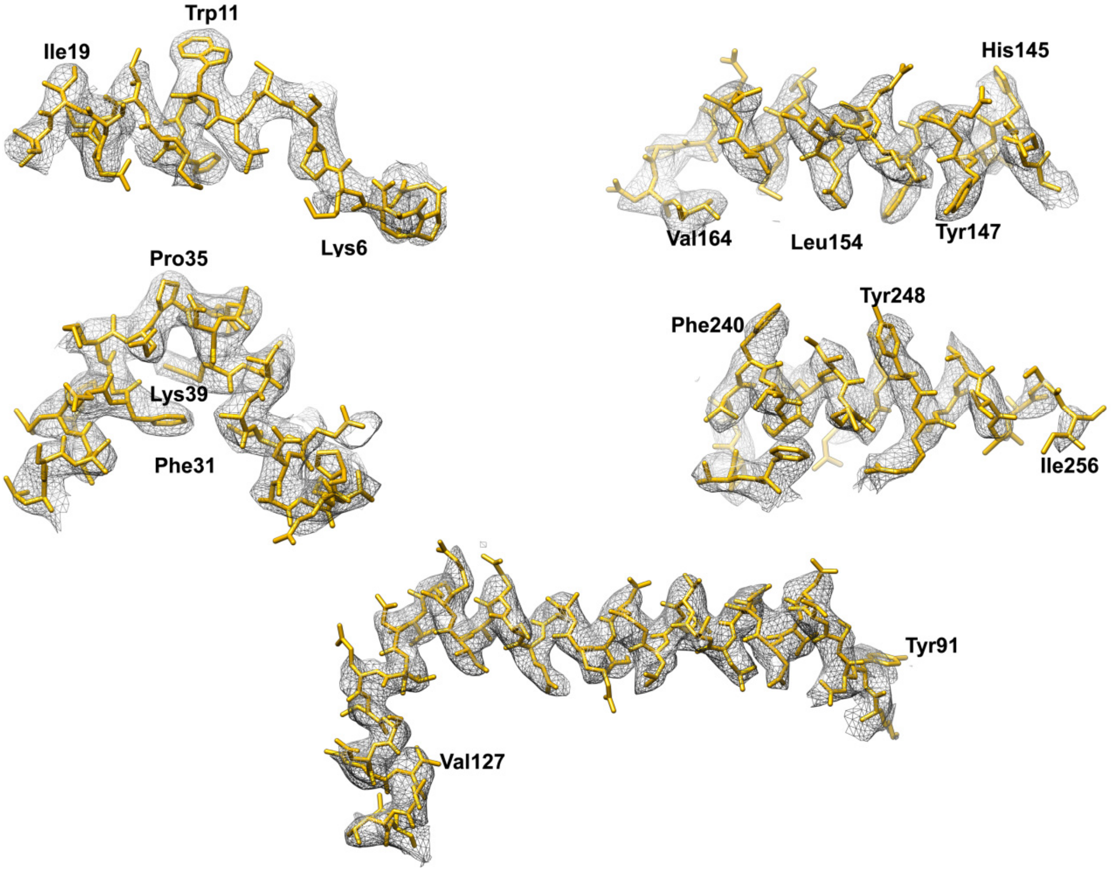


**Fig. S7. LAdV-2 internal minor coat proteins: polypeptide IIIa.** Details of several regions throughout the protein IIIa molecular model and their fit on the density map.

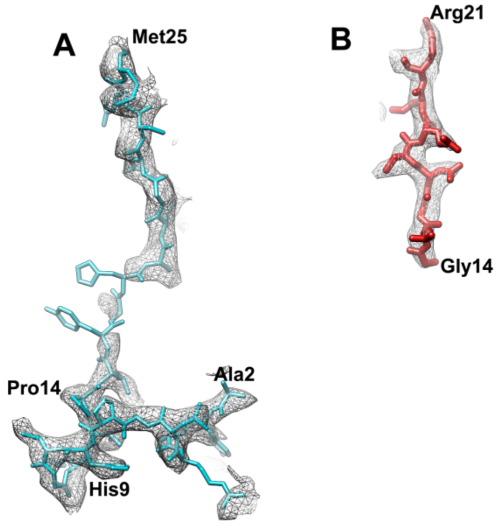


**C**

```
TR|A0A076FTF8|A0A076FTF8_9ADEN ---MAYSRLAPHCGLTPVYGH--AIGNSEML-GGFSWSSLGSSLSSGLSKIGSFLGST-- 52
SP|P24937|CAP6_ADE05           MEDINFASLAPRHGSRPFMGNWQDIGTSNMSGGAFSWGSLWSGIK---NFGSTVKNYGS 56
                                  : :: ***: *  *. *:   **.*:*  *.***.** *.:.     ::** : .

TR|A0A076FTF8|A0A076FTF8_9ADEN ----------ARRIGSSEGFQQAKQGFLKSGVIENVGALAGQTVSSLADIGRMKLEADLQ 102
SP|P24937|CAP6_ADE05           KAWNSSTGQMLRDKLKEQNFQQKVVDGLASGIS-----------GVVDLANQAVQNKIN 104
                                         *    ..:.***   . * **:            .:.*:..  :: .::

TR|A0A076FTF8|A0A076FTF8_9ADEN KLRERALGTPEQPPPLTQEQL------------AQLLASTQTELPPSA-STL--PMPIP 146
SP|P24937|CAP6_ADE05           SKLDP-RPPVEEPPPAVETVSPEGRGEKRPRPDREETL-VTQIDEPPSYEEALKQGLPTT 162
                               .  :       *:***  .:                  : *   ** : ***   .:*   :*

TR|A0A076FTF8|A0A076FTF8_9ADEN APTPPLVSAVRPEMVR-PEILPPPVVDR--GVALGP--LVEERPAR---PIAVPGSRPRK 198
SP|P24937|CAP6_ADE05           RPIAPMATGVLGQHTPVTLDLPPPADTQQKPVLPGPTAVVVTRPSRASLRRAASGPRSLR 222
                                *  *:.:.*  : .      ****.  :    *  **  :*  **:*      *. * *  :

TR|A0A076FTF8|A0A076FTF8_9ADEN RKRVRGWGTALEEMLGNGVDFGSKRYCY 226
SP|P24937|CAP6_ADE05           PVASGNWQSTLNSIVGLGVQSLKRRRCF 250
                                   .* ::*:.::***:  .:* *:
```

**D**

```
TR|A0A076FTB0|A0A076FTB0_9ADEN MSILISPSDNTGWGLGNCR-IRGTGARFTLSTPVPVRHYYRAQWGSKTGRVTEEH-LRRN 58
SP|P68951|NP_ADE05             MSILISPSNNTGWGLRFPSKMFGGAKKRSDQHPVRVRGHYRAPWGAHKRGRTGRTTVDDA 60
                               ********:*******      : * *. : : .  ** ** :*** **::.    *  .   :

TR|A0A076FTB0|A0A076FTB0_9ADEN LRKYRRRHHRRSIKTDRLLRSTPDTI-LIE--------TAKRV------RRTAHRRARR- 102
SP|P68951|NP_ADE05             IDAV--------VEEARNYTPTPPPVSTVDAAIQTVVRGARRYAKMKRRRRRVARRHRRR 112
                               :           ::  *     **  :  ::          *:*        ** . ** **

TR|A0A076FTB0|A0A076FTB0_9ADEN ----TTRGGPVLV-KVSRRRRGPRLRYTRRI---------------------------- 128
SP|P68951|NP_ADE05             PGTAAQRAAAALLNRARRTGRRAAMRAARRLAAGIVTVPPRSRRRAAAAAAAAISAMTQG 172
                                   : *.. .*: :. *  *   :* :**:

TR|A0A076FTB0|A0A076FTB0_9ADEN --------------------------
SP|P68951|NP_ADE05             RRGNVYWVRDSVSGLRVPVRTRPPRN 198
```

**Fig. S8. LAdV-2 proteins VI and VII. (A)** Example of density inside the hexon cavity proposed to correspond to the $pVI_N$ peptide (modelled in cyan). **(B)** Example of density inside the hexon cavity proposed to correspond to the $pVII_{N2}$ peptide (modelled in red). **(C)** Sequence alignment of protein VI in LAdV-2 (A0A076FTF8_9ADEN) and HAdV-C5 (CAP6_ADE05) with the residues modelled highlighted in cyan. **(D)** Sequence alignment of protein VII in LAdV-2 (A0A076FTB0_9ADEN) and HAdV-C5 (NP_ADE05) with the residues modelled highlighted in red. In (C) and (D), residues highlighted in yellow indicate those traced in the

HAdV-C5 structure (PDB ID 6B1T) (Dai *et al.*, 2017). Arrows indicate maturation cleavage sites.

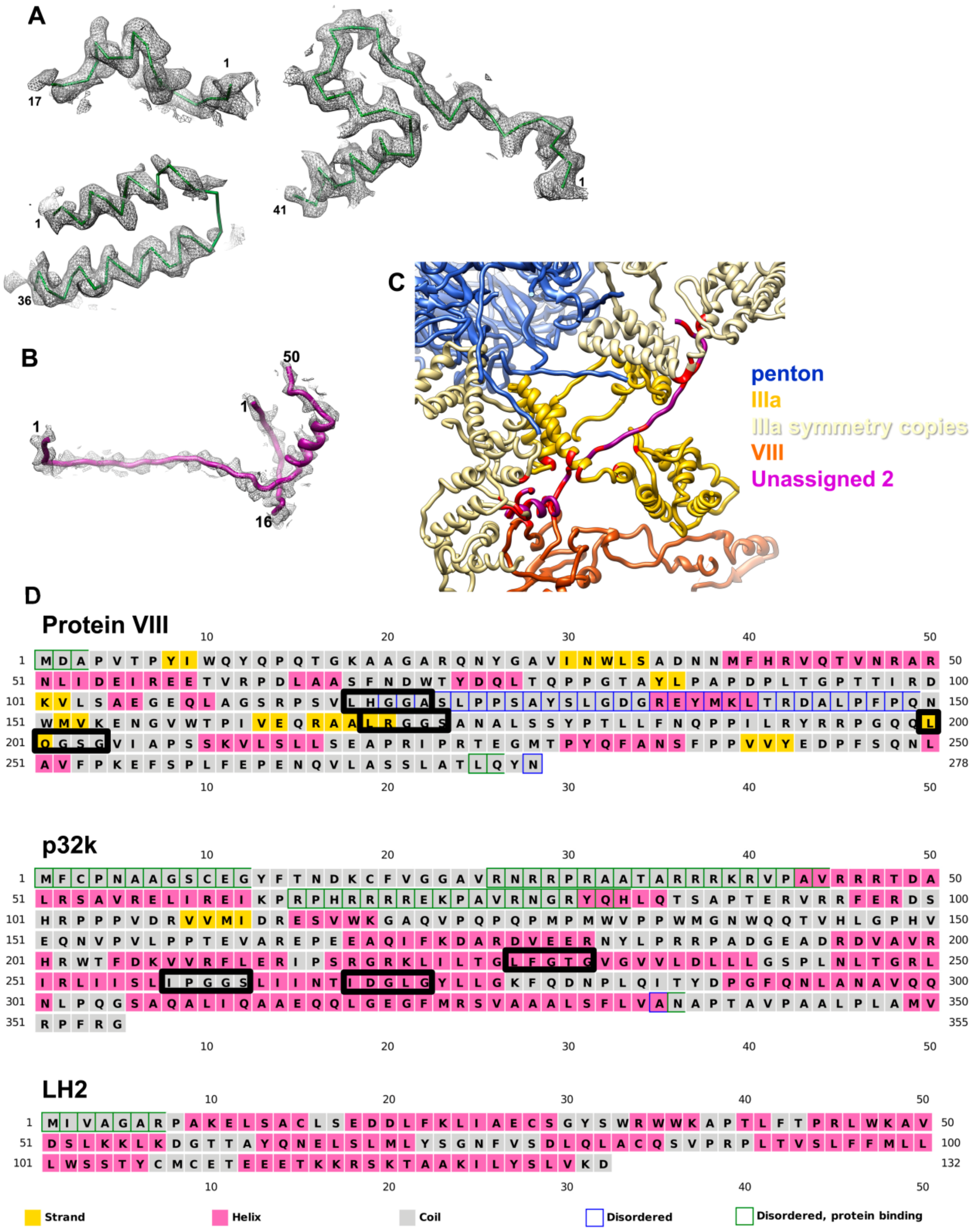


**Fig. S9. Interpretation of unassigned density on the inner capsid surface. (A)** Fit of the poly-alanine peptides traced in the U1 density of the cryo-EM map. **(B)** Fit of the poly-alanine peptides traced in the U2 density. **(C)** View of the vertex region with the residues potentially interacting with the U2 peptides in red. (D) PsiPred (Buchan and Jones, 2019) server secondary structure and disorder predictions for the precursor form of protein VIII and the genus specific proteins p32k (Uniprot A0A076FTA2_9ADEN, 355 amino acids) and LH2 (Uniprot A0A076FTE8_9ADEN, 132 amino acids). A large part of the cleaved peptides in VIII, and the arginine rich amino-terminal region of p32k, are predicted to be disordered. Black thick rectangles highlight consensus cleavage motifs for the maturation protease.

**Supplementary Files**

**File S1.** PDB validation report.
