## Supplemental file S1 for "Near Atomic Structure of an Atadenovirus Reveals a Conserved Capsid-Binding Motif and Intergenera Variations in Cementing Proteins"

### Full wwPDB/EMDataBank EM Map/Model Validation Report (i)

Feb 9, 2019 – 06:26 pm GMT

| | | |
|---|---|---|
| PDB ID | : | 6QI5 |
| EMDB ID: | : | EMD-4551 |
| Title | : | Near Atomic Structure of an Atadenovirus Shows a possible gene duplication event and Intergenera Variations in Cementing Proteins |
| Deposited on | : | 2019-01-17 |
| Resolution | : | 3.40 Å(reported) |

This is a Full wwPDB/EMDataBank EM Map/Model Validation Report.

This report is produced by the wwPDB biocuration pipeline after annotation of the structure.

We welcome your comments at **
A user guide is available at
https://www.wwpdb.org/validation/2017/EMValidationReportHelp
with specific help available everywhere you see the (i) symbol.

| | | |
|---|---|---|
| MolProbity | : | 4.02b-467 |
| Percentile statistics | : | 20171227.v01 (using entries in the PDB archive December 27th 2017) |
| Ideal geometry (proteins) | : | Engh & Huber (2001) |
| Ideal geometry (DNA, RNA) | : | Parkinson et. al. (1996) |
| Validation Pipeline (wwPDB-VP) | : | 2.1 |

CONFIDENTIAL VALIDATION REPORT

### 1 Overall quality at a glance (i)

The following experimental techniques were used to determine the structure:
*ELECTRON MICROSCOPY*

The reported resolution of this entry is 3.40 Å.

Percentile scores (ranging between 0-100) for global validation metrics of the entry are shown in the following graphic. The table shows the number of entries on which the scores are based.

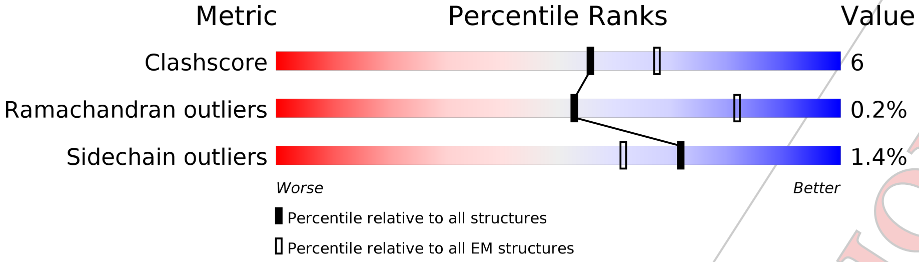


| Metric | Whole archive (#Entries) | EM structures (#Entries) |
|---|---|---|
| Clashscore | 136327 | 1886 |
| Ramachandran outliers | 132723 | 1663 |
| Sidechain outliers | 132532 | 1531 |

The table below summarises the geometric issues observed across the polymeric chains. The red, orange, yellow and green segments on the bar indicate the fraction of residues that contain outliers for >=3, 2, 1 and 0 types of geometric quality criteria. A grey segment represents the fraction of residues that are not modelled. The numeric value for each fraction is indicated below the corresponding segment, with a dot representing fractions <=5%

| Mol | Chain | Length | Quality of chain |
|---|---|---|---|
| 1 | A | 909 | 82% 17% |
| 1 | B | 909 | 79% 20% •• |
| 1 | C | 909 | 82% 18% |
| 1 | D | 909 | 81% 19% |
| 1 | E | 909 | 82% 18% |
| 1 | F | 909 | 84% 15% |
| 1 | G | 909 | 84% 15% • |
| 1 | H | 909 | 81% 18% |
| 1 | I | 909 | 82% 17% • |

*Continued on next page...*

CONFIDENTIAL VALIDATION REPORT

*Continued from previous page...*

| Mol | Chain | Length | Quality of chain |
|---|---|---|---|
| 1 | J | 909 | 84% 16% |
| 1 | K | 909 | 83% 16% • |
| 1 | L | 909 | 82% 16% • |
| 2 | Q | 370 | 84% 15% • |
| 2 | R | 370 | 83% 16% • |
| 2 | S | 370 | 83% 16% • |
| 2 | T | 370 | 86% 13% • |
| 3 | O | 278 | 60% 7% 33% |
| 3 | P | 278 | 60% 7% 33% |
| 4 | N | 609 | 37% 5% 58% |
| 5 | M | 451 | 80% 20% |

CONFIDENTIAL VALIDATION REPORT

### 2 Entry composition (i)

There are 5 unique types of molecules in this entry. The entry contains 106009 atoms, of which 0 are hydrogens and 0 are deuteriums.

In the tables below, the AltConf column contains the number of residues with at least one atom in alternate conformation and the Trace column contains the number of residues modelled with at most 2 atoms.

- Molecule 1 is a protein called Hexon protein.

| Mol | Chain | Residues | Atoms | | | | | AltConf | Trace |
|---|---|---|---|---|---|---|---|---|---|
| 1 | A | 905 | Total 7168 | C 4547 | N 1226 | O 1366 | S 29 | 0 | 0 |
| 1 | L | 906 | Total 7174 | C 4550 | N 1227 | O 1368 | S 29 | 0 | 0 |
| 1 | K | 908 | Total 7184 | C 4556 | N 1229 | O 1370 | S 29 | 0 | 0 |
| 1 | J | 906 | Total 7171 | C 4548 | N 1227 | O 1368 | S 28 | 0 | 0 |
| 1 | H | 905 | Total 7166 | C 4545 | N 1226 | O 1367 | S 28 | 0 | 0 |
| 1 | I | 906 | Total 7171 | C 4548 | N 1227 | O 1368 | S 28 | 0 | 0 |
| 1 | G | 906 | Total 7171 | C 4548 | N 1227 | O 1368 | S 28 | 0 | 0 |
| 1 | F | 906 | Total 7172 | C 4548 | N 1227 | O 1368 | S 29 | 0 | 0 |
| 1 | E | 906 | Total 7171 | C 4548 | N 1227 | O 1368 | S 28 | 0 | 0 |
| 1 | D | 906 | Total 7174 | C 4550 | N 1227 | O 1368 | S 29 | 0 | 0 |
| 1 | C | 907 | Total 7179 | C 4553 | N 1228 | O 1369 | S 29 | 0 | 0 |
| 1 | B | 904 | Total 7161 | C 4542 | N 1225 | O 1365 | S 29 | 0 | 0 |

- Molecule 2 is a protein called Protein LH3.

| Mol | Chain | Residues | Atoms | | | | | AltConf | Trace |
|---|---|---|---|---|---|---|---|---|---|
| 2 | S | 367 | Total 2874 | C 1830 | N 494 | O 529 | S 21 | 0 | 0 |
| 2 | T | 367 | Total 2874 | C 1830 | N 494 | O 529 | S 21 | 0 | 0 |
| 2 | R | 367 | Total 2874 | C 1830 | N 494 | O 529 | S 21 | 0 | 0 |

Continued on next page...

CONFIDENTIAL VALIDATION REPORT

*Continued from previous page...*

| Mol | Chain | Residues | Atoms | | | | | AltConf | Trace |
|---|---|---|---|---|---|---|---|---|---|
| | | | Total | C | N | O | S | | |
| 2 | Q | 367 | 2874 | 1830 | 494 | 529 | 21 | 0 | 0 |

- Molecule 3 is a protein called Pre-hexon-linking protein VIII.

| Mol | Chain | Residues | Atoms | | | | | AltConf | Trace |
|---|---|---|---|---|---|---|---|---|---|
| | | | Total | C | N | O | S | | |
| 3 | P | 186 | 1452 | 922 | 248 | 280 | 2 | 0 | 0 |
| 3 | O | 186 | 1452 | 922 | 248 | 280 | 2 | 0 | 0 |

- Molecule 4 is a protein called PIIIa.

| Mol | Chain | Residues | Atoms | | | | | AltConf | Trace |
|---|---|---|---|---|---|---|---|---|---|
| | | | Total | C | N | O | S | | |
| 4 | N | 255 | 1980 | 1250 | 346 | 378 | 6 | 0 | 0 |

- Molecule 5 is a protein called Penton protein.

| Mol | Chain | Residues | Atoms | | | | | AltConf | Trace |
|---|---|---|---|---|---|---|---|---|---|
| | | | Total | C | N | O | S | | |
| 5 | M | 451 | 3567 | 2270 | 598 | 687 | 12 | 0 | 0 |

CONFIDENTIAL VALIDATION REPORT

### 3 Residue-property plots

These plots are drawn for all protein, RNA and DNA chains in the entry. The first graphic for a chain summarises the proportions of the various outlier classes displayed in the second graphic. The second graphic shows the sequence view annotated by issues in geometry. Residues are color-coded according to the number of geometric quality criteria for which they contain at least one outlier: green = 0, yellow = 1, orange = 2 and red = 3 or more. Stretches of 2 or more consecutive residues without any outlier are shown as a green connector. Residues present in the sample, but not in the model, are shown in grey.

- Molecule 1: Hexon protein

Chain A:

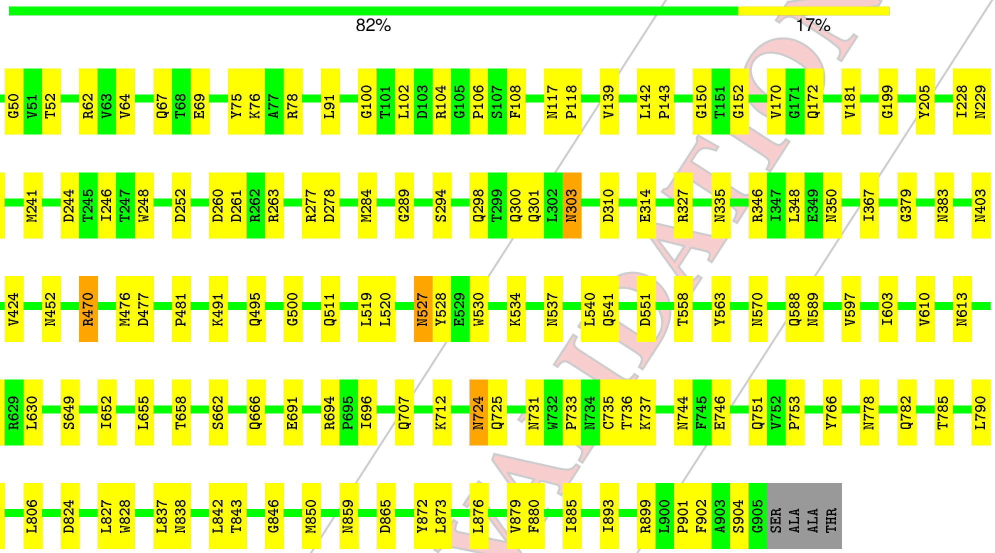


- Molecule 1: Hexon protein

Chain L:

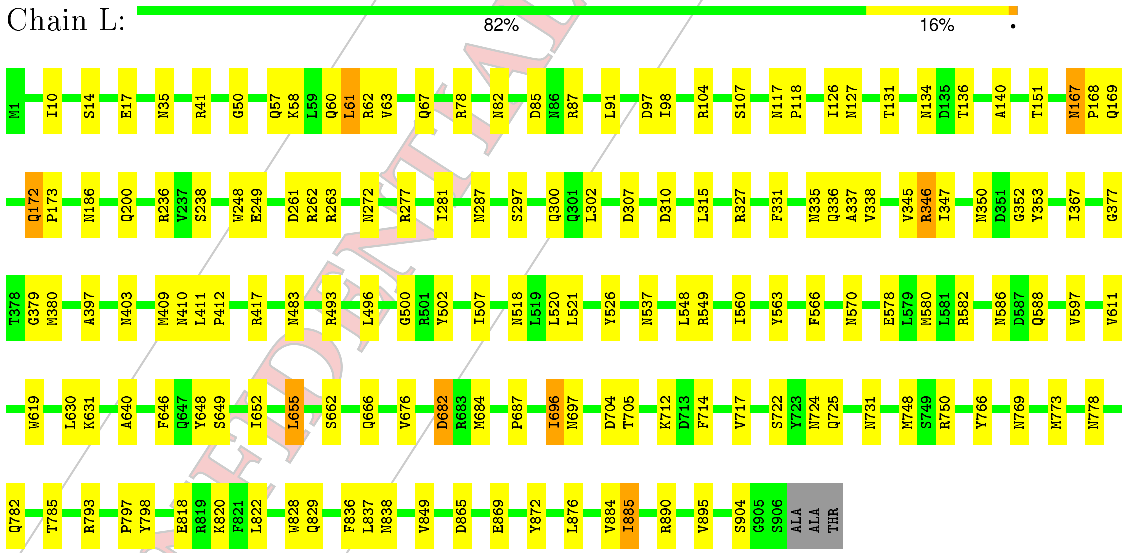


- Molecule 1: Hexon protein

CONFIDENTIAL VALIDATION REPORT

Chain K: 83% 16% •

M1 R5 R13 K16 I36 G50 V51 T52 E69 K76 A77 R78 N82 D97 G100 R104 G105 P106 S107 F108 K109 P110 Y111 N117 P118 Q141 L142 P143 Q144 T145 Y146 G150 N167 P168 Q169 V170 G171 Q172 N186 L189 S197

I228 N234 G235 R236 I246 T247 W248 D252 A256 R277 D278 M283 N287 N291 S296 N303 I304 D307 D310 E314 R327 R346 I347 N350 A361 F362 P363 P364 H365 V366 I367 G379 N383 Q386 I399 I404 N410

N414 L415 K416 I460 R470 M476 P481 K491 Q495 N499 G500 V509 L520 Y531 F532 R533 K534 D535 D551 T558 Y563 N570 R582 Q588 N589 V597 I614 R623 S626 I652 L655 T658 S662 H663 T664

F665 Q666 R667 C668 D673 P677 G680 M684 F690 K693 R694 P695 I696 Q707 K712 D713 F714 Q718 N724 Q725 N731 T736 K737 N743 Y755 G756 N778 C779 T787 N788 V789 L790 L791 E792 R793 H796 N810 Q815 R819

W828 Q829 I830 L837 N838 L842 Q847 S857 L858 Q861 F862 D865 L873 L876 V879 F880 I885 T889 I893 S894 V895 A896 Y897 P901 A908 THR

• Molecule 1: Hexon protein

Chain J: 84% 16%

MET E2 R13 E17 Y18 L19 N22 L23 N35 K39 G50 D70 Y75 N82 F96 D97 I98 L102 D103 R104 Y111 N117 P118 S128 Q141 L142 P143 Q159 Q160 N167 P168 Q169 V170 G171 N174 P178 R191 I192 Q201

Y205 I228 M241 D244 T245 I246 T247 W248 D252 A256 D260 D261 R262 R263 A266 R277 M284 N287 S297 Q298 T299 Q300 I304 D307 D310 R327 N335 Q336 A337 V338 R346 I347 L348 E349 N350 D351 G352 N359 H365

G377 A393 A397 L398 I399 L415 K416 R417 Y421 S422 N423 V424 A425 Y450 I451 N452 G453 R454 I460 V461 D466 W471 R486 R493 G500 V509 P510 Q511 L520 L521 Y526 N527 Y528 E529 W530 K534 N537 L540 I560 N570

Q588 N589 V597 V611 V612 N613 S626 R629 L630 S649 I652 S662 H663 T664 C668 F690 K693 N709 K712 S722 R750 Q751 V752 P753 I775 W776 N777 N778 Q782 Q783 K784 N788 E792 R793 L806 S807 T817 W828 Q829

S834 N838 T843 G846 N859 M860 Q861 Y872 L876 V884 I885 R890 V895 S906 A907 ALA THR

• Molecule 1: Hexon protein

Chain H: 81% 18%

MET E2 N22 N35 K39 T52 R62 N71 V72 N73 V89 D97 G100 T101 L102 D103 R104 K109 P110 Y111 N117 P118 A140 Q144 A148 G149 G150 E155 G163 N167 P168 Q169 V170 N174 N185 L189 G190 R191 Q200

Y205 Y208 I228 N229 T230 R236 D244 T245 I246 T247 W248 P251 D252 A253 H254 D261 N272 R277 D278 M283 N287 G293 S297 D307 D310 R311 N312 S313 E314 R327 N335 Q336 A337 R346 I347 L348 Y353 P363 P364 H365

V366 I367 T381 V382 N383 T392 L398 I399 I404 N410 Y421 V424 A425 L428 Y432 N452 I460 V475 M476 D477 P481 F482 N487 R493 S494 Q495 G500 D504 V509 P510 Q511 L519 L520 L521 Y526 N527 Y528 E529 W530

L540 Q541 R549 T558 Q559 I560 N561 L562 Y563 Q588 N589 G595 A596 V597 N598 N599 L600 V611 R623 S626 L630 Q642 D643 P644 N645 I652 L655 S662 C668 R694 P695 I696 E701 G702 S708 K712 A720 N724 Q725 N731

CONFIDENTIAL VALIDATION REPORT

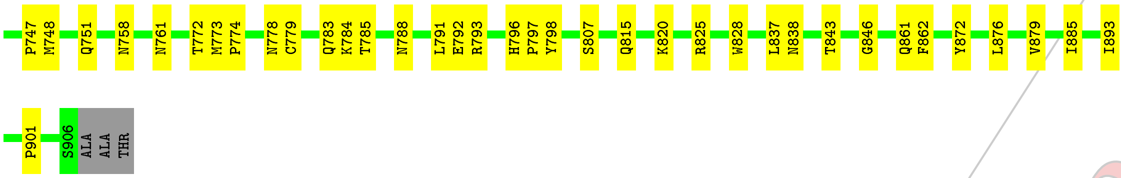


• Molecule 1: Hexon protein

Chain I:

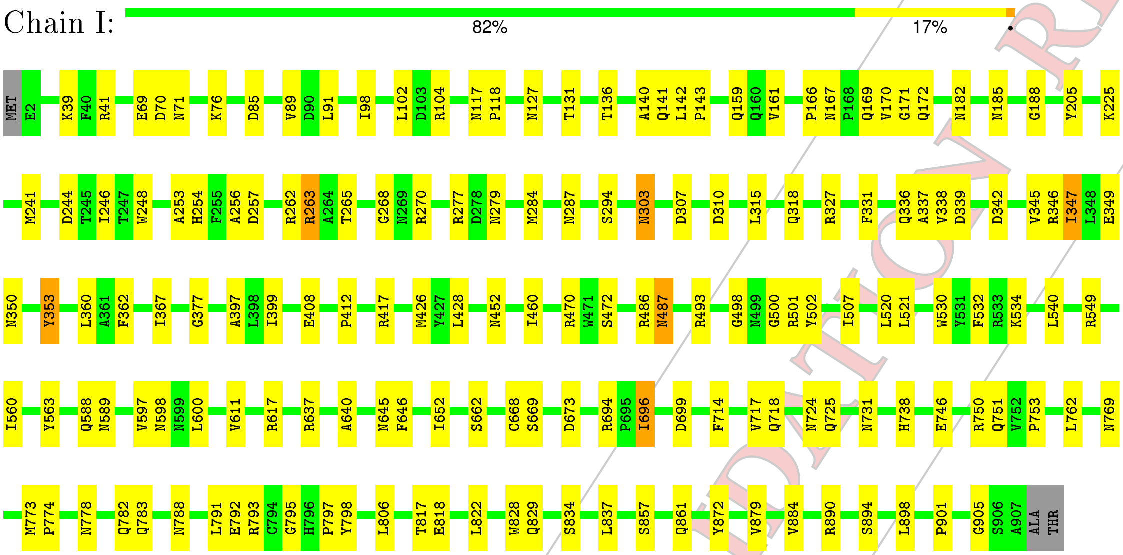


• Molecule 1: Hexon protein

Chain G:

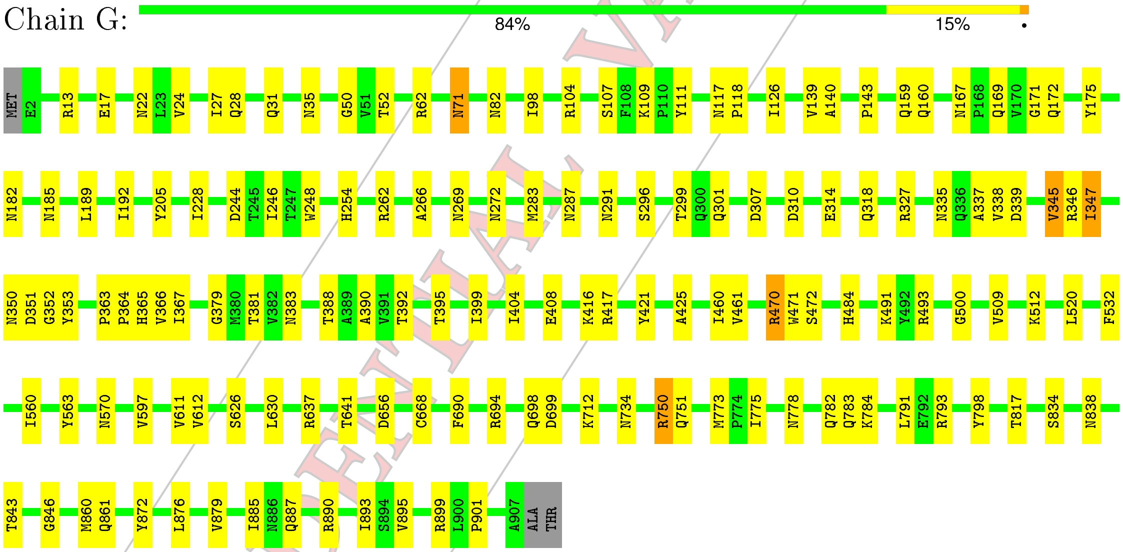


• Molecule 1: Hexon protein

Chain F:

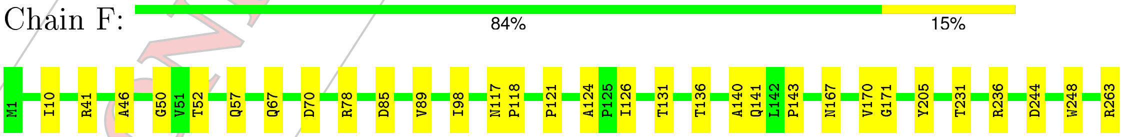


CONFIDENTIAL VALIDATION REPORT

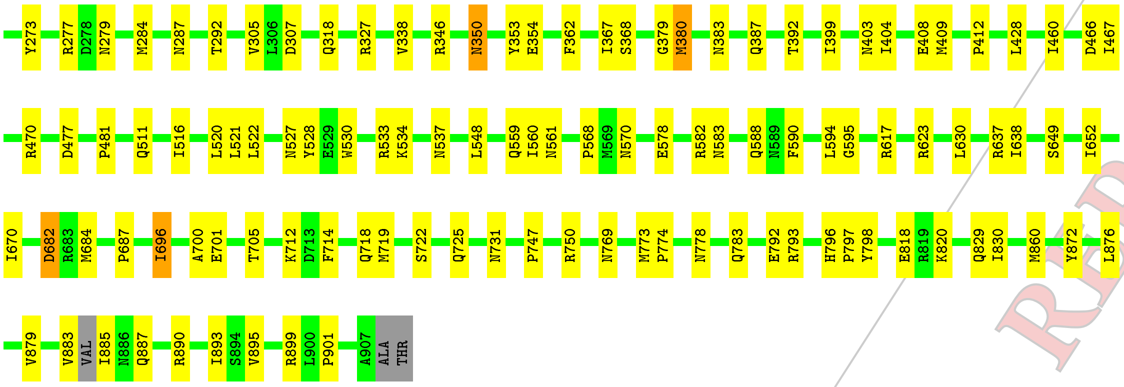


- Molecule 1: Hexon protein

Chain E:

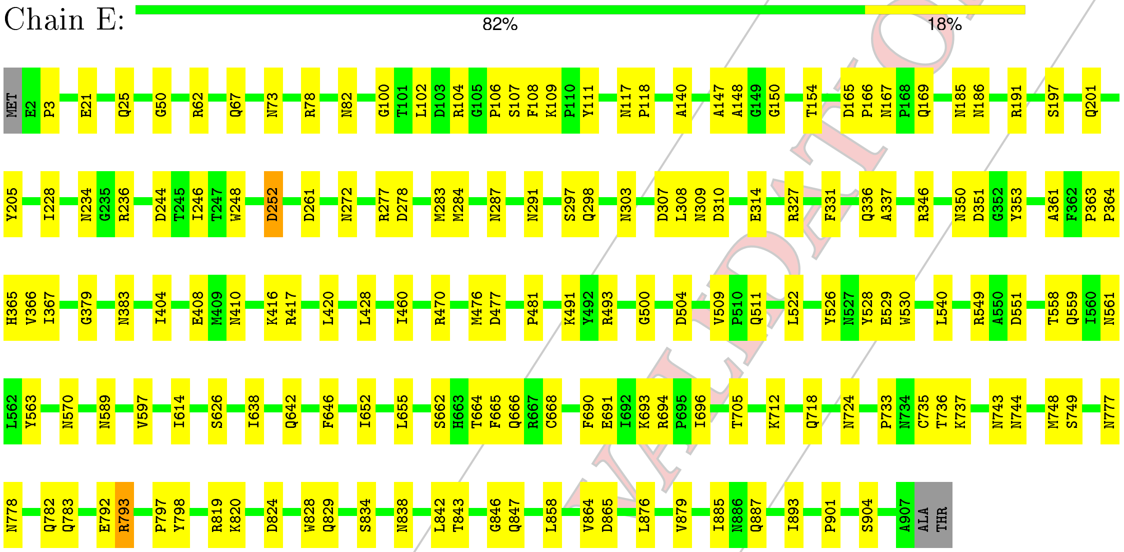


- Molecule 1: Hexon protein

Chain D:

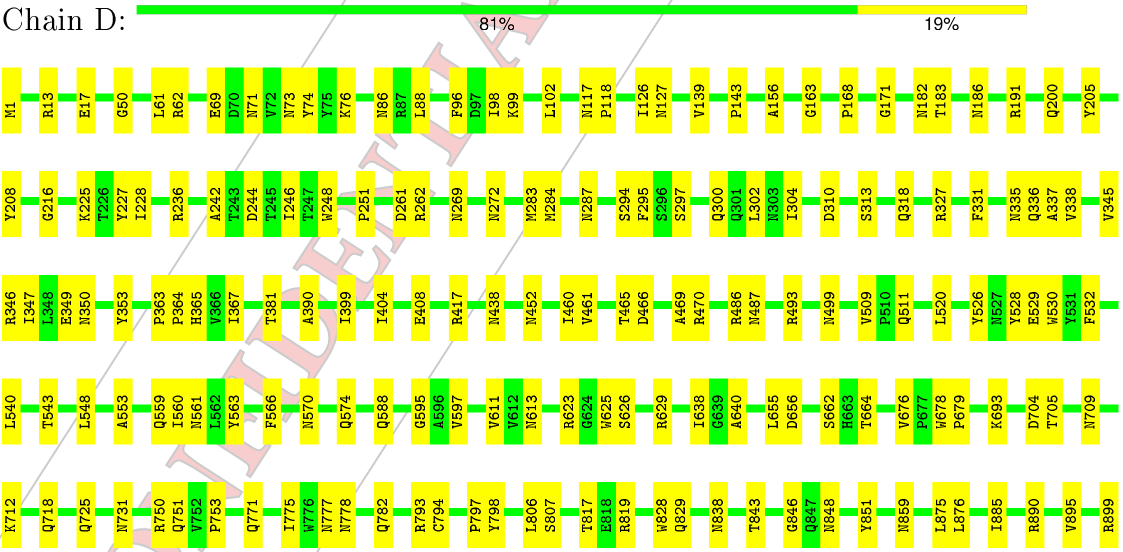


CONFIDENTIAL VALIDATION REPORT

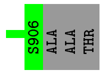


• Molecule 1: Hexon protein

Chain C:

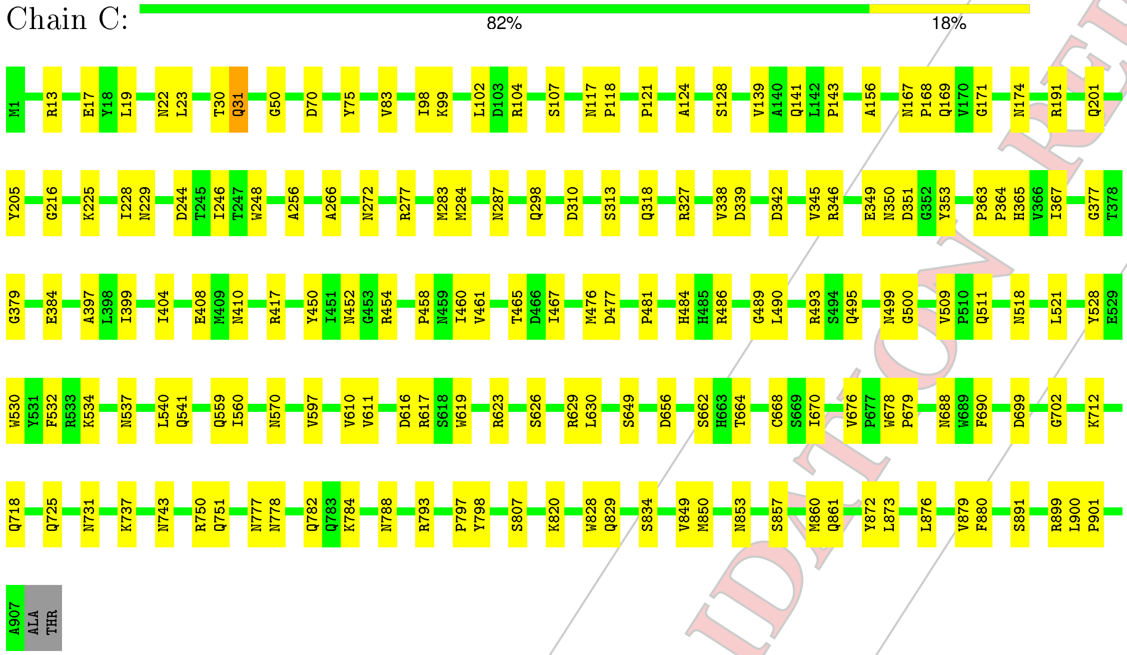


• Molecule 1: Hexon protein

Chain B:

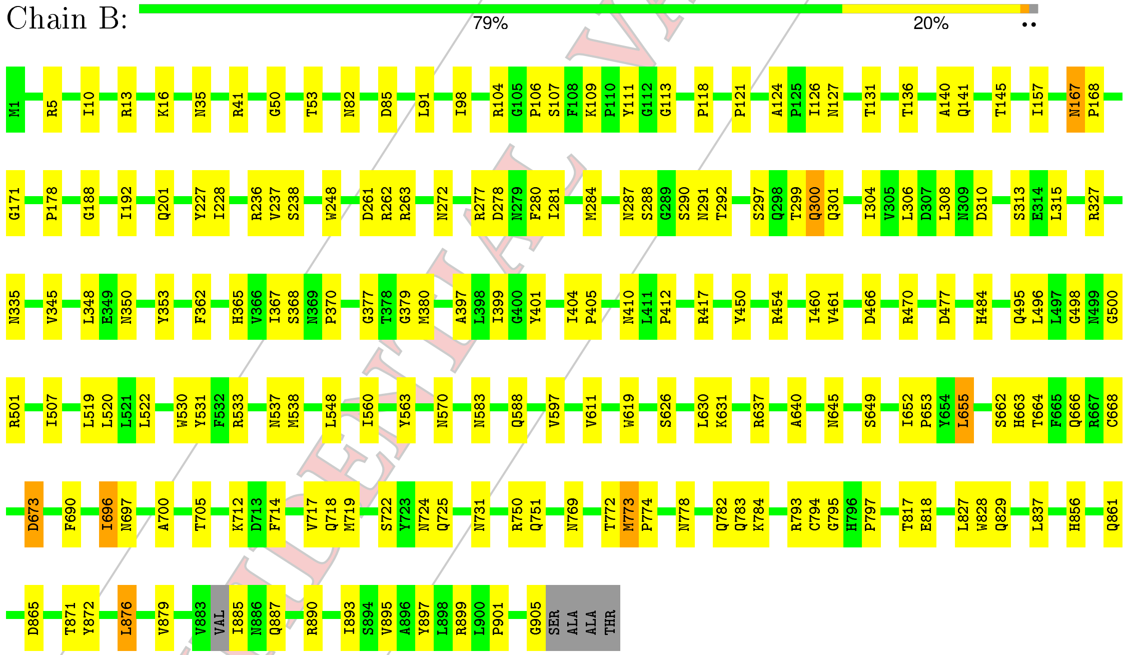


• Molecule 2: Protein LH3

Chain S:

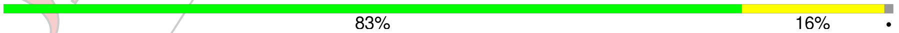


CONFIDENTIAL VALIDATION REPORT

MET THR SER V4 P17 L49 Q54 K58 G70 I83 L84 R90 T91 D95 V96 R97 D100 I126 R133 V134 T135 N140 F141 A148 V149 F150 F151 E152 E153 Y156 W161 S162 M163 Q164 I167 R171 F172 I179 R184 N194

F195 I200 N214 I215 A216 A217 N218 C219 R220 C221 N248 N251 H252 C253 G256 G257 N258 T262 F278 Y294 Y295 G296 I300 F303 G316 E330 T331 T332 L335 G343 V349 F352 G364 P370

• Molecule 2: Protein LH3

Chain T: 86% 13% •

MET THR SER V4 I13 A18 L49 G70 I83 L84 R90 R97 I126 R133 V134 T135 V149 F150 F151 E152 E153 Y156 M163 Q164 I167 F172 I179 R184 N194 F195 I200 N214 I215 A216 R220 C221 L224 N248

H252 C253 G256 G257 N258 T262 F278 G296 I300 F303 G316 E330 T331 T332 L335 G343 V349 F352 G364 P370

• Molecule 2: Protein LH3

Chain R: 83% 16% •

MET THR SER V4 N11 P17 L49 G70 I83 L84 R90 K93 R94 D95 V96 R97 D100 T109 E118 I126 R133 V134 T135 N140 F141 A148 V149 F150 F151 E152 E153 W161 S162 M163 Q164 I167 R171 F172 I179 Y187

N194 F195 I200 N214 I215 A216 A217 N218 C219 R220 C221 L224 H242 N248 N251 H252 C253 G256 G257 N258 T262 F278 Y294 Y295 G296 I300 F303 G316 E330 T331 T332 L335 G343 V349 F352 G364 P370

• Molecule 2: Protein LH3

Chain Q: 84% 15% •

MET THR SER V4 N11 P12 I13 P17 A18 L49 G70 I83 L84 R90 T91 D95 V96 R97 D100 I126 R133 V134 T135 N140 V149 F150 F151 E152 E153 Y156 W161 S162 M163 Q164 I167 F172 I179 R184 Y187 N194

F195 N214 I215 A216 A217 N218 C219 R220 M229 H242 N248 H252 C253 G256 G257 N258 T262 L266 F278 G296 I300 F303 G316 E330 T331 T332 L335 G343 V349 F352 G364 P370

• Molecule 3: Pre-hexon-linking protein VIII

Chain P: 60% 7% 33%

MET D2 Q15 R23 R50 R62 Y74 Q79 P89 T96 R99 G112 G120 GLY ALA SER LEU PRO PRO SER ALA TYR SER LEU GLY ASP GLY ARG GLU TYR MET LYS LEU THR ARG ASP ALA LEU PRO PHE PRO GLN ASN TRP MET VAL LYS GLU ASN

GLY VAL TRP THR PRO ILE VAL GLU GLN ARG ALA ALA LEU ARG GLY GLY SER ALA ASN ALA LEU SER SER TYR PRO THR LEU LEU PHE ASN GLN PRO PRO ILE LEU ARG TYR ARG ARG PRO GLY GLN GLN LEU GLN GLY SER GLY VAL ILE A207 V212 R221 I222 P223 R224 T225

N235 K255 P259 V267 A273 THR LEU GLN TYR ASN

• Molecule 3: Pre-hexon-linking protein VIII

CONFIDENTIAL VALIDATION REPORT

Chain O:

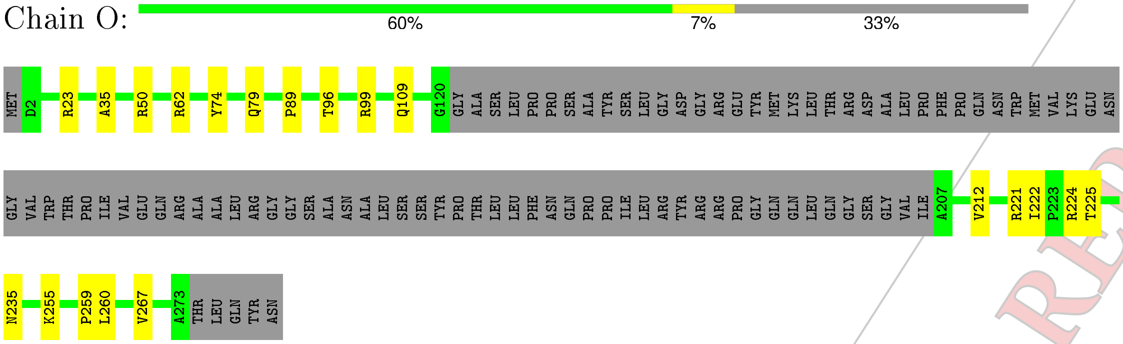


- Molecule 4: PIIIa

Chain N:

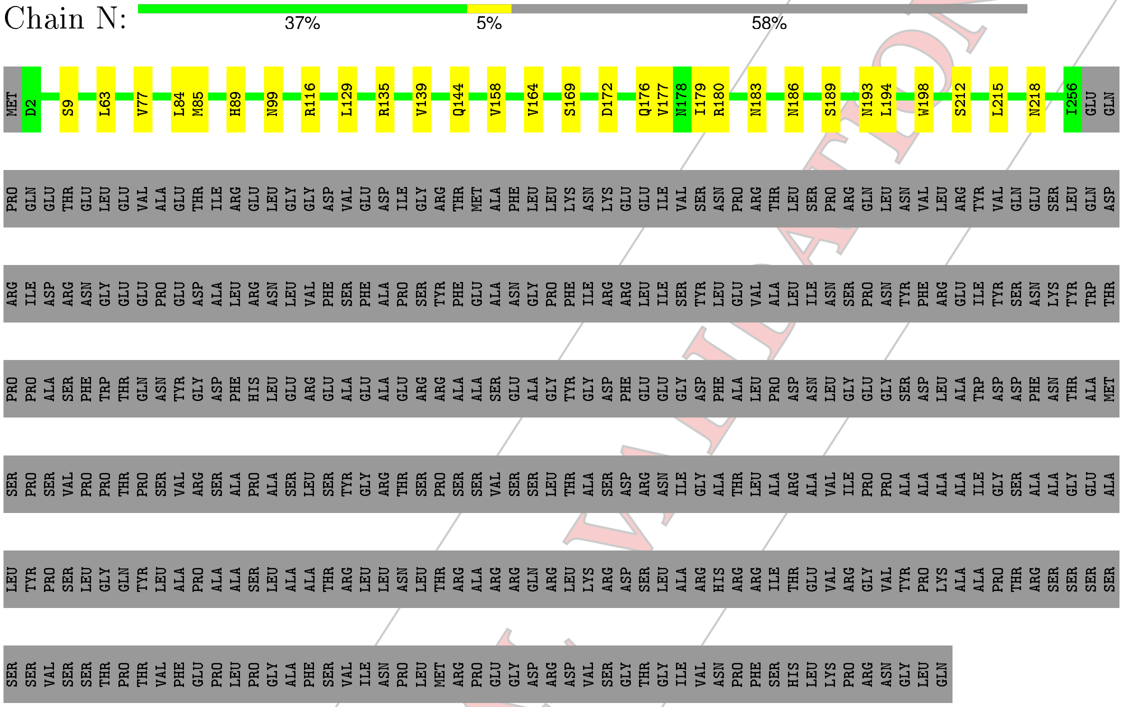


- Molecule 5: Penton protein

Chain M:

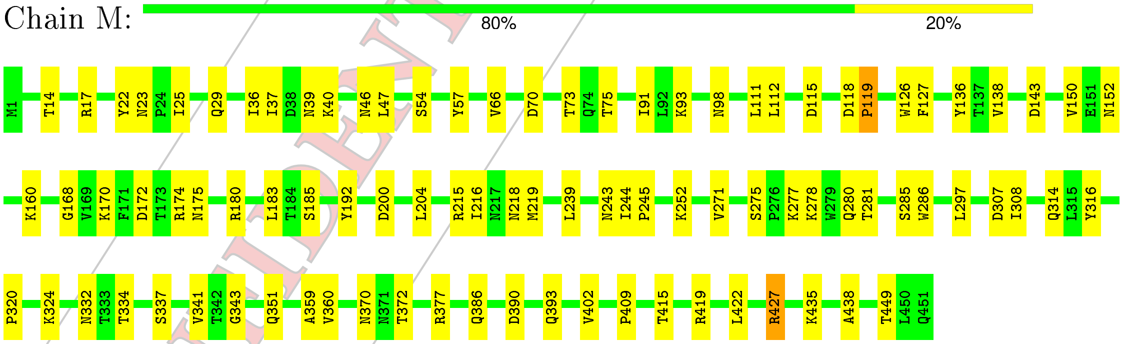


CONFIDENTIAL VALIDATION REPORT

### 4 Experimental information (i)

| Property | Value | Source |
|---|---|---|
| Reconstruction method | SINGLE PARTICLE | Depositor |
| Imposed symmetry | POINT, I | Depositor |
| Number of particles used | 16071 | Depositor |
| Resolution determination method | FSC 0.143 CUT-OFF | Depositor |
| CTF correction method | PHASE FLIPPING ONLY | Depositor |
| Microscope | FEI TITAN KRIOS | Depositor |
| Voltage (kV) | 300 | Depositor |
| Electron dose $(e^-/\text{Å}^2)$ | 54 | Depositor |
| Minimum defocus (nm) | Not provided | Depositor |
| Maximum defocus (nm) | Not provided | Depositor |
| Magnification | Not provided | Depositor |
| Image detector | FEI FALCON II (4k x 4k) | Depositor |

CONFIDENTIAL VALIDATION REPORT

### 5 Model quality (i)

#### 5.1 Standard geometry (i)

The Z score for a bond length (or angle) is the number of standard deviations the observed value is removed from the expected value. A bond length (or angle) with $|Z| > 5$ is considered an outlier worth inspection. RMSZ is the root-mean-square of all Z scores of the bond lengths (or angles).

| Mol | Chain | Bond lengths | | Bond angles | |
|---|---|---|---|---|---|
| | | RMSZ | #\|Z\| >2 | RMSZ | #\|Z\| >2 |
| 1 | A | 0.35 | 0/7373 | 0.59 | 4/10074 (0.0%) |
| 1 | B | 0.34 | 0/7365 | 0.60 | 6/10061 (0.1%) |
| 1 | C | 0.35 | 0/7384 | 0.59 | 1/10089 (0.0%) |
| 1 | D | 0.38 | 0/7379 | 0.59 | 0/10082 |
| 1 | E | 0.37 | 0/7376 | 0.60 | 2/10079 (0.0%) |
| 1 | F | 0.37 | 0/7376 | 0.60 | 4/10076 (0.0%) |
| 1 | G | 0.38 | 0/7376 | 0.59 | 0/10079 |
| 1 | H | 0.38 | 0/7371 | 0.61 | 3/10072 (0.0%) |
| 1 | I | 0.38 | 0/7376 | 0.61 | 6/10079 (0.1%) |
| 1 | J | 0.36 | 0/7376 | 0.58 | 2/10079 (0.0%) |
| 1 | K | 0.36 | 0/7389 | 0.59 | 2/10096 (0.0%) |
| 1 | L | 0.37 | 0/7379 | 0.61 | 5/10082 (0.0%) |
| 2 | Q | 0.34 | 0/2954 | 0.54 | 0/4016 |
| 2 | R | 0.34 | 0/2954 | 0.55 | 0/4016 |
| 2 | S | 0.34 | 0/2954 | 0.55 | 0/4016 |
| 2 | T | 0.34 | 0/2954 | 0.55 | 0/4016 |
| 3 | O | 0.34 | 0/1494 | 0.55 | 0/2048 |
| 3 | P | 0.34 | 0/1494 | 0.55 | 0/2048 |
| 4 | N | 0.30 | 0/2014 | 0.59 | 0/2741 |
| 5 | M | 0.30 | 0/3660 | 0.61 | 1/5005 (0.0%) |
| All | All | 0.36 | 0/108998 | 0.59 | 36/148854 (0.0%) |

Chiral center outliers are detected by calculating the chiral volume of a chiral center and verifying if the center is modelled as a planar moiety or with the opposite hand.A planarity outlier is detected by checking planarity of atoms in a peptide group, atoms in a mainchain group or atoms of a sidechain that are expected to be planar.

| Mol | Chain | #Chirality outliers | #Planarity outliers |
|---|---|---|---|
| 1 | A | 0 | 4 |
| 1 | B | 0 | 4 |
| 1 | C | 0 | 1 |
| 1 | D | 0 | 3 |
| 1 | E | 0 | 3 |

*Continued on next page...*

CONFIDENTIAL VALIDATION REPORT

*Continued from previous page...*

| Mol | Chain | #Chirality outliers | #Planarity outliers |
|---|---|---|---|
| 1 | G | 0 | 4 |
| 1 | H | 0 | 3 |
| 1 | I | 0 | 2 |
| 1 | J | 0 | 3 |
| 1 | K | 0 | 2 |
| 1 | L | 0 | 5 |
| 5 | M | 0 | 3 |
| All | All | 0 | 37 |

There are no bond length outliers.

All (36) bond angle outliers are listed below:

| Mol | Chain | Res | Type | Atoms | Z | Observed(°) | Ideal(°) |
|---|---|---|---|---|---|---|---|
| 1 | B | 348 | LEU | CA-CB-CG | 8.97 | 135.93 | 115.30 |
| 1 | L | 61 | LEU | CA-CB-CG | 7.24 | 131.96 | 115.30 |
| 1 | I | 315 | LEU | CA-CB-CG | 7.12 | 131.67 | 115.30 |
| 1 | E | 252 | ASP | CB-CG-OD1 | 6.78 | 124.40 | 118.30 |
| 1 | H | 791 | LEU | CA-CB-CG | 6.67 | 130.63 | 115.30 |
| 1 | L | 315 | LEU | CA-CB-CG | 6.34 | 129.89 | 115.30 |
| 1 | J | 252 | ASP | CB-CG-OD1 | 6.18 | 123.87 | 118.30 |
| 1 | K | 252 | ASP | CB-CG-OD1 | 6.11 | 123.80 | 118.30 |
| 1 | A | 252 | ASP | CB-CG-OD1 | 6.09 | 123.79 | 118.30 |
| 1 | A | 790 | LEU | CA-CB-CG | 5.94 | 128.96 | 115.30 |
| 1 | H | 252 | ASP | CB-CG-OD1 | 5.92 | 123.63 | 118.30 |
| 1 | C | 900 | LEU | CA-CB-CG | 5.86 | 128.77 | 115.30 |
| 1 | B | 773 | MET | C-N-CD | -5.74 | 107.98 | 120.60 |
| 1 | E | 696 | ILE | C-N-CA | 5.59 | 135.69 | 121.70 |
| 1 | B | 315 | LEU | CA-CB-CG | 5.58 | 128.13 | 115.30 |
| 1 | I | 696 | ILE | C-N-CA | 5.58 | 135.64 | 121.70 |
| 1 | L | 876 | LEU | CA-CB-CG | 5.57 | 128.11 | 115.30 |
| 1 | A | 696 | ILE | C-N-CA | 5.48 | 135.40 | 121.70 |
| 1 | B | 876 | LEU | CA-CB-CG | 5.46 | 127.87 | 115.30 |
| 1 | F | 876 | LEU | CA-CB-CG | 5.46 | 127.85 | 115.30 |
| 1 | H | 696 | ILE | C-N-CA | 5.40 | 135.20 | 121.70 |
| 1 | I | 898 | LEU | CA-CB-CG | 5.39 | 127.69 | 115.30 |
| 5 | M | 239 | LEU | CA-CB-CG | 5.38 | 127.68 | 115.30 |
| 1 | L | 696 | ILE | C-N-CA | 5.37 | 135.12 | 121.70 |
| 1 | I | 91 | LEU | CA-CB-CG | 5.35 | 127.60 | 115.30 |
| 1 | K | 696 | ILE | C-N-CA | 5.33 | 135.03 | 121.70 |
| 1 | F | 696 | ILE | C-N-CA | 5.32 | 134.99 | 121.70 |
| 1 | A | 873 | LEU | CA-CB-CG | 5.19 | 127.24 | 115.30 |
| 1 | B | 696 | ILE | C-N-CA | 5.17 | 134.63 | 121.70 |

*Continued on next page...*

CONFIDENTIAL VALIDATION REPORT

*Continued from previous page...*

| Mol | Chain | Res | Type | Atoms | Z | Observed($^o$) | Ideal($^o$) |
|---|---|---|---|---|---|---|---|
| 1 | J | 521 | LEU | CA-CB-CG | 5.16 | 127.17 | 115.30 |
| 1 | F | 682 | ASP | CB-CG-OD1 | 5.16 | 122.94 | 118.30 |
| 1 | I | 241 | MET | CA-CB-CG | 5.15 | 122.06 | 113.30 |
| 1 | I | 353 | TYR | CA-CB-CG | 5.11 | 123.10 | 113.40 |
| 1 | F | 368 | SER | C-N-CA | 5.10 | 134.46 | 121.70 |
| 1 | B | 673 | ASP | CB-CG-OD1 | 5.05 | 122.84 | 118.30 |
| 1 | L | 682 | ASP | CB-CG-OD1 | 5.04 | 122.84 | 118.30 |

There are no chirality outliers.

All (37) planarity outliers are listed below:

| Mol | Chain | Res | Type | Group |
|---|---|---|---|---|
| 1 | A | 260 | ASP | Peptide |
| 1 | A | 261 | ASP | Peptide |
| 1 | A | 298 | GLN | Peptide |
| 1 | A | 791 | LEU | Peptide |
| 1 | B | 178 | PRO | Peptide |
| 1 | B | 299 | THR | Peptide |
| 1 | B | 697 | ASN | Peptide |
| 1 | B | 772 | THR | Peptide |
| 1 | C | 365 | HIS | Peptide |
| 1 | D | 345 | VAL | Peptide |
| 1 | D | 346 | ARG | Peptide |
| 1 | D | 365 | HIS | Peptide |
| 1 | E | 298 | GLN | Peptide |
| 1 | E | 3 | PRO | Peptide |
| 1 | E | 365 | HIS | Peptide |
| 1 | G | 345 | VAL | Peptide |
| 1 | G | 346 | ARG | Peptide |
| 1 | G | 365 | HIS | Peptide |
| 1 | G | 791 | LEU | Peptide |
| 1 | H | 261 | ASP | Peptide |
| 1 | H | 365 | HIS | Peptide |
| 1 | H | 701 | GLU | Peptide |
| 1 | I | 345 | VAL | Peptide |
| 1 | I | 346 | ARG | Peptide |
| 1 | J | 299 | THR | Peptide |
| 1 | J | 346 | ARG | Peptide |
| 1 | J | 393 | ALA | Peptide |
| 1 | K | 146 | TYR | Peptide |
| 1 | K | 365 | HIS | Peptide |
| 1 | L | 172 | GLN | Peptide |

*Continued on next page...*

CONFIDENTIAL VALIDATION REPORT

*Continued from previous page...*

| Mol | Chain | Res | Type | Group |
|---|---|---|---|---|
| 1 | L | 345 | VAL | Peptide |
| 1 | L | 346 | ARG | Peptide |
| 1 | L | 697 | ASN | Peptide |
| 1 | L | 884 | VAL | Peptide |
| 5 | M | 118 | ASP | Peptide |
| 5 | M | 119 | PRO | Peptide |
| 5 | M | 40 | LYS | Peptide |

#### 5.2 Too-close contacts ⓘ

In the following table, the Non-H and H(model) columns list the number of non-hydrogen atoms and hydrogen atoms in the chain respectively. The H(added) column lists the number of hydrogen atoms added and optimized by MolProbity. The Clashes column lists the number of clashes within the asymmetric unit, whereas Symm-Clashes lists symmetry related clashes.

| Mol | Chain | Non-H | H(model) | H(added) | Clashes | Symm-Clashes |
|---|---|---|---|---|---|---|
| 1 | A | 7168 | 0 | 6800 | 100 | 0 |
| 1 | B | 7161 | 0 | 6790 | 115 | 0 |
| 1 | C | 7179 | 0 | 6810 | 102 | 0 |
| 1 | D | 7174 | 0 | 6805 | 107 | 0 |
| 1 | E | 7171 | 0 | 6798 | 103 | 0 |
| 1 | F | 7172 | 0 | 6800 | 90 | 0 |
| 1 | G | 7171 | 0 | 6798 | 91 | 0 |
| 1 | H | 7166 | 0 | 6793 | 104 | 0 |
| 1 | I | 7171 | 0 | 6798 | 105 | 0 |
| 1 | J | 7171 | 0 | 6798 | 85 | 0 |
| 1 | K | 7184 | 0 | 6815 | 103 | 0 |
| 1 | L | 7174 | 0 | 6805 | 96 | 0 |
| 2 | Q | 2874 | 0 | 2742 | 38 | 0 |
| 2 | R | 2874 | 0 | 2742 | 44 | 0 |
| 2 | S | 2874 | 0 | 2742 | 38 | 0 |
| 2 | T | 2874 | 0 | 2742 | 26 | 0 |
| 3 | O | 1452 | 0 | 1400 | 11 | 0 |
| 3 | P | 1452 | 0 | 1400 | 12 | 0 |
| 4 | N | 1980 | 0 | 2014 | 15 | 0 |
| 5 | M | 3567 | 0 | 3505 | 53 | 0 |
| All | All | 106009 | 0 | 100897 | 1180 | 0 |

The all-atom clashscore is defined as the number of clashes found per 1000 atoms (including hydrogen atoms). The all-atom clashscore for this structure is 6.

All (1180) close contacts within the same asymmetric unit are listed below, sorted by their clash magnitude.

CONFIDENTIAL VALIDATION REPORT

| Atom-1 | Atom-2 | Interatomic distance (Å) | Clash overlap (Å) |
|---|---|---|---|
| 1:H:312:ASN:HB2 | 1:H:598:ASN:HD21 | 1.48 | 0.78 |
| 1:B:50:GLY:HA2 | 1:B:570:ASN:HD22 | 1.56 | 0.71 |
| 4:N:186:ASN:HB3 | 4:N:218:ASN:HD21 | 1.56 | 0.70 |
| 1:F:50:GLY:HA2 | 1:F:570:ASN:HD22 | 1.55 | 0.70 |
| 1:K:246:ILE:HG22 | 1:K:248:TRP:H | 1.57 | 0.70 |
| 1:K:662:SER:O | 1:K:828:TRP:NE1 | 2.25 | 0.69 |
| 1:I:143:PRO:HB2 | 1:I:171:GLY:HA2 | 1.73 | 0.69 |
| 1:D:771:GLN:HG3 | 2:S:54:GLN:HG2 | 1.75 | 0.69 |
| 2:R:218:ASN:HD21 | 2:Q:161:TRP:HD1 | 1.41 | 0.69 |
| 1:G:327:ARG:HB3 | 1:G:338:VAL:HG11 | 1.75 | 0.69 |
| 1:F:705:THR:HG23 | 1:F:712:LYS:HG2 | 1.76 | 0.68 |
| 3:P:23:ARG:HH12 | 3:P:259:PRO:HD3 | 1.58 | 0.68 |
| 3:O:23:ARG:HH12 | 3:O:259:PRO:HD3 | 1.58 | 0.68 |
| 1:J:160:GLN:HB3 | 2:Q:91:THR:HG22 | 1.75 | 0.68 |
| 1:H:149:GLY:HA2 | 1:H:185:ASN:HA | 1.76 | 0.68 |
| 1:L:352:GLY:H | 1:J:111:TYR:HB2 | 1.60 | 0.67 |
| 1:D:50:GLY:HA2 | 1:D:570:ASN:HD22 | 1.58 | 0.67 |
| 1:E:662:SER:O | 1:E:828:TRP:NE1 | 2.26 | 0.67 |
| 1:H:662:SER:O | 1:H:828:TRP:NE1 | 2.27 | 0.67 |
| 1:E:246:ILE:HG22 | 1:E:248:TRP:H | 1.60 | 0.67 |
| 1:B:5:ARG:HG3 | 1:B:10:ILE:HB | 1.77 | 0.67 |
| 1:L:666:GLN:NE2 | 1:L:865:ASP:OD1 | 2.28 | 0.66 |
| 1:I:246:ILE:HG22 | 1:I:248:TRP:H | 1.61 | 0.66 |
| 1:A:520:LEU:HB3 | 1:A:588:GLN:HE22 | 1.61 | 0.66 |
| 1:J:143:PRO:HB2 | 1:J:171:GLY:HA2 | 1.78 | 0.66 |
| 1:I:349:GLU:HG2 | 1:I:470:ARG:HG2 | 1.78 | 0.66 |
| 1:D:246:ILE:HG22 | 1:D:248:TRP:H | 1.61 | 0.65 |
| 1:C:287:ASN:HD21 | 1:C:313:SER:H | 1.45 | 0.65 |
| 1:L:327:ARG:HB3 | 1:L:338:VAL:HG21 | 1.79 | 0.65 |
| 1:C:143:PRO:HB2 | 1:C:171:GLY:HA2 | 1.77 | 0.65 |
| 1:A:150:GLY:H | 1:A:236:ARG:HH12 | 1.45 | 0.65 |
| 1:F:797:PRO:HD3 | 1:E:404:ILE:HD11 | 1.79 | 0.65 |
| 1:E:50:GLY:HA2 | 1:E:570:ASN:HD22 | 1.61 | 0.64 |
| 1:L:50:GLY:HA2 | 1:L:570:ASN:HD22 | 1.61 | 0.64 |
| 1:I:520:LEU:HB3 | 1:I:588:GLN:HE22 | 1.62 | 0.64 |
| 1:D:294:SER:HB3 | 1:D:529:GLU:HB3 | 1.80 | 0.64 |
| 1:G:626:SER:HB3 | 1:G:876:LEU:HB2 | 1.79 | 0.64 |
| 1:I:140:ALA:HB3 | 1:G:399:ILE:HG22 | 1.80 | 0.64 |
| 1:B:104:ARG:NH2 | 1:B:500:GLY:O | 2.31 | 0.63 |
| 1:B:705:THR:HG23 | 1:B:712:LYS:HG2 | 1.81 | 0.63 |
| 1:B:666:GLN:NE2 | 1:B:865:ASP:OD1 | 2.32 | 0.63 |
| 1:K:725:GLN:NE2 | 1:K:731:ASN:OD1 | 2.31 | 0.63 |

*Continued on next page...*

CONFIDENTIAL VALIDATION REPORT

*Continued from previous page...*

| Atom-1 | Atom-2 | Interatomic distance (Å) | Clash overlap (Å) |
|---|---|---|---|
| 1:K:626:SER:HB2 | 1:K:876:LEU:HB2 | 1.80 | 0.62 |
| 1:L:140:ALA:HB3 | 1:J:399:ILE:HG22 | 1.81 | 0.62 |
| 1:L:520:LEU:HB3 | 1:L:588:GLN:HE22 | 1.64 | 0.62 |
| 1:G:318:GLN:HG3 | 1:G:656:ASP:HA | 1.81 | 0.62 |
| 4:N:139:VAL:HA | 4:N:144:GLN:HE22 | 1.64 | 0.62 |
| 1:B:345:VAL:HG11 | 1:B:484:HIS:HE2 | 1.65 | 0.62 |
| 1:A:797:PRO:HD3 | 1:C:404:ILE:HD11 | 1.81 | 0.61 |
| 1:D:287:ASN:HD21 | 1:D:313:SER:H | 1.48 | 0.61 |
| 1:J:890:ARG:HD2 | 3:P:120:GLY:HA3 | 1.82 | 0.61 |
| 1:B:696:ILE:HD11 | 1:B:818:GLU:HA | 1.82 | 0.61 |
| 1:A:5:ARG:NH2 | 1:C:850:MET:SD | 2.67 | 0.61 |
| 1:E:511:GLN:NE2 | 1:E:528:TYR:OH | 2.33 | 0.61 |
| 1:H:797:PRO:HD3 | 1:G:404:ILE:HD11 | 1.82 | 0.61 |
| 1:B:724:ASN:HD22 | 1:B:837:LEU:HB2 | 1.66 | 0.61 |
| 1:F:10:ILE:HG22 | 1:E:842:LEU:HD12 | 1.81 | 0.61 |
| 1:I:104:ARG:NH2 | 1:I:500:GLY:O | 2.33 | 0.61 |
| 1:H:589:ASN:HB2 | 1:I:41:ARG:H | 1.64 | 0.61 |
| 1:L:104:ARG:NH2 | 1:L:500:GLY:O | 2.34 | 0.61 |
| 1:B:287:ASN:HD21 | 1:B:313:SER:H | 1.48 | 0.61 |
| 1:B:353:TYR:HB2 | 1:B:412:PRO:HB2 | 1.81 | 0.61 |
| 1:I:725:GLN:NE2 | 1:I:731:ASN:OD1 | 2.33 | 0.61 |
| 2:S:218:ASN:HD21 | 2:R:161:TRP:HD1 | 1.46 | 0.61 |
| 1:F:399:ILE:HG22 | 1:E:140:ALA:HB3 | 1.83 | 0.61 |
| 1:I:879:VAL:HG12 | 1:I:901:PRO:HD2 | 1.83 | 0.60 |
| 1:E:197:SER:HG | 1:E:248:TRP:HE1 | 1.50 | 0.60 |
| 1:I:696:ILE:HD11 | 1:I:818:GLU:HA | 1.83 | 0.60 |
| 1:F:404:ILE:HD11 | 1:D:797:PRO:HD3 | 1.82 | 0.60 |
| 1:H:398:LEU:HB2 | 1:G:139:VAL:HG22 | 1.84 | 0.60 |
| 1:K:143:PRO:O | 1:K:172:GLN:NE2 | 2.35 | 0.60 |
| 1:G:637:ARG:NH1 | 1:G:872:TYR:OH | 2.34 | 0.60 |
| 1:J:626:SER:HB2 | 1:J:876:LEU:HB2 | 1.84 | 0.60 |
| 1:A:662:SER:O | 1:A:828:TRP:NE1 | 2.26 | 0.60 |
| 1:B:284:MET:HB3 | 1:B:530:TRP:HE1 | 1.67 | 0.60 |
| 1:H:246:ILE:HG22 | 1:H:248:TRP:H | 1.65 | 0.60 |
| 1:J:246:ILE:HG22 | 1:J:248:TRP:H | 1.66 | 0.60 |
| 1:H:205:TYR:HH | 1:G:353:TYR:HH | 1.49 | 0.60 |
| 1:L:127:ASN:OD1 | 1:K:796:HIS:ND1 | 2.34 | 0.60 |
| 5:M:200:ASP:HB2 | 5:M:286:TRP:HB3 | 1.83 | 0.60 |
| 1:A:278:ASP:OD2 | 1:A:327:ARG:NH2 | 2.32 | 0.60 |
| 1:G:417:ARG:NH2 | 1:G:782:GLN:OE1 | 2.35 | 0.60 |
| 1:L:676:VAL:HG22 | 1:I:905:GLY:H | 1.67 | 0.60 |

*Continued on next page...*

CONFIDENTIAL VALIDATION REPORT

*Continued from previous page...*

| Atom-1 | Atom-2 | Interatomic distance (Å) | Clash overlap (Å) |
|---|---|---|---|
| 1:L:682:ASP:HA | 1:L:687:PRO:HB3 | 1.83 | 0.60 |
| 1:G:104:ARG:NH2 | 1:G:500:GLY:O | 2.35 | 0.60 |
| 2:R:171:ARG:HH22 | 2:Q:135:THR:HG1 | 1.48 | 0.60 |
| 1:B:725:GLN:NE2 | 1:B:731:ASN:OD1 | 2.35 | 0.60 |
| 1:E:346:ARG:HD3 | 1:E:481:PRO:HB3 | 1.83 | 0.60 |
| 1:L:346:ARG:HH21 | 1:L:822:LEU:HB3 | 1.67 | 0.60 |
| 1:K:287:ASN:ND2 | 1:K:307:ASP:OD1 | 2.35 | 0.59 |
| 2:S:161:TRP:HD1 | 2:Q:218:ASN:HD21 | 1.47 | 0.59 |
| 1:C:246:ILE:HG22 | 1:C:248:TRP:H | 1.66 | 0.59 |
| 1:E:904:SER:HA | 1:C:676:VAL:HG22 | 1.84 | 0.59 |
| 1:E:350:ASN:OD1 | 1:E:416:LYS:NZ | 2.34 | 0.59 |
| 1:G:337:ALA:O | 1:G:493:ARG:NH1 | 2.35 | 0.59 |
| 1:G:310:ASP:HB3 | 1:G:597:VAL:HG13 | 1.85 | 0.59 |
| 1:H:170:VAL:HG22 | 1:G:784:LYS:HE2 | 1.84 | 0.59 |
| 1:H:97:ASP:HB3 | 1:H:561:ASN:HB2 | 1.84 | 0.59 |
| 1:K:563:TYR:HE2 | 1:J:712:LYS:HZ3 | 1.50 | 0.59 |
| 1:A:796:HIS:ND1 | 1:B:127:ASN:OD1 | 2.35 | 0.59 |
| 1:H:783:GLN:NE2 | 1:H:792:GLU:O | 2.36 | 0.59 |
| 1:K:170:VAL:HG22 | 1:J:784:LYS:HE2 | 1.84 | 0.59 |
| 1:C:310:ASP:HB3 | 1:C:597:VAL:HG13 | 1.85 | 0.59 |
| 1:F:140:ALA:HB3 | 1:D:399:ILE:HG22 | 1.83 | 0.59 |
| 1:H:626:SER:HB2 | 1:H:876:LEU:HB2 | 1.83 | 0.59 |
| 1:F:118:PRO:O | 1:E:410:ASN:ND2 | 2.36 | 0.59 |
| 1:J:104:ARG:NH2 | 1:J:500:GLY:O | 2.36 | 0.59 |
| 1:L:797:PRO:HD3 | 1:K:404:ILE:HD11 | 1.82 | 0.59 |
| 1:K:792:GLU:OE1 | 1:K:793:ARG:NH2 | 2.36 | 0.59 |
| 1:L:724:ASN:HD22 | 1:L:837:LEU:HB2 | 1.67 | 0.59 |
| 1:C:99:LYS:HD3 | 1:C:559:GLN:HE21 | 1.68 | 0.59 |
| 1:F:403:ASN:ND2 | 1:D:794:CYS:O | 2.34 | 0.59 |
| 1:L:337:ALA:O | 1:L:493:ARG:NH1 | 2.36 | 0.59 |
| 1:A:842:LEU:HD12 | 1:B:10:ILE:HG22 | 1.84 | 0.59 |
| 1:B:310:ASP:HB3 | 1:B:597:VAL:HG13 | 1.85 | 0.59 |
| 1:F:696:ILE:HD11 | 1:F:818:GLU:HA | 1.84 | 0.59 |
| 1:G:139:VAL:HG11 | 1:G:262:ARG:HH12 | 1.68 | 0.59 |
| 1:K:106:PRO:HD2 | 1:K:551:ASP:HB2 | 1.84 | 0.59 |
| 1:L:631:LYS:NZ | 1:L:869:GLU:OE2 | 2.36 | 0.59 |
| 1:A:410:ASN:ND2 | 1:B:118:PRO:O | 2.36 | 0.58 |
| 1:A:301:GLN:HB2 | 1:L:61:LEU:HD22 | 1.83 | 0.58 |
| 1:F:617:ARG:NH2 | 1:F:901:PRO:O | 2.36 | 0.58 |
| 1:C:70:ASP:OD1 | 1:C:534:LYS:NZ | 2.35 | 0.58 |
| 1:F:778:ASN:HB3 | 1:D:118:PRO:HD2 | 1.84 | 0.58 |

*Continued on next page...*

CONFIDENTIAL VALIDATION REPORT

*Continued from previous page...*

| Atom-1 | Atom-2 | Interatomic distance (Å) | Clash overlap (Å) |
|---|---|---|---|
| 1:H:563:TYR:HE2 | 1:G:712:LYS:HZ3 | 1.51 | 0.58 |
| 1:K:144:GLN:HB2 | 1:K:189:LEU:HB3 | 1.86 | 0.58 |
| 1:A:626:SER:HB2 | 1:A:876:LEU:HB2 | 1.86 | 0.58 |
| 1:C:399:ILE:HG22 | 1:B:140:ALA:HB3 | 1.86 | 0.58 |
| 1:H:519:LEU:HD11 | 1:H:885:ILE:HD13 | 1.85 | 0.58 |
| 1:K:346:ARG:HD3 | 1:K:481:PRO:HB3 | 1.86 | 0.58 |
| 1:L:885:ILE:HG12 | 1:L:895:VAL:HG12 | 1.83 | 0.58 |
| 1:I:694:ARG:NH1 | 1:I:699:ASP:OD2 | 2.37 | 0.58 |
| 4:N:63:LEU:HD12 | 4:N:77:VAL:HG12 | 1.86 | 0.58 |
| 3:O:255:LYS:NZ | 3:O:267:VAL:O | 2.37 | 0.58 |
| 1:B:16:LYS:O | 3:O:221:ARG:NH1 | 2.37 | 0.58 |
| 1:C:104:ARG:NH2 | 1:C:500:GLY:O | 2.37 | 0.58 |
| 1:L:725:GLN:NE2 | 1:L:731:ASN:OD1 | 2.35 | 0.58 |
| 1:K:186:ASN:O | 1:K:236:ARG:NH2 | 2.36 | 0.58 |
| 1:B:277:ARG:HH12 | 1:B:652:ILE:HD12 | 1.68 | 0.58 |
| 1:C:327:ARG:HB3 | 1:C:338:VAL:HG11 | 1.85 | 0.58 |
| 1:F:284:MET:HB3 | 1:F:530:TRP:HE1 | 1.69 | 0.58 |
| 1:I:89:VAL:HB | 1:I:521:LEU:HB3 | 1.84 | 0.58 |
| 1:J:284:MET:HB3 | 1:J:530:TRP:HE1 | 1.69 | 0.58 |
| 1:L:705:THR:HG23 | 1:L:712:LYS:HG2 | 1.85 | 0.58 |
| 3:P:255:LYS:NZ | 3:P:267:VAL:O | 2.37 | 0.58 |
| 2:T:49:LEU:HD12 | 2:T:70:GLY:HA2 | 1.86 | 0.58 |
| 1:A:346:ARG:HD3 | 1:A:481:PRO:HB3 | 1.86 | 0.57 |
| 1:L:277:ARG:HH12 | 1:L:652:ILE:HD12 | 1.69 | 0.57 |
| 1:A:104:ARG:NH2 | 1:A:500:GLY:O | 2.37 | 0.57 |
| 1:E:104:ARG:NH2 | 1:E:500:GLY:O | 2.37 | 0.57 |
| 1:E:626:SER:HB2 | 1:E:876:LEU:HB2 | 1.84 | 0.57 |
| 1:K:737:LYS:O | 1:K:743:ASN:ND2 | 2.37 | 0.57 |
| 5:M:183:LEU:O | 5:M:278:LYS:NZ | 2.37 | 0.57 |
| 1:B:121:PRO:HG2 | 1:B:124:ALA:HB2 | 1.85 | 0.57 |
| 1:E:109:LYS:HE2 | 1:E:252:ASP:HB2 | 1.86 | 0.57 |
| 2:Q:49:LEU:HD12 | 2:Q:70:GLY:HA2 | 1.86 | 0.57 |
| 1:A:117:ASN:HD22 | 1:C:778:ASN:HA | 1.70 | 0.57 |
| 1:F:379:GLY:HA2 | 1:E:228:ILE:HA | 1.86 | 0.57 |
| 5:M:390:ASP:OD1 | 5:M:393:GLN:NE2 | 2.37 | 0.57 |
| 1:A:778:ASN:HB3 | 1:B:118:PRO:HD2 | 1.86 | 0.57 |
| 1:L:778:ASN:HB3 | 1:J:118:PRO:HD2 | 1.86 | 0.57 |
| 3:O:96:THR:HG22 | 3:O:99:ARG:HD3 | 1.86 | 0.57 |
| 2:S:49:LEU:HD12 | 2:S:70:GLY:HA2 | 1.86 | 0.57 |
| 1:H:254:HIS:NE2 | 1:H:452:ASN:O | 2.34 | 0.57 |
| 1:H:694:ARG:NH2 | 1:H:702:GLY:O | 2.38 | 0.57 |

*Continued on next page...*

CONFIDENTIAL VALIDATION REPORT

*Continued from previous page...*

| Atom-1 | Atom-2 | Interatomic distance (Å) | Clash overlap (Å) |
|---|---|---|---|
| 1:J:192:ILE:HG12 | 1:J:241:MET:HB3 | 1.87 | 0.57 |
| 1:J:277:ARG:HH12 | 1:J:652:ILE:HD12 | 1.70 | 0.57 |
| 1:K:50:GLY:HA2 | 1:K:570:ASN:HD22 | 1.69 | 0.57 |
| 3:P:96:THR:HG22 | 3:P:99:ARG:HD3 | 1.86 | 0.57 |
| 1:C:559:GLN:NE2 | 1:B:700:ALA:O | 2.34 | 0.56 |
| 1:E:417:ARG:NH2 | 1:E:782:GLN:OE1 | 2.38 | 0.56 |
| 1:G:611:VAL:HG22 | 1:G:861:GLN:HG2 | 1.87 | 0.56 |
| 1:G:694:ARG:NH2 | 1:G:698:GLN:O | 2.38 | 0.56 |
| 1:H:283:MET:HG2 | 1:H:509:VAL:HG11 | 1.86 | 0.56 |
| 1:K:16:LYS:O | 3:P:221:ARG:NH1 | 2.37 | 0.56 |
| 5:M:243:ASN:HB2 | 5:M:280:GLN:HE21 | 1.69 | 0.56 |
| 1:B:127:ASN:HB3 | 1:B:192:ILE:HD13 | 1.86 | 0.56 |
| 1:E:283:MET:HG2 | 1:E:509:VAL:HG11 | 1.87 | 0.56 |
| 1:J:111:TYR:OH | 1:J:252:ASP:OD2 | 2.21 | 0.56 |
| 5:M:93:LYS:HB2 | 5:M:435:LYS:HB3 | 1.86 | 0.56 |
| 1:A:100:GLY:HA2 | 1:A:558:THR:HG22 | 1.86 | 0.56 |
| 3:O:79:GLN:HE22 | 3:O:224:ARG:H | 1.52 | 0.56 |
| 3:P:79:GLN:HE22 | 3:P:224:ARG:H | 1.52 | 0.56 |
| 2:R:49:LEU:HD12 | 2:R:70:GLY:HA2 | 1.86 | 0.56 |
| 1:A:346:ARG:NH1 | 1:A:476:MET:O | 2.39 | 0.56 |
| 1:F:700:ALA:O | 1:D:559:GLN:NE2 | 2.38 | 0.56 |
| 1:G:291:ASN:HD21 | 1:G:641:THR:HG22 | 1.71 | 0.56 |
| 1:H:353:TYR:HH | 1:I:205:TYR:HH | 1.53 | 0.56 |
| 1:I:256:ALA:HB3 | 1:I:265:THR:HG23 | 1.88 | 0.56 |
| 1:C:50:GLY:HA2 | 1:C:570:ASN:HD22 | 1.70 | 0.56 |
| 1:E:310:ASP:HB3 | 1:E:597:VAL:HG13 | 1.87 | 0.56 |
| 1:F:887:GLN:HG3 | 1:F:893:ILE:HG22 | 1.88 | 0.56 |
| 1:J:843:THR:HG23 | 1:J:846:GLY:H | 1.70 | 0.56 |
| 1:L:85:ASP:OD1 | 1:L:890:ARG:NH1 | 2.38 | 0.56 |
| 1:A:404:ILE:HD11 | 1:B:797:PRO:HD3 | 1.88 | 0.56 |
| 1:H:549:ARG:HH12 | 1:H:644:PRO:HA | 1.71 | 0.56 |
| 1:K:346:ARG:NH1 | 1:K:476:MET:O | 2.39 | 0.56 |
| 1:A:244:ASP:OD2 | 1:C:798:TYR:OH | 2.24 | 0.56 |
| 1:I:337:ALA:O | 1:I:493:ARG:NH1 | 2.39 | 0.56 |
| 1:I:277:ARG:HH12 | 1:I:652:ILE:HD12 | 1.71 | 0.56 |
| 1:A:470:ARG:NH2 | 1:A:746:GLU:OE1 | 2.39 | 0.56 |
| 1:D:626:SER:HB2 | 1:D:876:LEU:HB2 | 1.87 | 0.56 |
| 1:H:778:ASN:HA | 1:I:117:ASN:HD22 | 1.71 | 0.56 |
| 1:I:70:ASP:OD1 | 1:I:534:LYS:NZ | 2.38 | 0.56 |
| 1:A:904:SER:O | 1:K:861:GLN:NE2 | 2.38 | 0.56 |
| 1:D:613:ASN:HD21 | 2:R:11:ASN:HD21 | 1.52 | 0.56 |

*Continued on next page...*

CONFIDENTIAL VALIDATION REPORT

*Continued from previous page...*

| Atom-1 | Atom-2 | Interatomic distance (Å) | Clash overlap (Å) |
|---|---|---|---|
| 1:E:504:ASP:H | 1:D:817:THR:HG21 | 1.71 | 0.55 |
| 1:G:246:ILE:HG22 | 1:G:248:TRP:H | 1.70 | 0.55 |
| 1:A:879:VAL:HG12 | 1:A:901:PRO:HD2 | 1.88 | 0.55 |
| 1:C:718:GLN:NE2 | 1:C:829:GLN:O | 2.39 | 0.55 |
| 1:E:749:SER:OG | 1:E:819:ARG:NH1 | 2.39 | 0.55 |
| 1:K:533:ARG:HG2 | 1:K:535:ASP:H | 1.71 | 0.55 |
| 1:B:450:TYR:OH | 1:B:454:ARG:NH1 | 2.39 | 0.55 |
| 1:F:350:ASN:HB3 | 1:F:467:ILE:HA | 1.89 | 0.55 |
| 1:H:879:VAL:HG12 | 1:H:901:PRO:HD2 | 1.87 | 0.55 |
| 1:I:284:MET:HB3 | 1:I:530:TRP:HE1 | 1.71 | 0.55 |
| 1:J:310:ASP:HB3 | 1:J:597:VAL:HG13 | 1.87 | 0.55 |
| 2:S:97:ARG:NH1 | 2:Q:100:ASP:OD1 | 2.33 | 0.55 |
| 1:C:532:PHE:HE2 | 1:C:560:ILE:HG21 | 1.72 | 0.55 |
| 1:E:150:GLY:O | 1:E:236:ARG:NH1 | 2.39 | 0.55 |
| 1:E:244:ASP:OD2 | 1:D:798:TYR:OH | 2.23 | 0.55 |
| 1:A:794:CYS:HB2 | 1:B:171:GLY:HA2 | 1.88 | 0.55 |
| 1:C:169:GLN:HE21 | 1:B:773:MET:HG3 | 1.71 | 0.55 |
| 1:K:495:GLN:NE2 | 1:J:351:ASP:OD1 | 2.40 | 0.55 |
| 5:M:168:GLY:HA2 | 5:M:204:LEU:HD12 | 1.87 | 0.55 |
| 4:N:194:LEU:O | 4:N:198:TRP:NE1 | 2.39 | 0.55 |
| 1:C:117:ASN:HD22 | 1:B:778:ASN:HA | 1.72 | 0.55 |
| 1:E:666:GLN:NE2 | 1:E:865:ASP:OD1 | 2.40 | 0.55 |
| 1:H:140:ALA:HB3 | 1:I:399:ILE:HG22 | 1.87 | 0.55 |
| 1:J:885:ILE:HG12 | 1:J:895:VAL:HG12 | 1.88 | 0.55 |
| 1:K:117:ASN:HD22 | 1:J:778:ASN:HA | 1.72 | 0.55 |
| 1:L:411:LEU:HD21 | 1:J:415:LEU:HD12 | 1.89 | 0.55 |
| 4:N:85:MET:O | 4:N:89:HIS:ND1 | 2.38 | 0.55 |
| 1:A:143:PRO:O | 1:A:172:GLN:NE2 | 2.40 | 0.55 |
| 1:F:98:ILE:HG22 | 1:F:560:ILE:HG23 | 1.88 | 0.55 |
| 1:H:798:TYR:OH | 1:I:244:ASP:OD2 | 2.25 | 0.55 |
| 1:I:640:ALA:HB2 | 2:Q:17:PRO:HD3 | 1.89 | 0.55 |
| 1:K:278:ASP:OD2 | 1:K:327:ARG:NH2 | 2.39 | 0.55 |
| 1:L:346:ARG:HE | 1:L:822:LEU:HD13 | 1.71 | 0.55 |
| 1:L:57:GLN:HB2 | 1:L:87:ARG:HH12 | 1.72 | 0.55 |
| 2:R:100:ASP:OD1 | 2:Q:97:ARG:NH1 | 2.32 | 0.55 |
| 1:L:563:TYR:HE2 | 1:K:712:LYS:HZ2 | 1.55 | 0.55 |
| 1:C:256:ALA:O | 1:C:788:ASN:ND2 | 2.40 | 0.55 |
| 1:F:701:GLU:OE2 | 1:D:561:ASN:ND2 | 2.40 | 0.55 |
| 1:E:735:CYS:SG | 1:E:737:LYS:NZ | 2.76 | 0.55 |
| 1:I:724:ASN:HD22 | 1:I:837:LEU:HB2 | 1.71 | 0.55 |
| 1:K:104:ARG:NH2 | 1:K:500:GLY:O | 2.40 | 0.55 |

*Continued on next page...*

CONFIDENTIAL VALIDATION REPORT

*Continued from previous page...*

| Atom-1 | Atom-2 | Interatomic distance (Å) | Clash overlap (Å) |
|---|---|---|---|
| 1:K:100:GLY:HA2 | 1:K:558:THR:HG22 | 1.87 | 0.55 |
| 1:L:97:ASP:OD1 | 1:K:712:LYS:NZ | 2.40 | 0.55 |
| 1:K:5:ARG:HG3 | 3:P:15:GLN:HB2 | 1.89 | 0.55 |
| 1:H:104:ARG:NH2 | 1:H:500:GLY:O | 2.41 | 0.54 |
| 1:H:150:GLY:O | 1:H:236:ARG:NH1 | 2.41 | 0.54 |
| 1:H:346:ARG:HD3 | 1:H:481:PRO:HB3 | 1.89 | 0.54 |
| 1:H:52:THR:HG21 | 1:G:834:SER:HB3 | 1.89 | 0.54 |
| 1:H:520:LEU:HB3 | 1:H:588:GLN:HE22 | 1.71 | 0.54 |
| 1:D:337:ALA:O | 1:D:493:ARG:NH1 | 2.40 | 0.54 |
| 1:E:169:GLN:NE2 | 1:D:775:ILE:O | 2.40 | 0.54 |
| 1:G:460:ILE:HD12 | 1:G:461:VAL:HG23 | 1.89 | 0.54 |
| 1:H:118:PRO:HD2 | 1:G:778:ASN:HB3 | 1.89 | 0.54 |
| 1:D:13:ARG:HB3 | 1:D:17:GLU:HG3 | 1.89 | 0.54 |
| 1:E:549:ARG:NH1 | 1:E:646:PHE:O | 2.40 | 0.54 |
| 1:H:101:THR:HA | 1:H:504:ASP:HA | 1.89 | 0.54 |
| 1:I:327:ARG:HH11 | 1:I:338:VAL:HG21 | 1.71 | 0.54 |
| 1:J:50:GLY:HA2 | 1:J:570:ASN:HD22 | 1.72 | 0.54 |
| 1:L:118:PRO:O | 1:K:410:ASN:ND2 | 2.40 | 0.54 |
| 1:H:399:ILE:HG13 | 1:G:228:ILE:HD13 | 1.89 | 0.54 |
| 1:J:337:ALA:O | 1:J:493:ARG:NH1 | 2.40 | 0.54 |
| 1:C:121:PRO:HG2 | 1:C:124:ALA:HB2 | 1.89 | 0.54 |
| 1:C:75:TYR:HB2 | 1:C:534:LYS:HE3 | 1.89 | 0.54 |
| 1:C:98:ILE:HG12 | 1:C:560:ILE:HG12 | 1.88 | 0.54 |
| 1:D:460:ILE:HD12 | 1:D:461:VAL:HG23 | 1.89 | 0.54 |
| 1:L:696:ILE:HD11 | 1:L:818:GLU:HA | 1.87 | 0.54 |
| 1:L:798:TYR:OH | 1:J:244:ASP:OD2 | 2.25 | 0.54 |
| 1:L:131:THR:HA | 1:L:136:THR:HA | 1.88 | 0.54 |
| 1:A:563:TYR:HE2 | 1:C:712:LYS:HZ3 | 1.54 | 0.54 |
| 1:E:309:ASN:OD1 | 1:C:688:ASN:ND2 | 2.39 | 0.54 |
| 1:I:645:ASN:ND2 | 2:Q:18:ALA:O | 2.40 | 0.54 |
| 1:K:666:GLN:NE2 | 1:K:865:ASP:OD1 | 2.41 | 0.54 |
| 2:Q:253:CYS:HB2 | 2:Q:296:GLY:HA2 | 1.90 | 0.54 |
| 2:S:252:HIS:HD2 | 2:S:256:GLY:HA3 | 1.73 | 0.54 |
| 2:T:252:HIS:HD2 | 2:T:256:GLY:HA3 | 1.73 | 0.54 |
| 1:D:417:ARG:NH2 | 1:D:782:GLN:OE1 | 2.38 | 0.54 |
| 1:G:379:GLY:H | 1:G:395:THR:HG21 | 1.73 | 0.54 |
| 1:K:350:ASN:OD1 | 1:K:416:LYS:NZ | 2.36 | 0.54 |
| 1:C:460:ILE:HD12 | 1:C:461:VAL:HG23 | 1.90 | 0.54 |
| 1:D:327:ARG:HB3 | 1:D:338:VAL:HG11 | 1.90 | 0.54 |
| 1:L:200:GLN:NE2 | 1:L:261:ASP:OD1 | 2.41 | 0.54 |
| 1:F:277:ARG:HH12 | 1:F:652:ILE:HD12 | 1.72 | 0.54 |

*Continued on next page...*

CONFIDENTIAL VALIDATION REPORT

*Continued from previous page...*

| Atom-1 | Atom-2 | Interatomic distance (Å) | Clash overlap (Å) |
|---|---|---|---|
| 2:R:253:CYS:HB2 | 2:R:296:GLY:HA2 | 1.90 | 0.54 |
| 2:S:253:CYS:HB2 | 2:S:296:GLY:HA2 | 1.90 | 0.54 |
| 1:F:354:GLU:OE2 | 1:D:486:ARG:NH2 | 2.40 | 0.53 |
| 1:I:600:LEU:HD12 | 2:Q:13:ILE:HD11 | 1.90 | 0.53 |
| 1:I:894:SER:HA | 3:P:112:GLY:HA3 | 1.90 | 0.53 |
| 1:J:460:ILE:HD12 | 1:J:461:VAL:HG23 | 1.89 | 0.53 |
| 1:K:862:PHE:HD2 | 1:K:873:LEU:HD11 | 1.72 | 0.53 |
| 1:L:549:ARG:NH1 | 1:L:646:PHE:O | 2.40 | 0.53 |
| 1:B:370:PRO:HB3 | 1:B:405:PRO:HB3 | 1.90 | 0.53 |
| 1:B:887:GLN:HG3 | 1:B:893:ILE:HG12 | 1.89 | 0.53 |
| 1:E:793:ARG:NH1 | 1:D:404:ILE:O | 2.41 | 0.53 |
| 1:G:352:GLY:O | 1:G:416:LYS:NZ | 2.41 | 0.53 |
| 5:M:98:ASN:ND2 | 5:M:136:TYR:O | 2.41 | 0.53 |
| 1:B:520:LEU:HB3 | 1:B:588:GLN:HE22 | 1.74 | 0.53 |
| 1:C:298:GLN:HB2 | 1:C:891:SER:HB3 | 1.89 | 0.53 |
| 1:E:109:LYS:NZ | 1:E:111:TYR:O | 2.40 | 0.53 |
| 1:G:143:PRO:HB2 | 1:G:171:GLY:HA2 | 1.90 | 0.53 |
| 2:Q:252:HIS:HD2 | 2:Q:256:GLY:HA3 | 1.73 | 0.53 |
| 1:E:102:LEU:HD11 | 1:E:540:LEU:HD11 | 1.89 | 0.53 |
| 1:F:885:ILE:HG12 | 1:F:895:VAL:HG12 | 1.89 | 0.53 |
| 1:H:100:GLY:HA2 | 1:H:558:THR:HG22 | 1.90 | 0.53 |
| 5:M:47:LEU:HB3 | 5:M:57:TYR:HB2 | 1.90 | 0.53 |
| 4:N:164:VAL:HG22 | 4:N:177:VAL:HG12 | 1.90 | 0.53 |
| 1:A:367:ILE:HG22 | 1:C:141:GLN:HB2 | 1.90 | 0.53 |
| 1:C:128:SER:OG | 1:C:201:GLN:NE2 | 2.41 | 0.53 |
| 1:G:62:ARG:HG3 | 1:G:563:TYR:HE1 | 1.74 | 0.53 |
| 1:J:304:ILE:HG23 | 1:J:895:VAL:HG11 | 1.91 | 0.53 |
| 1:E:346:ARG:NH2 | 1:E:477:ASP:O | 2.41 | 0.53 |
| 1:E:718:GLN:NE2 | 1:E:829:GLN:O | 2.39 | 0.53 |
| 1:F:167:ASN:HB3 | 1:F:170:VAL:HG23 | 1.90 | 0.53 |
| 1:G:299:THR:O | 1:G:301:GLN:NE2 | 2.42 | 0.53 |
| 1:K:491:LYS:NZ | 1:J:349:GLU:OE1 | 2.41 | 0.53 |
| 2:R:252:HIS:HD2 | 2:R:256:GLY:HA3 | 1.73 | 0.53 |
| 1:A:613:ASN:OD1 | 1:A:859:ASN:ND2 | 2.40 | 0.53 |
| 1:B:722:SER:OG | 1:B:829:GLN:NE2 | 2.41 | 0.53 |
| 1:H:815:GLN:HE21 | 1:I:502:TYR:H | 1.56 | 0.53 |
| 1:H:89:VAL:HG22 | 1:H:521:LEU:HB2 | 1.90 | 0.53 |
| 1:L:118:PRO:HD2 | 1:K:778:ASN:HB3 | 1.91 | 0.53 |
| 1:A:603:ILE:HG12 | 1:A:610:VAL:HG21 | 1.91 | 0.53 |
| 1:A:537:ASN:HD22 | 1:A:649:SER:HB3 | 1.74 | 0.53 |
| 1:B:637:ARG:HB3 | 5:M:46:ASN:HD21 | 1.74 | 0.53 |

*Continued on next page...*

CONFIDENTIAL VALIDATION REPORT

*Continued from previous page...*

| Atom-1 | Atom-2 | Interatomic distance (Å) | Clash overlap (Å) |
|---|---|---|---|
| 1:B:879:VAL:HG12 | 1:B:901:PRO:HD2 | 1.90 | 0.53 |
| 1:F:121:PRO:HG2 | 1:F:124:ALA:HB2 | 1.90 | 0.53 |
| 1:J:417:ARG:NH2 | 1:J:782:GLN:OE1 | 2.41 | 0.53 |
| 1:L:640:ALA:HB2 | 2:R:17:PRO:HD3 | 1.91 | 0.53 |
| 1:B:85:ASP:OD1 | 1:B:890:ARG:NH1 | 2.42 | 0.53 |
| 1:I:549:ARG:NH1 | 1:I:646:PHE:O | 2.41 | 0.53 |
| 2:T:253:CYS:HB2 | 2:T:296:GLY:HA2 | 1.90 | 0.53 |
| 1:A:350:ASN:OD1 | 1:A:416:LYS:NZ | 2.42 | 0.53 |
| 1:A:495:GLN:NE2 | 1:C:351:ASP:OD1 | 2.42 | 0.53 |
| 1:A:205:TYR:OH | 1:C:353:TYR:OH | 2.26 | 0.53 |
| 1:C:617:ARG:NH2 | 1:C:901:PRO:O | 2.41 | 0.53 |
| 1:F:718:GLN:NE2 | 1:F:829:GLN:O | 2.42 | 0.53 |
| 1:G:887:GLN:HG2 | 1:G:893:ILE:HG22 | 1.90 | 0.53 |
| 1:K:677:PRO:HG2 | 1:K:680:GLY:HA3 | 1.91 | 0.53 |
| 1:F:205:TYR:HH | 1:E:353:TYR:HH | 1.57 | 0.52 |
| 1:F:353:TYR:HB2 | 1:F:412:PRO:HB2 | 1.91 | 0.52 |
| 1:H:611:VAL:HG22 | 1:H:861:GLN:HG2 | 1.90 | 0.52 |
| 1:K:310:ASP:HB3 | 1:K:597:VAL:HG13 | 1.92 | 0.52 |
| 2:S:332:THR:HB | 2:S:335:LEU:HD13 | 1.92 | 0.52 |
| 2:T:330:GLU:HA | 2:T:352:PHE:HB2 | 1.92 | 0.52 |
| 1:H:97:ASP:OD1 | 1:G:712:LYS:NZ | 2.42 | 0.52 |
| 1:I:98:ILE:HG22 | 1:I:560:ILE:HG23 | 1.90 | 0.52 |
| 1:J:70:ASP:OD1 | 1:J:534:LYS:NZ | 2.40 | 0.52 |
| 1:K:414:ASN:HB3 | 1:J:359:ASN:HD22 | 1.75 | 0.52 |
| 1:K:582:ARG:NH1 | 1:K:889:THR:O | 2.41 | 0.52 |
| 1:A:199:GLY:HA3 | 1:A:248:TRP:HB2 | 1.91 | 0.52 |
| 1:H:174:ASN:HD21 | 1:G:783:GLN:HE22 | 1.55 | 0.52 |
| 1:H:511:GLN:NE2 | 1:H:528:TYR:OH | 2.41 | 0.52 |
| 1:K:144:GLN:NE2 | 1:K:170:VAL:O | 2.42 | 0.52 |
| 1:A:310:ASP:HB3 | 1:A:597:VAL:HG13 | 1.92 | 0.52 |
| 1:E:797:PRO:HD3 | 1:D:404:ILE:HD11 | 1.91 | 0.52 |
| 1:K:283:MET:HG2 | 1:K:509:VAL:HG11 | 1.91 | 0.52 |
| 1:C:277:ARG:HE | 1:C:541:GLN:HB2 | 1.75 | 0.52 |
| 1:E:331:PHE:HB3 | 1:E:336:GLN:HB3 | 1.92 | 0.52 |
| 1:H:102:LEU:HD11 | 1:H:540:LEU:HD11 | 1.91 | 0.52 |
| 1:K:718:GLN:NE2 | 1:K:829:GLN:O | 2.42 | 0.52 |
| 5:M:307:ASP:HA | 5:M:351:GLN:HE22 | 1.75 | 0.52 |
| 1:A:205:TYR:HH | 1:C:353:TYR:HH | 1.51 | 0.52 |
| 1:C:118:PRO:HD2 | 1:B:778:ASN:HB3 | 1.92 | 0.52 |
| 1:F:41:ARG:H | 1:E:589:ASN:HB2 | 1.73 | 0.52 |
| 1:G:24:VAL:HG12 | 1:G:28:GLN:HE22 | 1.74 | 0.52 |

*Continued on next page...*

CONFIDENTIAL VALIDATION REPORT

*Continued from previous page...*

| Atom-1 | Atom-2 | Interatomic distance (Å) | Clash overlap (Å) |
|---|---|---|---|
| 1:G:381:THR:HB | 1:G:390:ALA:HB3 | 1.91 | 0.52 |
| 1:H:144:GLN:HB2 | 1:H:189:LEU:HB3 | 1.92 | 0.52 |
| 2:R:330:GLU:HA | 2:R:352:PHE:HB2 | 1.92 | 0.52 |
| 1:A:102:LEU:HD11 | 1:A:540:LEU:HD11 | 1.92 | 0.52 |
| 1:C:168:PRO:HG3 | 1:C:191:ARG:HD3 | 1.92 | 0.52 |
| 1:J:520:LEU:HB3 | 1:J:588:GLN:HE22 | 1.75 | 0.52 |
| 1:L:186:ASN:O | 1:L:236:ARG:NH2 | 2.42 | 0.52 |
| 1:L:722:SER:OG | 1:L:829:GLN:NE2 | 2.43 | 0.52 |
| 1:A:301:GLN:NE2 | 1:L:60:GLN:O | 2.43 | 0.52 |
| 1:A:277:ARG:HH12 | 1:A:652:ILE:HD12 | 1.75 | 0.52 |
| 1:F:798:TYR:OH | 1:D:244:ASP:OD2 | 2.24 | 0.52 |
| 1:E:106:PRO:HD2 | 1:E:551:ASP:HB2 | 1.92 | 0.52 |
| 1:F:67:GLN:HE21 | 1:F:78:ARG:HE | 1.58 | 0.52 |
| 1:L:98:ILE:HG22 | 1:L:560:ILE:HG23 | 1.92 | 0.52 |
| 1:C:737:LYS:O | 1:C:743:ASN:ND2 | 2.38 | 0.51 |
| 1:I:778:ASN:HB3 | 1:G:118:PRO:HD2 | 1.92 | 0.51 |
| 1:I:798:TYR:OH | 1:G:244:ASP:OD2 | 2.24 | 0.51 |
| 4:N:177:VAL:O | 4:N:183:ASN:ND2 | 2.40 | 0.51 |
| 4:N:189:SER:OG | 4:N:218:ASN:ND2 | 2.43 | 0.51 |
| 2:R:332:THR:HB | 2:R:335:LEU:HD13 | 1.92 | 0.51 |
| 1:B:662:SER:O | 1:B:828:TRP:NE1 | 2.34 | 0.51 |
| 1:E:73:ASN:ND2 | 1:E:642:GLN:OE1 | 2.43 | 0.51 |
| 1:I:668:CYS:SG | 1:I:669:SER:N | 2.83 | 0.51 |
| 1:J:613:ASN:OD1 | 1:J:859:ASN:ND2 | 2.42 | 0.51 |
| 2:S:330:GLU:HA | 2:S:352:PHE:HB2 | 1.92 | 0.51 |
| 1:D:163:GLY:O | 1:D:191:ARG:NH1 | 2.43 | 0.51 |
| 1:F:346:ARG:HD3 | 1:F:481:PRO:HB3 | 1.91 | 0.51 |
| 1:H:843:THR:HG23 | 1:H:846:GLY:H | 1.74 | 0.51 |
| 1:L:41:ARG:H | 1:K:589:ASN:HB2 | 1.75 | 0.51 |
| 1:D:310:ASP:HB3 | 1:D:597:VAL:HG13 | 1.91 | 0.51 |
| 1:F:725:GLN:NE2 | 1:F:731:ASN:O | 2.43 | 0.51 |
| 1:G:160:GLN:HB3 | 2:S:91:THR:HG22 | 1.91 | 0.51 |
| 1:L:502:TYR:H | 1:K:815:GLN:HE21 | 1.59 | 0.51 |
| 5:M:180:ARG:HD2 | 5:M:245:PRO:HG3 | 1.92 | 0.51 |
| 2:R:151:PHE:HE2 | 2:R:167:ILE:HD11 | 1.76 | 0.51 |
| 1:A:118:PRO:HD2 | 1:C:778:ASN:HB3 | 1.93 | 0.51 |
| 1:D:725:GLN:NE2 | 1:D:731:ASN:OD1 | 2.43 | 0.51 |
| 1:G:347:ILE:HA | 1:G:472:SER:HB3 | 1.92 | 0.51 |
| 1:H:337:ALA:O | 1:H:493:ARG:NH1 | 2.44 | 0.51 |
| 1:L:379:GLY:HA2 | 1:K:228:ILE:HA | 1.91 | 0.51 |
| 2:S:151:PHE:HE2 | 2:S:167:ILE:HD11 | 1.76 | 0.51 |

*Continued on next page...*

CONFIDENTIAL VALIDATION REPORT

Continued from previous page...

| Atom-1 | Atom-2 | Interatomic distance (Å) | Clash overlap (Å) |
|---|---|---|---|
| 1:B:131:THR:HA | 1:B:136:THR:HA | 1.91 | 0.51 |
| 1:B:308:LEU:HD22 | 5:M:73:THR:HG23 | 1.92 | 0.51 |
| 1:E:367:ILE:HD13 | 1:D:139:VAL:HG23 | 1.92 | 0.51 |
| 1:E:118:PRO:HD2 | 1:D:778:ASN:HB3 | 1.92 | 0.51 |
| 1:K:97:ASP:OD1 | 1:J:712:LYS:NZ | 2.40 | 0.51 |
| 1:A:231:THR:HG21 | 1:A:236:ARG:HD3 | 1.93 | 0.51 |
| 1:D:156:ALA:O | 1:D:225:LYS:NZ | 2.43 | 0.51 |
| 1:E:337:ALA:O | 1:E:493:ARG:NH1 | 2.44 | 0.51 |
| 1:E:737:LYS:O | 1:E:743:ASN:ND2 | 2.44 | 0.51 |
| 1:B:537:ASN:HD22 | 1:B:649:SER:HB3 | 1.75 | 0.51 |
| 1:C:384:GLU:HG2 | 1:B:157:ILE:HD11 | 1.93 | 0.51 |
| 1:D:99:LYS:HD3 | 1:D:559:GLN:HE21 | 1.75 | 0.51 |
| 1:D:613:ASN:OD1 | 1:D:859:ASN:ND2 | 2.38 | 0.51 |
| 5:M:180:ARG:NH2 | 5:M:280:GLN:OE1 | 2.42 | 0.51 |
| 2:Q:330:GLU:HA | 2:Q:352:PHE:HB2 | 1.92 | 0.51 |
| 2:R:83:ILE:HG22 | 2:R:84:LEU:HB2 | 1.93 | 0.51 |
| 1:D:885:ILE:HG12 | 1:D:895:VAL:HG12 | 1.92 | 0.51 |
| 1:E:559:GLN:OE1 | 1:E:561:ASN:ND2 | 2.44 | 0.51 |
| 1:E:664:THR:HB | 1:E:865:ASP:HB2 | 1.93 | 0.51 |
| 1:H:163:GLY:O | 1:H:191:ARG:NH1 | 2.44 | 0.51 |
| 2:Q:332:THR:HB | 2:Q:335:LEU:HD13 | 1.92 | 0.51 |
| 2:Q:83:ILE:HG22 | 2:Q:84:LEU:HB2 | 1.93 | 0.51 |
| 1:C:879:VAL:HG12 | 1:C:901:PRO:HD2 | 1.93 | 0.51 |
| 1:F:141:GLN:HB2 | 1:D:367:ILE:HG22 | 1.92 | 0.51 |
| 1:F:273:TYR:OH | 1:E:351:ASP:OD1 | 2.29 | 0.51 |
| 1:H:117:ASN:HD22 | 1:G:778:ASN:HA | 1.75 | 0.51 |
| 4:N:169:SER:OG | 4:N:172:ASP:OD1 | 2.29 | 0.51 |
| 2:T:151:PHE:HE2 | 2:T:167:ILE:HD11 | 1.76 | 0.51 |
| 2:T:83:ILE:HG22 | 2:T:84:LEU:HB2 | 1.93 | 0.51 |
| 1:B:718:GLN:NE2 | 1:B:829:GLN:O | 2.44 | 0.50 |
| 1:D:705:THR:HG22 | 1:D:712:LYS:HD3 | 1.93 | 0.50 |
| 1:E:117:ASN:HD22 | 1:D:778:ASN:HA | 1.75 | 0.50 |
| 1:F:520:LEU:HB3 | 1:F:588:GLN:HE22 | 1.76 | 0.50 |
| 1:H:410:ASN:ND2 | 1:I:118:PRO:O | 2.37 | 0.50 |
| 1:K:277:ARG:HH12 | 1:K:652:ILE:HD12 | 1.76 | 0.50 |
| 1:F:327:ARG:HB3 | 1:F:338:VAL:HG21 | 1.94 | 0.50 |
| 1:H:630:LEU:O | 1:H:872:TYR:N | 2.44 | 0.50 |
| 1:I:347:ILE:HD13 | 1:I:472:SER:HB3 | 1.92 | 0.50 |
| 2:Q:151:PHE:HE2 | 2:Q:167:ILE:HD11 | 1.76 | 0.50 |
| 1:H:645:ASN:ND2 | 2:T:18:ALA:O | 2.44 | 0.50 |
| 1:C:339:ASP:OD1 | 1:C:493:ARG:NH2 | 2.45 | 0.50 |

Continued on next page...

CONFIDENTIAL VALIDATION REPORT

*Continued from previous page...*

| Atom-1 | Atom-2 | Interatomic distance (Å) | Clash overlap (Å) |
|---|---|---|---|
| 1:C:611:VAL:HG22 | 1:C:861:GLN:HG2 | 1.92 | 0.50 |
| 1:G:310:ASP:HB2 | 1:G:899:ARG:HH22 | 1.77 | 0.50 |
| 1:A:52:THR:HG21 | 1:C:834:SER:HB3 | 1.92 | 0.50 |
| 1:B:145:THR:HA | 1:B:188:GLY:HA2 | 1.92 | 0.50 |
| 1:D:704:ASP:O | 1:D:712:LYS:NZ | 2.44 | 0.50 |
| 1:J:168:PRO:HG3 | 1:J:191:ARG:HD3 | 1.93 | 0.50 |
| 1:A:885:ILE:HD11 | 1:A:893:ILE:HD11 | 1.94 | 0.50 |
| 1:D:284:MET:HB3 | 1:D:530:TRP:HE1 | 1.77 | 0.50 |
| 1:E:100:GLY:HA2 | 1:E:558:THR:HG22 | 1.94 | 0.50 |
| 1:F:725:GLN:HE21 | 1:F:731:ASN:H | 1.59 | 0.50 |
| 1:G:109:LYS:NZ | 1:G:111:TYR:O | 2.35 | 0.50 |
| 1:J:13:ARG:HB3 | 1:J:17:GLU:HG3 | 1.94 | 0.50 |
| 1:D:318:GLN:HG3 | 1:D:656:ASP:HA | 1.94 | 0.50 |
| 1:G:314:GLU:OE2 | 1:G:512:LYS:NZ | 2.43 | 0.50 |
| 1:J:352:GLY:O | 1:J:416:LYS:NZ | 2.43 | 0.50 |
| 1:L:236:ARG:NH1 | 1:L:238:SER:OG | 2.44 | 0.50 |
| 1:L:704:ASP:H | 1:L:712:LYS:HE3 | 1.75 | 0.50 |
| 1:B:201:GLN:NE2 | 1:B:261:ASP:O | 2.45 | 0.50 |
| 1:E:287:ASN:HA | 1:E:307:ASP:HB3 | 1.93 | 0.50 |
| 1:G:13:ARG:HB3 | 1:G:17:GLU:HG3 | 1.94 | 0.50 |
| 1:G:381:THR:HG1 | 1:G:392:THR:HG1 | 1.60 | 0.50 |
| 1:G:879:VAL:HG12 | 1:G:901:PRO:HD2 | 1.93 | 0.50 |
| 1:H:495:GLN:NE2 | 1:G:351:ASP:OD1 | 2.45 | 0.50 |
| 1:K:109:LYS:NZ | 1:K:111:TYR:O | 2.43 | 0.50 |
| 1:E:308:LEU:HD11 | 3:O:109:GLN:HE22 | 1.77 | 0.50 |
| 1:D:438:ASN:HD21 | 2:S:58:LYS:HB3 | 1.75 | 0.50 |
| 1:C:477:ASP:OD2 | 1:C:820:LYS:NZ | 2.37 | 0.50 |
| 1:D:62:ARG:HG3 | 1:D:563:TYR:HE1 | 1.77 | 0.50 |
| 1:I:143:PRO:O | 1:I:172:GLN:NE2 | 2.44 | 0.50 |
| 1:J:260:ASP:HB3 | 1:J:263:ARG:HB2 | 1.93 | 0.50 |
| 1:B:377:GLY:N | 1:B:397:ALA:O | 2.42 | 0.50 |
| 1:B:91:LEU:HD12 | 1:B:519:LEU:HB3 | 1.94 | 0.50 |
| 1:C:346:ARG:HD3 | 1:C:481:PRO:HB3 | 1.94 | 0.50 |
| 5:M:275:SER:HB3 | 5:M:278:LYS:HB2 | 1.94 | 0.50 |
| 1:A:64:VAL:O | 1:E:67:GLN:NE2 | 2.45 | 0.49 |
| 1:C:377:GLY:HA2 | 1:C:397:ALA:H | 1.77 | 0.49 |
| 1:C:619:TRP:HB2 | 1:C:849:VAL:HG11 | 1.93 | 0.49 |
| 1:D:843:THR:HG23 | 1:D:846:GLY:H | 1.77 | 0.49 |
| 1:I:673:ASP:HA | 1:I:857:SER:HB2 | 1.94 | 0.49 |
| 1:L:537:ASN:HD22 | 1:L:649:SER:HB3 | 1.77 | 0.49 |
| 1:B:905:GLY:HA2 | 5:M:75:THR:H | 1.77 | 0.49 |

*Continued on next page...*

CONFIDENTIAL VALIDATION REPORT

*Continued from previous page...*

| Atom-1 | Atom-2 | Interatomic distance (Å) | Clash overlap (Å) |
|---|---|---|---|
| 2:S:83:ILE:HG22 | 2:S:84:LEU:HB2 | 1.93 | 0.49 |
| 2:T:332:THR:HB | 2:T:335:LEU:HD13 | 1.92 | 0.49 |
| 1:F:353:TYR:HH | 1:D:205:TYR:HH | 1.58 | 0.49 |
| 1:E:107:SER:O | 1:E:272:ASN:ND2 | 2.45 | 0.49 |
| 1:E:614:ILE:HB | 1:E:858:LEU:HB3 | 1.94 | 0.49 |
| 1:I:783:GLN:NE2 | 1:I:792:GLU:O | 2.45 | 0.49 |
| 2:S:100:ASP:OD1 | 2:R:97:ARG:NH1 | 2.39 | 0.49 |
| 1:A:379:GLY:HA2 | 1:C:228:ILE:HG22 | 1.94 | 0.49 |
| 1:C:490:LEU:HB2 | 1:C:493:ARG:HH21 | 1.78 | 0.49 |
| 1:I:182:ASN:ND2 | 1:I:185:ASN:OD1 | 2.45 | 0.49 |
| 1:L:91:LEU:HD13 | 1:L:521:LEU:HD13 | 1.94 | 0.49 |
| 1:I:339:ASP:HA | 1:I:487:ASN:HD21 | 1.78 | 0.49 |
| 1:I:762:LEU:HD12 | 1:I:774:PRO:HB2 | 1.92 | 0.49 |
| 1:K:520:LEU:HB3 | 1:K:588:GLN:HE22 | 1.78 | 0.49 |
| 5:M:22:TYR:HD2 | 5:M:25:ILE:HG12 | 1.76 | 0.49 |
| 5:M:271:VAL:HG12 | 5:M:281:THR:HG22 | 1.94 | 0.49 |
| 1:B:53:THR:HB | 1:B:570:ASN:HA | 1.93 | 0.49 |
| 1:B:885:ILE:HG12 | 1:B:895:VAL:HG12 | 1.93 | 0.49 |
| 1:E:346:ARG:NH1 | 1:E:476:MET:O | 2.45 | 0.49 |
| 1:I:778:ASN:HA | 1:G:117:ASN:HD22 | 1.75 | 0.49 |
| 1:I:349:GLU:OE1 | 1:G:491:LYS:NZ | 2.46 | 0.49 |
| 1:H:297:SER:HA | 1:H:526:TYR:HA | 1.94 | 0.49 |
| 1:I:131:THR:HA | 1:I:136:THR:HA | 1.95 | 0.49 |
| 1:C:128:SER:H | 1:C:139:VAL:HG13 | 1.78 | 0.49 |
| 1:C:417:ARG:NH2 | 1:C:782:GLN:OE1 | 2.38 | 0.49 |
| 1:A:108:PHE:O | 1:C:807:SER:OG | 2.31 | 0.49 |
| 1:F:362:PHE:O | 1:D:408:GLU:N | 2.40 | 0.49 |
| 1:H:208:TYR:HB3 | 1:H:251:PRO:HD2 | 1.94 | 0.49 |
| 1:J:792:GLU:HG3 | 1:J:793:ARG:HD3 | 1.93 | 0.49 |
| 1:L:249:GLU:OE1 | 1:K:810:ASN:ND2 | 2.45 | 0.49 |
| 5:M:320:PRO:HG2 | 5:M:419:ARG:HD3 | 1.94 | 0.49 |
| 4:N:179:ILE:O | 4:N:183:ASN:ND2 | 2.45 | 0.49 |
| 1:A:246:ILE:HG22 | 1:A:248:TRP:H | 1.78 | 0.49 |
| 1:A:91:LEU:HD12 | 1:A:519:LEU:HB3 | 1.93 | 0.49 |
| 1:A:843:THR:HG23 | 1:A:846:GLY:H | 1.77 | 0.49 |
| 1:H:885:ILE:HD11 | 1:H:893:ILE:HD11 | 1.95 | 0.49 |
| 1:B:106:PRO:HD3 | 1:B:501:ARG:HH21 | 1.77 | 0.49 |
| 1:E:666:GLN:HA | 1:E:693:LYS:H | 1.76 | 0.49 |
| 1:E:522:LEU:HD12 | 1:E:887:GLN:HB2 | 1.94 | 0.49 |
| 1:J:611:VAL:HG22 | 1:J:861:GLN:HG2 | 1.95 | 0.49 |
| 1:K:696:ILE:HD13 | 1:K:819:ARG:HH11 | 1.76 | 0.49 |

*Continued on next page...*

CONFIDENTIAL VALIDATION REPORT

*Continued from previous page...*

| Atom-1 | Atom-2 | Interatomic distance (Å) | Clash overlap (Å) |
|---|---|---|---|
| 1:K:885:ILE:HD11 | 1:K:893:ILE:HD11 | 1.95 | 0.49 |
| 1:L:367:ILE:HG22 | 1:K:141:GLN:HB2 | 1.95 | 0.49 |
| 4:N:129:LEU:HG | 4:N:158:VAL:HG21 | 1.95 | 0.49 |
| 1:B:783:GLN:HE21 | 1:B:784:LYS:HG2 | 1.78 | 0.49 |
| 1:E:879:VAL:HG12 | 1:E:901:PRO:HD2 | 1.94 | 0.49 |
| 2:S:195:PHE:HB2 | 2:S:216:ALA:HA | 1.95 | 0.49 |
| 1:D:304:ILE:HG23 | 1:D:895:VAL:HG11 | 1.94 | 0.49 |
| 1:D:548:LEU:HD12 | 1:D:553:ALA:HB2 | 1.94 | 0.49 |
| 1:L:107:SER:O | 1:L:272:ASN:ND2 | 2.46 | 0.49 |
| 1:L:126:ILE:HD11 | 1:L:248:TRP:HH2 | 1.77 | 0.49 |
| 1:D:96:PHE:HB2 | 1:D:509:VAL:HG22 | 1.94 | 0.48 |
| 1:J:722:SER:OG | 1:J:829:GLN:NE2 | 2.46 | 0.48 |
| 1:L:10:ILE:HG22 | 1:K:842:LEU:HD12 | 1.94 | 0.48 |
| 5:M:192:TYR:OH | 5:M:245:PRO:O | 2.30 | 0.48 |
| 5:M:216:ILE:HA | 5:M:219:MET:HG3 | 1.94 | 0.48 |
| 2:S:171:ARG:NH2 | 2:R:135:THR:OG1 | 2.36 | 0.48 |
| 1:A:228:ILE:HA | 1:B:379:GLY:HA2 | 1.94 | 0.48 |
| 1:E:500:GLY:H | 1:D:751:GLN:HE21 | 1.61 | 0.48 |
| 1:C:19:LEU:HD13 | 1:C:23:LEU:HD12 | 1.95 | 0.48 |
| 1:D:168:PRO:HG3 | 1:D:191:ARG:HD3 | 1.95 | 0.48 |
| 1:F:131:THR:HA | 1:F:136:THR:HA | 1.95 | 0.48 |
| 1:F:778:ASN:HA | 1:D:117:ASN:HD22 | 1.77 | 0.48 |
| 1:J:668:CYS:HB3 | 1:J:690:PHE:HB2 | 1.96 | 0.48 |
| 1:L:353:TYR:HB2 | 1:L:412:PRO:HB2 | 1.93 | 0.48 |
| 5:M:36:ILE:HD11 | 5:M:438:ALA:HB3 | 1.94 | 0.48 |
| 2:R:195:PHE:HB2 | 2:R:216:ALA:HA | 1.95 | 0.48 |
| 1:A:691:GLU:OE1 | 1:A:694:ARG:NH2 | 2.44 | 0.48 |
| 1:A:39:LYS:NZ | 1:C:518:ASN:O | 2.46 | 0.48 |
| 1:D:350:ASN:ND2 | 1:D:465:THR:O | 2.38 | 0.48 |
| 1:D:297:SER:HA | 1:D:526:TYR:HA | 1.94 | 0.48 |
| 1:F:522:LEU:HB2 | 1:F:887:GLN:HG2 | 1.95 | 0.48 |
| 1:G:143:PRO:O | 1:G:172:GLN:NE2 | 2.46 | 0.48 |
| 5:M:75:THR:HG22 | 5:M:415:THR:HA | 1.94 | 0.48 |
| 5:M:54:SER:OG | 5:M:422:LEU:O | 2.27 | 0.48 |
| 1:C:283:MET:HG2 | 1:C:509:VAL:HG11 | 1.95 | 0.48 |
| 1:C:662:SER:O | 1:C:828:TRP:NE1 | 2.35 | 0.48 |
| 1:F:682:ASP:HA | 1:F:687:PRO:HB3 | 1.95 | 0.48 |
| 1:H:778:ASN:HB3 | 1:I:118:PRO:HD2 | 1.96 | 0.48 |
| 1:L:630:LEU:O | 1:L:872:TYR:N | 2.46 | 0.48 |
| 2:T:195:PHE:HB2 | 2:T:216:ALA:HA | 1.95 | 0.48 |
| 1:C:537:ASN:HD22 | 1:C:649:SER:HB3 | 1.78 | 0.48 |

*Continued on next page...*

CONFIDENTIAL VALIDATION REPORT

*Continued from previous page...*

| Atom-1 | Atom-2 | Interatomic distance (Å) | Clash overlap (Å) |
|---|---|---|---|
| 1:D:629:ARG:HD3 | 1:D:664:THR:HG23 | 1.94 | 0.48 |
| 1:F:287:ASN:HA | 1:F:307:ASP:HB3 | 1.94 | 0.48 |
| 1:G:287:ASN:ND2 | 1:G:307:ASP:OD1 | 2.46 | 0.48 |
| 1:G:612:VAL:HG22 | 1:G:860:MET:HB2 | 1.95 | 0.48 |
| 1:H:144:GLN:NE2 | 1:H:170:VAL:O | 2.46 | 0.48 |
| 1:I:141:GLN:HE21 | 1:G:367:ILE:HA | 1.79 | 0.48 |
| 1:L:281:ILE:HG21 | 1:L:655:LEU:HD13 | 1.94 | 0.48 |
| 1:E:314:GLU:HG3 | 1:E:655:LEU:HD22 | 1.94 | 0.48 |
| 1:G:471:TRP:CE2 | 1:G:750:ARG:HG2 | 2.49 | 0.48 |
| 1:I:362:PHE:O | 1:G:408:GLU:N | 2.36 | 0.48 |
| 2:Q:195:PHE:HB2 | 2:Q:216:ALA:HA | 1.95 | 0.48 |
| 1:A:181:VAL:HB | 1:B:401:TYR:HB3 | 1.95 | 0.48 |
| 1:A:228:ILE:HD13 | 1:B:399:ILE:HD13 | 1.95 | 0.48 |
| 1:B:417:ARG:NH2 | 1:B:782:GLN:OE1 | 2.47 | 0.48 |
| 1:K:623:ARG:HD3 | 1:K:880:PHE:HE1 | 1.78 | 0.48 |
| 1:H:200:GLN:O | 1:H:248:TRP:NE1 | 2.46 | 0.48 |
| 1:I:327:ARG:HD2 | 1:I:338:VAL:HG11 | 1.94 | 0.48 |
| 1:I:751:GLN:HG3 | 1:I:817:THR:HG22 | 1.96 | 0.48 |
| 1:G:182:ASN:ND2 | 1:G:185:ASN:OD1 | 2.46 | 0.48 |
| 1:H:230:THR:HG22 | 1:I:377:GLY:HA3 | 1.96 | 0.48 |
| 1:H:748:MET:SD | 1:H:820:LYS:NZ | 2.83 | 0.48 |
| 1:L:778:ASN:HA | 1:J:117:ASN:HD22 | 1.78 | 0.48 |
| 1:K:894:SER:HB3 | 3:O:35:ALA:HA | 1.95 | 0.48 |
| 1:A:150:GLY:O | 1:A:236:ARG:NH1 | 2.46 | 0.47 |
| 1:A:623:ARG:HD3 | 1:A:880:PHE:HE1 | 1.79 | 0.47 |
| 1:F:637:ARG:NH2 | 1:F:872:TYR:OH | 2.47 | 0.47 |
| 1:G:347:ILE:HG21 | 1:G:470:ARG:HE | 1.79 | 0.47 |
| 1:G:345:VAL:HG11 | 1:G:484:HIS:HE2 | 1.79 | 0.47 |
| 1:L:518:ASN:HB3 | 1:J:39:LYS:HE2 | 1.95 | 0.47 |
| 1:K:383:ASN:OD1 | 1:K:386:GLN:NE2 | 2.47 | 0.47 |
| 1:K:256:ALA:O | 1:K:788:ASN:ND2 | 2.47 | 0.47 |
| 1:L:619:TRP:HB2 | 1:L:849:VAL:HG11 | 1.96 | 0.47 |
| 1:B:640:ALA:H | 5:M:46:ASN:HD22 | 1.62 | 0.47 |
| 1:B:611:VAL:HG22 | 1:B:861:GLN:HG2 | 1.96 | 0.47 |
| 1:C:346:ARG:NH1 | 1:C:476:MET:O | 2.46 | 0.47 |
| 1:E:205:TYR:HH | 1:D:353:TYR:HH | 1.53 | 0.47 |
| 1:F:533:ARG:HH22 | 1:F:638:ILE:HG23 | 1.78 | 0.47 |
| 1:I:611:VAL:HG22 | 1:I:861:GLN:HG3 | 1.96 | 0.47 |
| 1:L:417:ARG:NH2 | 1:L:782:GLN:OE1 | 2.47 | 0.47 |
| 1:A:346:ARG:NH2 | 1:A:477:ASP:O | 2.42 | 0.47 |
| 1:F:387:GLN:HG2 | 1:E:154:THR:HB | 1.96 | 0.47 |

*Continued on next page...*

CONFIDENTIAL VALIDATION REPORT

*Continued from previous page...*

| Atom-1 | Atom-2 | Interatomic distance (Å) | Clash overlap (Å) |
|---|---|---|---|
| 1:H:73:ASN:ND2 | 1:H:642:GLN:OE1 | 2.47 | 0.47 |
| 1:I:287:ASN:HA | 1:I:307:ASP:HB3 | 1.96 | 0.47 |
| 1:K:169:GLN:NE2 | 1:J:775:ILE:O | 2.47 | 0.47 |
| 2:R:253:CYS:O | 2:R:258:ASN:ND2 | 2.48 | 0.47 |
| 1:D:331:PHE:HB3 | 1:D:336:GLN:HB3 | 1.97 | 0.47 |
| 1:H:520:LEU:HB2 | 1:I:39:LYS:HE2 | 1.96 | 0.47 |
| 1:I:470:ARG:NH2 | 1:I:746:GLU:OE2 | 2.47 | 0.47 |
| 1:K:118:PRO:HD2 | 1:J:778:ASN:HB3 | 1.96 | 0.47 |
| 5:M:370:ASN:O | 5:M:372:THR:OG1 | 2.28 | 0.47 |
| 1:D:98:ILE:HG13 | 1:D:560:ILE:HG12 | 1.97 | 0.47 |
| 1:F:590:PHE:H | 1:F:883:VAL:HG22 | 1.79 | 0.47 |
| 2:T:253:CYS:O | 2:T:258:ASN:ND2 | 2.48 | 0.47 |
| 1:B:630:LEU:O | 1:B:872:TYR:N | 2.41 | 0.47 |
| 1:C:489:GLY:O | 1:C:493:ARG:N | 2.40 | 0.47 |
| 1:D:183:THR:O | 1:D:236:ARG:NH1 | 2.47 | 0.47 |
| 1:D:1:MET:N | 1:D:1:MET:SD | 2.73 | 0.47 |
| 1:H:109:LYS:NZ | 1:H:111:TYR:O | 2.44 | 0.47 |
| 1:H:421:TYR:HA | 1:H:425:ALA:HB3 | 1.96 | 0.47 |
| 1:K:108:PHE:O | 1:J:807:SER:OG | 2.33 | 0.47 |
| 1:K:117:ASN:HA | 1:J:778:ASN:HB3 | 1.97 | 0.47 |
| 2:Q:97:ARG:HG2 | 2:Q:135:THR:HB | 1.97 | 0.47 |
| 1:J:471:TRP:CE2 | 1:J:750:ARG:HG2 | 2.49 | 0.47 |
| 2:Q:253:CYS:O | 2:Q:258:ASN:ND2 | 2.48 | 0.47 |
| 1:A:753:PRO:HD3 | 1:A:806:LEU:HD21 | 1.96 | 0.47 |
| 1:B:637:ARG:HH22 | 1:B:645:ASN:HB2 | 1.79 | 0.47 |
| 1:B:719:MET:HG3 | 1:B:725:GLN:HB2 | 1.96 | 0.47 |
| 1:C:668:CYS:HB3 | 1:C:690:PHE:HB2 | 1.97 | 0.47 |
| 1:E:166:PRO:HG2 | 1:E:191:ARG:HH11 | 1.79 | 0.47 |
| 1:E:885:ILE:HD11 | 1:E:893:ILE:HD11 | 1.96 | 0.47 |
| 1:H:346:ARG:NH2 | 1:H:477:ASP:O | 2.44 | 0.47 |
| 1:I:417:ARG:NH2 | 1:I:782:GLN:OE1 | 2.44 | 0.47 |
| 1:K:788:ASN:O | 1:K:790:LEU:N | 2.48 | 0.47 |
| 1:B:531:TYR:OH | 5:M:334:THR:O | 2.32 | 0.47 |
| 1:A:744:ASN:ND2 | 1:A:824:ASP:O | 2.47 | 0.47 |
| 1:A:712:LYS:HZ3 | 1:B:563:TYR:HE2 | 1.62 | 0.47 |
| 1:D:200:GLN:NE2 | 1:D:261:ASP:OD1 | 2.48 | 0.47 |
| 1:D:61:LEU:HG | 1:D:566:PHE:HE2 | 1.78 | 0.47 |
| 1:F:783:GLN:NE2 | 1:F:792:GLU:O | 2.48 | 0.47 |
| 1:J:589:ASN:HB3 | 1:J:884:VAL:HG12 | 1.97 | 0.47 |
| 1:L:409:MET:HB3 | 1:K:361:ALA:HA | 1.96 | 0.47 |
| 1:K:714:PHE:HE1 | 1:K:830:ILE:HD11 | 1.79 | 0.47 |

*Continued on next page...*

CONFIDENTIAL VALIDATION REPORT

*Continued from previous page...*

| Atom-1 | Atom-2 | Interatomic distance (Å) | Clash overlap (Å) |
|---|---|---|---|
| 1:K:885:ILE:HB | 1:K:895:VAL:HG12 | 1.97 | 0.47 |
| 5:M:17:ARG:HH12 | 5:M:449:THR:HG21 | 1.80 | 0.47 |
| 2:S:171:ARG:HH22 | 2:R:135:THR:HG1 | 1.57 | 0.47 |
| 2:S:253:CYS:O | 2:S:258:ASN:ND2 | 2.48 | 0.47 |
| 1:H:600:LEU:HD12 | 2:T:13:ILE:HD11 | 1.97 | 0.47 |
| 1:D:466:ASP:HB3 | 1:D:469:ALA:HB3 | 1.95 | 0.47 |
| 1:D:625:TRP:HD1 | 1:D:875:LEU:HD21 | 1.80 | 0.47 |
| 1:E:277:ARG:HH12 | 1:E:652:ILE:HD12 | 1.80 | 0.47 |
| 1:H:381:THR:HG23 | 1:H:392:THR:HB | 1.96 | 0.47 |
| 1:H:772:THR:HG23 | 1:H:774:PRO:HD3 | 1.96 | 0.47 |
| 1:J:693:LYS:HB2 | 1:J:709:ASN:HB3 | 1.97 | 0.47 |
| 1:K:150:GLY:O | 1:K:236:ARG:NH1 | 2.47 | 0.47 |
| 2:R:126:ILE:HB | 2:R:149:VAL:HG12 | 1.97 | 0.47 |
| 1:C:670:ILE:HG12 | 1:C:860:MET:HG2 | 1.95 | 0.47 |
| 1:D:381:THR:HB | 1:D:390:ALA:HB3 | 1.96 | 0.47 |
| 1:E:408:GLU:HB2 | 1:D:364:PRO:HG3 | 1.97 | 0.47 |
| 1:E:62:ARG:NH1 | 1:E:563:TYR:OH | 2.47 | 0.47 |
| 1:F:409:MET:HB3 | 1:E:361:ALA:HA | 1.95 | 0.47 |
| 1:H:310:ASP:HB3 | 1:H:597:VAL:HG13 | 1.97 | 0.47 |
| 1:H:725:GLN:NE2 | 1:H:731:ASN:OD1 | 2.48 | 0.47 |
| 1:I:532:PHE:HE2 | 1:I:560:ILE:HG21 | 1.79 | 0.47 |
| 1:K:379:GLY:HA2 | 1:J:228:ILE:HA | 1.97 | 0.47 |
| 1:K:614:ILE:HB | 1:K:858:LEU:HB3 | 1.97 | 0.47 |
| 2:S:97:ARG:HG2 | 2:S:135:THR:HB | 1.97 | 0.47 |
| 1:A:67:GLN:O | 1:A:76:LYS:NZ | 2.49 | 0.46 |
| 1:B:477:ASP:OD2 | 1:B:663:HIS:NE2 | 2.41 | 0.46 |
| 1:C:284:MET:HB3 | 1:C:530:TRP:HE1 | 1.80 | 0.46 |
| 1:D:693:LYS:HB2 | 1:D:709:ASN:HB3 | 1.96 | 0.46 |
| 1:A:491:LYS:NZ | 1:C:349:GLU:OE1 | 2.48 | 0.46 |
| 1:F:126:ILE:HD11 | 1:F:248:TRP:HH2 | 1.80 | 0.46 |
| 1:K:668:CYS:O | 1:K:690:PHE:N | 2.49 | 0.46 |
| 2:T:90:ARG:NH1 | 2:T:153:GLU:OE2 | 2.39 | 0.46 |
| 2:T:97:ARG:HG2 | 2:T:135:THR:HB | 1.96 | 0.46 |
| 1:B:236:ARG:NH1 | 1:B:238:SER:OG | 2.48 | 0.46 |
| 1:C:626:SER:HB2 | 1:C:876:LEU:HB2 | 1.96 | 0.46 |
| 1:D:182:ASN:O | 1:D:186:ASN:N | 2.47 | 0.46 |
| 1:I:85:ASP:OD1 | 1:I:890:ARG:NH1 | 2.48 | 0.46 |
| 1:K:499:ASN:ND2 | 1:J:466:ASP:OD1 | 2.47 | 0.46 |
| 1:B:301:GLN:NE2 | 5:M:332:ASN:OD1 | 2.49 | 0.46 |
| 1:E:165:ASP:OD1 | 2:R:93:LYS:NZ | 2.47 | 0.46 |
| 2:S:126:ILE:HB | 2:S:149:VAL:HG12 | 1.97 | 0.46 |

*Continued on next page...*

CONFIDENTIAL VALIDATION REPORT

*Continued from previous page...*

| Atom-1 | Atom-2 | Interatomic distance (Å) | Clash overlap (Å) |
|---|---|---|---|
| 1:C:118:PRO:O | 1:B:410:ASN:ND2 | 2.47 | 0.46 |
| 1:E:278:ASP:OD2 | 1:E:327:ARG:NH2 | 2.48 | 0.46 |
| 1:G:751:GLN:HG2 | 1:G:817:THR:HG22 | 1.97 | 0.46 |
| 1:I:342:ASP:OD2 | 1:I:486:ARG:NE | 2.48 | 0.46 |
| 1:I:662:SER:O | 1:I:828:TRP:NE1 | 2.41 | 0.46 |
| 1:J:256:ALA:O | 1:J:788:ASN:ND2 | 2.48 | 0.46 |
| 1:L:748:MET:HE3 | 1:L:820:LYS:HG3 | 1.98 | 0.46 |
| 2:R:97:ARG:HG2 | 2:R:135:THR:HB | 1.97 | 0.46 |
| 1:A:152:GLY:HA3 | 1:A:237:VAL:H | 1.80 | 0.46 |
| 1:A:628:THR:HG23 | 1:A:827:LEU:HA | 1.98 | 0.46 |
| 1:A:827:LEU:HD21 | 1:A:876:LEU:HD12 | 1.98 | 0.46 |
| 1:F:70:ASP:OD1 | 1:F:534:LYS:NZ | 2.41 | 0.46 |
| 1:H:244:ASP:OD2 | 1:G:798:TYR:OH | 2.33 | 0.46 |
| 1:H:773:MET:HB2 | 1:I:167:ASN:HD21 | 1.81 | 0.46 |
| 1:I:253:ALA:HA | 1:I:268:GLY:HA2 | 1.97 | 0.46 |
| 1:I:753:PRO:HD3 | 1:I:806:LEU:HD21 | 1.97 | 0.46 |
| 1:L:578:GLU:OE2 | 1:L:582:ARG:NE | 2.38 | 0.46 |
| 2:T:126:ILE:HB | 2:T:149:VAL:HG12 | 1.97 | 0.46 |
| 1:A:229:ASN:HB2 | 1:B:380:MET:HE1 | 1.97 | 0.46 |
| 1:A:294:SER:HA | 1:A:303:ASN:HD21 | 1.81 | 0.46 |
| 1:D:102:LEU:HD11 | 1:D:540:LEU:HD21 | 1.97 | 0.46 |
| 1:I:284:MET:SD | 1:I:530:TRP:NE1 | 2.89 | 0.46 |
| 2:R:90:ARG:NH1 | 2:R:153:GLU:OE2 | 2.39 | 0.46 |
| 1:H:708:SER:HB3 | 1:H:747:PRO:HB3 | 1.98 | 0.46 |
| 1:J:327:ARG:HB3 | 1:J:338:VAL:HG11 | 1.97 | 0.46 |
| 2:R:171:ARG:NH2 | 2:Q:135:THR:OG1 | 2.37 | 0.46 |
| 1:A:78:ARG:NH2 | 1:A:527:ASN:OD1 | 2.47 | 0.46 |
| 1:C:169:GLN:HA | 1:B:795:GLY:HA3 | 1.97 | 0.46 |
| 1:I:353:TYR:HB2 | 1:I:412:PRO:HB2 | 1.97 | 0.46 |
| 1:I:673:ASP:HB3 | 1:I:857:SER:H | 1.80 | 0.46 |
| 1:K:367:ILE:HG22 | 1:J:141:GLN:HB2 | 1.97 | 0.46 |
| 1:A:289:GLY:O | 1:K:694:ARG:NH2 | 2.49 | 0.46 |
| 5:M:150:VAL:HB | 5:M:377:ARG:HH21 | 1.81 | 0.46 |
| 1:A:751:GLN:HE22 | 1:B:498:GLY:HA3 | 1.81 | 0.46 |
| 1:B:522:LEU:HD12 | 1:B:887:GLN:HB2 | 1.97 | 0.46 |
| 1:E:491:LYS:NZ | 1:D:349:GLU:OE1 | 2.49 | 0.46 |
| 1:F:714:PHE:HE1 | 1:F:830:ILE:HD11 | 1.81 | 0.46 |
| 1:C:205:TYR:HB3 | 1:C:266:ALA:HB3 | 1.97 | 0.46 |
| 1:G:532:PHE:HE2 | 1:G:560:ILE:HG21 | 1.81 | 0.46 |
| 1:H:559:GLN:HE22 | 1:H:561:ASN:ND2 | 2.13 | 0.46 |
| 1:L:61:LEU:HB2 | 1:L:566:PHE:HE2 | 1.81 | 0.46 |

*Continued on next page...*

CONFIDENTIAL VALIDATION REPORT

*Continued from previous page...*

| Atom-1 | Atom-2 | Interatomic distance (Å) | Clash overlap (Å) |
|---|---|---|---|
| 1:B:306:LEU:H | 1:B:897:TYR:HE2 | 1.63 | 0.45 |
| 1:C:318:GLN:HG3 | 1:C:656:ASP:HA | 1.98 | 0.45 |
| 1:D:283:MET:HG2 | 1:D:509:VAL:HG11 | 1.99 | 0.45 |
| 1:F:466:ASP:OD1 | 1:D:499:ASN:ND2 | 2.49 | 0.45 |
| 1:F:511:GLN:OE1 | 1:F:528:TYR:OH | 2.33 | 0.45 |
| 1:F:719:MET:HG3 | 1:F:725:GLN:HB3 | 1.98 | 0.45 |
| 1:C:350:ASN:ND2 | 1:C:465:THR:O | 2.43 | 0.45 |
| 1:D:595:GLY:O | 1:D:623:ARG:NH2 | 2.49 | 0.45 |
| 1:E:783:GLN:NE2 | 1:E:792:GLU:O | 2.48 | 0.45 |
| 1:G:50:GLY:HA2 | 1:G:570:ASN:HD22 | 1.82 | 0.45 |
| 1:I:257:ASP:HB2 | 1:I:265:THR:HG22 | 1.98 | 0.45 |
| 1:H:277:ARG:HH12 | 1:H:652:ILE:HD12 | 1.81 | 0.45 |
| 1:L:496:LEU:HD23 | 1:K:707:GLN:HE21 | 1.81 | 0.45 |
| 1:A:511:GLN:NE2 | 1:A:528:TYR:OH | 2.41 | 0.45 |
| 1:A:69:GLU:HB3 | 1:A:76:LYS:HB3 | 1.97 | 0.45 |
| 1:A:666:GLN:NE2 | 1:A:865:ASP:OD1 | 2.49 | 0.45 |
| 1:B:297:SER:OG | 1:B:300:GLN:O | 2.33 | 0.45 |
| 1:B:619:TRP:HA | 1:B:901:PRO:HB3 | 1.98 | 0.45 |
| 5:M:138:VAL:HB | 5:M:308:ILE:HD11 | 1.99 | 0.45 |
| 5:M:57:TYR:OH | 5:M:419:ARG:NH1 | 2.50 | 0.45 |
| 1:J:159:GLN:HE22 | 2:R:109:THR:HG21 | 1.81 | 0.45 |
| 1:A:106:PRO:HD2 | 1:A:551:ASP:HB2 | 1.99 | 0.45 |
| 1:A:470:ARG:N | 1:B:495:GLN:OE1 | 2.49 | 0.45 |
| 1:E:148:ALA:O | 1:E:236:ARG:NH2 | 2.41 | 0.45 |
| 1:E:416:LYS:HE2 | 1:E:420:LEU:HD11 | 1.97 | 0.45 |
| 1:H:668:CYS:HG | 1:H:862:PHE:HE1 | 1.64 | 0.45 |
| 1:K:668:CYS:HB3 | 1:K:690:PHE:HB2 | 1.97 | 0.45 |
| 1:L:580:MET:O | 1:L:586:ASN:ND2 | 2.45 | 0.45 |
| 1:A:724:ASN:HD22 | 1:A:837:LEU:HD22 | 1.82 | 0.45 |
| 1:E:147:ALA:HA | 1:E:186:ASN:HD22 | 1.82 | 0.45 |
| 1:I:142:LEU:HD13 | 1:I:188:GLY:HA3 | 1.98 | 0.45 |
| 4:N:63:LEU:HD21 | 4:N:84:LEU:HD12 | 1.97 | 0.45 |
| 1:B:281:ILE:HG21 | 1:B:655:LEU:HD13 | 1.98 | 0.45 |
| 1:C:623:ARG:HD3 | 1:C:880:PHE:HE1 | 1.81 | 0.45 |
| 1:D:71:ASN:HB3 | 1:D:74:TYR:H | 1.82 | 0.45 |
| 1:K:664:THR:HB | 1:K:865:ASP:HB2 | 1.99 | 0.45 |
| 1:A:300:GLN:HG3 | 1:L:63:VAL:HG23 | 1.99 | 0.45 |
| 5:M:91:ILE:HG12 | 5:M:402:VAL:HG22 | 1.98 | 0.45 |
| 2:T:220:ARG:HE | 2:T:262:THR:HG22 | 1.82 | 0.45 |
| 1:A:314:GLU:HG3 | 1:A:655:LEU:HD22 | 1.98 | 0.45 |
| 1:B:537:ASN:HD21 | 1:B:548:LEU:H | 1.63 | 0.45 |

*Continued on next page...*

CONFIDENTIAL VALIDATION REPORT

*Continued from previous page...*

| Atom-1 | Atom-2 | Interatomic distance (Å) | Clash overlap (Å) |
|---|---|---|---|
| 1:E:297:SER:HA | 1:E:526:TYR:HA | 1.98 | 0.45 |
| 1:H:314:GLU:HG3 | 1:H:655:LEU:HD22 | 1.98 | 0.45 |
| 1:J:511:GLN:OE1 | 1:J:528:TYR:OH | 2.33 | 0.45 |
| 2:R:220:ARG:HE | 2:R:262:THR:HG22 | 1.82 | 0.45 |
| 1:G:159:GLN:HE21 | 1:G:189:LEU:HD22 | 1.81 | 0.45 |
| 1:G:27:ILE:O | 1:G:31:GLN:NE2 | 2.50 | 0.45 |
| 2:Q:126:ILE:HB | 2:Q:149:VAL:HG12 | 1.97 | 0.45 |
| 2:S:294:TYR:HE2 | 2:R:242:HIS:HB3 | 1.81 | 0.45 |
| 2:T:316:GLY:N | 2:T:343:GLY:O | 2.48 | 0.45 |
| 1:C:83:VAL:HG21 | 1:C:521:LEU:HD11 | 1.99 | 0.45 |
| 1:F:594:LEU:O | 1:F:623:ARG:NH2 | 2.50 | 0.45 |
| 2:S:316:GLY:N | 2:S:343:GLY:O | 2.48 | 0.45 |
| 1:A:725:GLN:NE2 | 1:A:731:ASN:OD1 | 2.49 | 0.44 |
| 1:D:662:SER:O | 1:D:828:TRP:NE1 | 2.41 | 0.44 |
| 1:F:537:ASN:HD22 | 1:F:649:SER:HB3 | 1.83 | 0.44 |
| 1:H:807:SER:HA | 1:I:501:ARG:HG2 | 1.97 | 0.44 |
| 1:K:197:SER:HG | 1:K:248:TRP:HE1 | 1.65 | 0.44 |
| 1:K:314:GLU:HG3 | 1:K:655:LEU:HD22 | 1.99 | 0.44 |
| 1:K:755:TYR:HA | 1:K:756:GLY:HA2 | 1.69 | 0.44 |
| 5:M:14:THR:HA | 5:M:29:GLN:HE21 | 1.81 | 0.44 |
| 2:Q:316:GLY:N | 2:Q:343:GLY:O | 2.48 | 0.44 |
| 1:A:118:PRO:O | 1:C:410:ASN:ND2 | 2.37 | 0.44 |
| 1:B:109:LYS:NZ | 1:B:111:TYR:O | 2.49 | 0.44 |
| 1:B:310:ASP:HB2 | 1:B:899:ARG:HH22 | 1.82 | 0.44 |
| 1:A:589:ASN:HB2 | 1:B:41:ARG:H | 1.83 | 0.44 |
| 1:B:533:ARG:HH11 | 1:B:538:MET:HG2 | 1.82 | 0.44 |
| 1:C:495:GLN:HE21 | 1:B:470:ARG:HG3 | 1.82 | 0.44 |
| 1:F:630:LEU:O | 1:F:872:TYR:N | 2.45 | 0.44 |
| 1:H:404:ILE:HD11 | 1:I:797:PRO:HD3 | 1.98 | 0.44 |
| 1:K:879:VAL:HG12 | 1:K:901:PRO:HD2 | 1.99 | 0.44 |
| 2:R:133:ARG:HG2 | 2:R:164:GLN:HG2 | 1.99 | 0.44 |
| 1:D:225:LYS:HA | 1:D:242:ALA:HA | 2.00 | 0.44 |
| 1:E:748:MET:SD | 1:E:820:LYS:NZ | 2.86 | 0.44 |
| 1:G:694:ARG:NH1 | 1:G:699:ASP:OD1 | 2.51 | 0.44 |
| 1:L:904:SER:HA | 1:D:676:VAL:HG22 | 1.98 | 0.44 |
| 3:P:89:PRO:HG3 | 3:P:212:VAL:HG22 | 2.00 | 0.44 |
| 2:T:133:ARG:HG2 | 2:T:164:GLN:HG2 | 1.99 | 0.44 |
| 1:A:277:ARG:HD3 | 1:A:541:GLN:HB2 | 2.00 | 0.44 |
| 1:C:156:ALA:O | 1:C:225:LYS:NZ | 2.41 | 0.44 |
| 1:E:21:GLU:OE2 | 1:E:25:GLN:NE2 | 2.50 | 0.44 |
| 1:E:78:ARG:HG3 | 1:E:529:GLU:HB3 | 1.99 | 0.44 |

*Continued on next page...*

CONFIDENTIAL VALIDATION REPORT

*Continued from previous page...*

| Atom-1 | Atom-2 | Interatomic distance (Å) | Clash overlap (Å) |
|---|---|---|---|
| 1:F:477:ASP:OD2 | 1:F:820:LYS:NZ | 2.40 | 0.44 |
| 1:F:85:ASP:OD1 | 1:F:890:ARG:NH1 | 2.50 | 0.44 |
| 1:H:773:MET:SD | 1:H:785:THR:OG1 | 2.72 | 0.44 |
| 1:L:287:ASN:HA | 1:L:307:ASP:HB3 | 1.98 | 0.44 |
| 1:B:227:TYR:HD1 | 1:B:237:VAL:HG11 | 1.83 | 0.44 |
| 1:C:379:GLY:HA2 | 1:B:228:ILE:HD12 | 1.98 | 0.44 |
| 1:D:640:ALA:HB2 | 2:S:17:PRO:HD3 | 2.00 | 0.44 |
| 1:E:504:ASP:O | 1:D:819:ARG:NH2 | 2.51 | 0.44 |
| 1:F:559:GLN:OE1 | 1:F:561:ASN:ND2 | 2.50 | 0.44 |
| 1:F:57:GLN:HE21 | 1:F:568:PRO:HA | 1.81 | 0.44 |
| 1:G:283:MET:HG2 | 1:G:509:VAL:HG11 | 1.99 | 0.44 |
| 1:H:779:CYS:HB2 | 1:I:169:GLN:HB3 | 2.00 | 0.44 |
| 1:I:783:GLN:HE21 | 1:I:791:LEU:HG | 1.81 | 0.44 |
| 1:K:291:ASN:HB2 | 1:K:531:TYR:CD2 | 2.53 | 0.44 |
| 1:K:363:PRO:HA | 1:K:364:PRO:HD3 | 1.83 | 0.44 |
| 2:Q:220:ARG:HE | 2:Q:262:THR:HG22 | 1.82 | 0.44 |
| 1:B:109:LYS:NZ | 1:B:113:GLY:O | 2.47 | 0.44 |
| 1:E:638:ILE:HD11 | 1:E:655:LEU:HD21 | 2.00 | 0.44 |
| 1:K:673:ASP:HA | 1:K:857:SER:HB3 | 1.99 | 0.44 |
| 1:L:377:GLY:O | 1:L:397:ALA:N | 2.51 | 0.44 |
| 2:R:300:ILE:HB | 2:R:331:THR:HG22 | 2.00 | 0.44 |
| 1:E:108:PHE:O | 1:D:807:SER:OG | 2.31 | 0.44 |
| 1:E:379:GLY:HA2 | 1:D:228:ILE:HG22 | 2.00 | 0.44 |
| 1:H:720:ALA:HB1 | 1:H:837:LEU:HD23 | 2.00 | 0.44 |
| 1:J:75:TYR:HB2 | 1:J:534:LYS:HE3 | 2.00 | 0.44 |
| 1:J:753:PRO:HD3 | 1:J:806:LEU:HD21 | 2.00 | 0.44 |
| 1:L:331:PHE:HB3 | 1:L:336:GLN:HE21 | 1.83 | 0.44 |
| 2:S:300:ILE:HB | 2:S:331:THR:HG22 | 2.00 | 0.44 |
| 1:A:50:GLY:HA2 | 1:A:570:ASN:HD22 | 1.83 | 0.44 |
| 1:A:630:LEU:O | 1:A:872:TYR:N | 2.51 | 0.44 |
| 1:C:616:ASP:HA | 1:C:857:SER:HB3 | 2.00 | 0.44 |
| 1:E:705:THR:HG23 | 1:E:712:LYS:HG2 | 2.00 | 0.44 |
| 1:F:98:ILE:HB | 1:F:560:ILE:HG12 | 1.99 | 0.44 |
| 1:I:834:SER:HB3 | 1:G:52:THR:HG21 | 2.00 | 0.44 |
| 1:I:347:ILE:HD11 | 1:I:822:LEU:HD12 | 2.00 | 0.44 |
| 1:H:751:GLN:HE22 | 1:I:498:GLY:HA3 | 1.82 | 0.44 |
| 5:M:57:TYR:HE1 | 5:M:419:ARG:HG3 | 1.83 | 0.44 |
| 2:S:220:ARG:HA | 2:S:258:ASN:HD21 | 1.83 | 0.44 |
| 1:B:630:LEU:HD22 | 1:B:653:PRO:HG2 | 2.00 | 0.44 |
| 1:C:363:PRO:HA | 1:C:364:PRO:HD3 | 1.81 | 0.44 |
| 1:G:98:ILE:HG12 | 1:G:560:ILE:HG12 | 2.00 | 0.44 |

*Continued on next page...*

CONFIDENTIAL VALIDATION REPORT

*Continued from previous page...*

| Atom-1 | Atom-2 | Interatomic distance (Å) | Clash overlap (Å) |
|---|---|---|---|
| 1:H:348:LEU:HD13 | 1:H:424:VAL:HG21 | 1.99 | 0.44 |
| 1:H:712:LYS:HG3 | 1:I:563:TYR:HE2 | 1.83 | 0.44 |
| 1:H:773:MET:HB3 | 1:I:169:GLN:HE21 | 1.83 | 0.44 |
| 1:I:310:ASP:HB3 | 1:I:597:VAL:HG13 | 1.99 | 0.44 |
| 1:L:249:GLU:HB2 | 1:K:810:ASN:HD21 | 1.83 | 0.44 |
| 1:L:773:MET:HB3 | 1:J:169:GLN:HE21 | 1.82 | 0.44 |
| 4:N:176:GLN:HG2 | 4:N:183:ASN:HB3 | 2.00 | 0.44 |
| 3:O:89:PRO:HG3 | 3:O:212:VAL:HG22 | 2.00 | 0.44 |
| 1:A:766:TYR:OH | 1:A:785:THR:O | 2.34 | 0.43 |
| 1:B:278:ASP:OD2 | 1:B:327:ARG:NH2 | 2.51 | 0.43 |
| 1:B:631:LYS:HG2 | 1:B:871:THR:HG22 | 1.99 | 0.43 |
| 1:C:107:SER:O | 1:C:272:ASN:ND2 | 2.50 | 0.43 |
| 1:E:363:PRO:HA | 1:E:364:PRO:HD3 | 1.87 | 0.43 |
| 1:F:118:PRO:HD2 | 1:E:778:ASN:HB3 | 2.00 | 0.43 |
| 1:G:140:ALA:HB2 | 1:G:192:ILE:HD13 | 2.00 | 0.43 |
| 1:G:339:ASP:N | 1:G:339:ASP:OD1 | 2.51 | 0.43 |
| 1:I:360:LEU:HD23 | 1:I:408:GLU:HB2 | 2.00 | 0.43 |
| 2:S:220:ARG:HE | 2:S:262:THR:HG22 | 1.82 | 0.43 |
| 1:B:107:SER:O | 1:B:272:ASN:ND2 | 2.51 | 0.43 |
| 1:D:511:GLN:OE1 | 1:D:528:TYR:OH | 2.35 | 0.43 |
| 1:D:848:ASN:HB3 | 1:D:851:TYR:HD2 | 1.83 | 0.43 |
| 1:F:231:THR:HG21 | 1:F:236:ARG:HD3 | 1.99 | 0.43 |
| 1:H:595:GLY:O | 1:H:623:ARG:NH2 | 2.52 | 0.43 |
| 1:L:297:SER:HA | 1:L:526:TYR:HA | 2.01 | 0.43 |
| 2:Q:90:ARG:NH1 | 2:Q:153:GLU:OE2 | 2.39 | 0.43 |
| 2:Q:300:ILE:HB | 2:Q:331:THR:HG22 | 2.00 | 0.43 |
| 2:T:349:VAL:H | 2:T:364:GLY:HA2 | 1.83 | 0.43 |
| 1:B:619:TRP:NE1 | 1:B:856:HIS:O | 2.45 | 0.43 |
| 1:F:470:ARG:NH2 | 1:F:747:PRO:O | 2.49 | 0.43 |
| 1:I:69:GLU:HB3 | 1:I:76:LYS:HB3 | 2.00 | 0.43 |
| 1:L:58:LYS:NZ | 1:K:684:MET:O | 2.49 | 0.43 |
| 1:A:850:MET:HA | 3:O:260:LEU:HD22 | 2.00 | 0.43 |
| 2:Q:349:VAL:H | 2:Q:364:GLY:HA2 | 1.84 | 0.43 |
| 1:C:408:GLU:N | 1:B:362:PHE:O | 2.43 | 0.43 |
| 1:C:797:PRO:HD3 | 1:B:404:ILE:HD11 | 2.00 | 0.43 |
| 1:B:827:LEU:HD13 | 1:B:876:LEU:HD21 | 1.99 | 0.43 |
| 1:E:691:GLU:OE2 | 1:E:694:ARG:NE | 2.43 | 0.43 |
| 1:H:363:PRO:HA | 1:H:364:PRO:HD3 | 1.84 | 0.43 |
| 1:H:778:ASN:HB3 | 1:I:117:ASN:HA | 1.99 | 0.43 |
| 1:I:98:ILE:HD11 | 1:I:507:ILE:HD11 | 2.00 | 0.43 |
| 1:J:19:LEU:HD13 | 1:J:23:LEU:HD12 | 2.00 | 0.43 |

*Continued on next page...*

CONFIDENTIAL VALIDATION REPORT

*Continued from previous page...*

| Atom-1 | Atom-2 | Interatomic distance (Å) | Clash overlap (Å) |
|---|---|---|---|
| 2:T:300:ILE:HB | 2:T:331:THR:HG22 | 2.00 | 0.43 |
| 1:A:2:GLU:HB3 | 1:A:5:ARG:HD2 | 1.99 | 0.43 |
| 1:B:98:ILE:HG22 | 1:B:560:ILE:HG23 | 1.99 | 0.43 |
| 1:D:225:LYS:HE2 | 1:D:227:TYR:HE1 | 1.84 | 0.43 |
| 1:D:718:GLN:NE2 | 1:D:829:GLN:O | 2.50 | 0.43 |
| 1:G:421:TYR:HA | 1:G:425:ALA:HB3 | 2.01 | 0.43 |
| 1:J:287:ASN:HA | 1:J:307:ASP:HB3 | 2.00 | 0.43 |
| 1:J:777:ASN:HD22 | 1:J:778:ASN:H | 1.67 | 0.43 |
| 1:K:666:GLN:HA | 1:K:693:LYS:H | 1.83 | 0.43 |
| 1:B:460:ILE:HD12 | 1:B:461:VAL:HG13 | 1.99 | 0.43 |
| 1:A:707:GLN:HE21 | 1:B:496:LEU:HD23 | 1.84 | 0.43 |
| 1:D:86:ASN:HD22 | 1:D:574:GLN:HE21 | 1.65 | 0.43 |
| 1:E:733:PRO:HG2 | 1:E:736:THR:HG22 | 2.01 | 0.43 |
| 1:F:879:VAL:HG12 | 1:F:901:PRO:HD2 | 2.00 | 0.43 |
| 1:G:363:PRO:HA | 1:G:364:PRO:HD3 | 1.86 | 0.43 |
| 1:I:279:ASN:ND2 | 1:I:318:GLN:OE1 | 2.48 | 0.43 |
| 1:J:128:SER:OG | 1:J:201:GLN:NE2 | 2.52 | 0.43 |
| 1:J:537:ASN:HD22 | 1:J:649:SER:HB3 | 1.83 | 0.43 |
| 1:K:724:ASN:HD22 | 1:K:837:LEU:HD22 | 1.83 | 0.43 |
| 5:M:112:LEU:HD12 | 5:M:127:PHE:HZ | 1.83 | 0.43 |
| 2:R:251:ASN:HD22 | 2:Q:187:TYR:HE1 | 1.66 | 0.43 |
| 2:S:133:ARG:HG2 | 2:S:164:GLN:HG2 | 1.99 | 0.43 |
| 1:F:52:THR:HG21 | 1:E:834:SER:HB3 | 1.99 | 0.43 |
| 1:G:383:ASN:OD1 | 1:G:388:THR:OG1 | 2.36 | 0.43 |
| 1:H:428:LEU:HD13 | 1:H:460:ILE:HD11 | 2.00 | 0.43 |
| 1:L:14:SER:OG | 1:L:17:GLU:OE1 | 2.30 | 0.43 |
| 1:L:766:TYR:OH | 1:L:785:THR:O | 2.28 | 0.43 |
| 5:M:271:VAL:HA | 5:M:281:THR:HA | 2.01 | 0.43 |
| 3:O:74:TYR:HB3 | 3:O:225:THR:HG22 | 2.01 | 0.43 |
| 2:Q:133:ARG:HG2 | 2:Q:164:GLN:HG2 | 1.99 | 0.43 |
| 1:A:284:MET:HB3 | 1:A:530:TRP:HE1 | 1.84 | 0.43 |
| 1:D:143:PRO:HB2 | 1:D:171:GLY:HA2 | 2.00 | 0.43 |
| 1:F:367:ILE:H | 1:F:367:ILE:HG13 | 1.67 | 0.43 |
| 1:F:305:VAL:HG22 | 1:F:516:ILE:HD11 | 2.00 | 0.43 |
| 1:J:98:ILE:HG13 | 1:J:560:ILE:HG12 | 2.01 | 0.43 |
| 5:M:341:VAL:HG12 | 5:M:343:GLY:H | 1.83 | 0.43 |
| 1:C:630:LEU:O | 1:C:872:TYR:N | 2.45 | 0.43 |
| 1:F:773:MET:SD | 1:F:773:MET:N | 2.92 | 0.43 |
| 1:I:331:PHE:HB3 | 1:I:336:GLN:HB3 | 1.99 | 0.43 |
| 1:K:842:LEU:HD21 | 1:K:847:GLN:HB3 | 2.00 | 0.43 |
| 1:L:537:ASN:HD21 | 1:L:548:LEU:H | 1.66 | 0.43 |

*Continued on next page...*

CONFIDENTIAL VALIDATION REPORT

*Continued from previous page...*

| Atom-1 | Atom-2 | Interatomic distance (Å) | Clash overlap (Å) |
|---|---|---|---|
| 2:R:294:TYR:HE2 | 2:Q:242:HIS:HB3 | 1.82 | 0.43 |
| 2:R:140:ASN:ND2 | 2:Q:95:ASP:OD2 | 2.43 | 0.43 |
| 2:R:220:ARG:HA | 2:R:258:ASN:HD21 | 1.83 | 0.43 |
| 2:R:349:VAL:H | 2:R:364:GLY:HA2 | 1.83 | 0.43 |
| 1:C:13:ARG:HB3 | 1:C:17:GLU:HG3 | 2.00 | 0.43 |
| 1:C:450:TYR:CZ | 1:C:454:ARG:HD2 | 2.53 | 0.43 |
| 1:H:62:ARG:HG3 | 1:H:563:TYR:HE1 | 1.84 | 0.43 |
| 1:L:62:ARG:HG3 | 1:L:563:TYR:HE1 | 1.83 | 0.43 |
| 1:L:67:GLN:HE21 | 1:L:78:ARG:CZ | 2.32 | 0.43 |
| 5:M:37:ILE:HD12 | 5:M:39:ASN:HB2 | 2.00 | 0.43 |
| 2:S:172:PHE:HE2 | 2:S:179:ILE:HG13 | 1.84 | 0.43 |
| 1:B:280:PHE:CZ | 1:B:335:ASN:HB2 | 2.54 | 0.42 |
| 1:E:843:THR:HG23 | 1:E:846:GLY:H | 1.82 | 0.42 |
| 1:K:36:ILE:HD12 | 1:J:520:LEU:HD22 | 2.00 | 0.42 |
| 1:A:139:VAL:HG22 | 1:B:367:ILE:HG21 | 2.01 | 0.42 |
| 1:C:678:TRP:CD1 | 1:C:679:PRO:HA | 2.54 | 0.42 |
| 1:F:117:ASN:HA | 1:E:778:ASN:HB3 | 1.99 | 0.42 |
| 1:F:46:ALA:O | 1:E:847:GLN:NE2 | 2.52 | 0.42 |
| 1:G:885:ILE:HG12 | 1:G:895:VAL:HG12 | 2.01 | 0.42 |
| 1:K:296:SER:HB3 | 1:K:303:ASN:HA | 2.02 | 0.42 |
| 1:K:304:ILE:O | 1:K:897:TYR:OH | 2.29 | 0.42 |
| 1:L:662:SER:O | 1:L:828:TRP:NE1 | 2.33 | 0.42 |
| 5:M:185:SER:HB2 | 5:M:278:LYS:HD3 | 2.01 | 0.42 |
| 2:Q:220:ARG:HA | 2:Q:258:ASN:HD21 | 1.83 | 0.42 |
| 1:A:899:ARG:NE | 1:A:902:PHE:O | 2.52 | 0.42 |
| 1:F:205:TYR:OH | 1:E:353:TYR:OH | 2.32 | 0.42 |
| 1:I:795:GLY:HA3 | 1:G:169:GLN:HA | 2.01 | 0.42 |
| 1:I:102:LEU:HD11 | 1:I:540:LEU:HD21 | 2.01 | 0.42 |
| 1:K:399:ILE:HG13 | 1:J:228:ILE:HD13 | 2.02 | 0.42 |
| 1:L:502:TYR:H | 1:K:815:GLN:NE2 | 2.17 | 0.42 |
| 1:L:717:VAL:HG23 | 1:L:836:PHE:HB3 | 2.01 | 0.42 |
| 5:M:54:SER:HA | 5:M:422:LEU:HB3 | 2.01 | 0.42 |
| 2:S:349:VAL:H | 2:S:364:GLY:HA2 | 1.84 | 0.42 |
| 1:A:348:LEU:HD13 | 1:A:424:VAL:HG21 | 2.01 | 0.42 |
| 1:D:88:LEU:HD12 | 1:D:520:LEU:HD11 | 2.02 | 0.42 |
| 1:D:69:GLU:HB3 | 1:D:76:LYS:HB3 | 2.01 | 0.42 |
| 1:F:143:PRO:HB2 | 1:F:171:GLY:HA2 | 2.01 | 0.42 |
| 1:G:205:TYR:HB3 | 1:G:266:ALA:HB3 | 2.02 | 0.42 |
| 1:I:327:ARG:HG2 | 1:I:738:HIS:CE1 | 2.55 | 0.42 |
| 1:K:787:THR:HB | 1:K:788:ASN:H | 1.61 | 0.42 |
| 2:R:172:PHE:HE2 | 2:R:179:ILE:HG13 | 1.84 | 0.42 |

*Continued on next page...*

CONFIDENTIAL VALIDATION REPORT

*Continued from previous page...*

| Atom-1 | Atom-2 | Interatomic distance (Å) | Clash overlap (Å) |
|---|---|---|---|
| 2:R:316:GLY:N | 2:R:343:GLY:O | 2.48 | 0.42 |
| 2:S:140:ASN:ND2 | 2:R:95:ASP:OD2 | 2.48 | 0.42 |
| 1:C:367:ILE:HG22 | 1:B:141:GLN:HB2 | 2.00 | 0.42 |
| 1:C:511:GLN:OE1 | 1:C:528:TYR:OH | 2.37 | 0.42 |
| 1:D:300:GLN:HE21 | 1:D:302:LEU:HD12 | 1.85 | 0.42 |
| 1:I:637:ARG:NH2 | 1:I:872:TYR:OH | 2.49 | 0.42 |
| 5:M:320:PRO:HD2 | 5:M:419:ARG:HB3 | 2.01 | 0.42 |
| 2:Q:172:PHE:HE2 | 2:Q:179:ILE:HG13 | 1.85 | 0.42 |
| 1:B:288:SER:O | 1:B:292:THR:OG1 | 2.33 | 0.42 |
| 1:B:304:ILE:HG23 | 1:B:895:VAL:HG11 | 2.01 | 0.42 |
| 1:C:342:ASP:OD2 | 1:C:486:ARG:NH1 | 2.52 | 0.42 |
| 1:E:665:PHE:HA | 1:E:864:VAL:HG12 | 2.00 | 0.42 |
| 1:I:263:ARG:HB2 | 1:G:175:TYR:CG | 2.55 | 0.42 |
| 1:H:39:LYS:HE2 | 1:G:520:LEU:HB2 | 2.02 | 0.42 |
| 1:I:270:ARG:NH1 | 1:I:426:MET:O | 2.41 | 0.42 |
| 1:J:423:ASN:HD21 | 1:J:486:ARG:HH21 | 1.67 | 0.42 |
| 5:M:324:LYS:HD2 | 5:M:409:PRO:HB2 | 2.01 | 0.42 |
| 1:C:30:THR:OG1 | 1:C:31:GLN:N | 2.50 | 0.42 |
| 1:C:629:ARG:HD3 | 1:C:664:THR:HG23 | 2.01 | 0.42 |
| 1:L:310:ASP:HB3 | 1:L:597:VAL:HG13 | 2.00 | 0.42 |
| 1:B:13:ARG:NH1 | 3:O:222:ILE:O | 2.45 | 0.42 |
| 1:A:75:TYR:HB2 | 1:A:534:LYS:HE2 | 2.02 | 0.42 |
| 1:C:499:ASN:ND2 | 1:B:466:ASP:OD1 | 2.52 | 0.42 |
| 1:E:201:GLN:NE2 | 1:E:261:ASP:OD2 | 2.41 | 0.42 |
| 1:H:287:ASN:ND2 | 1:H:307:ASP:OD1 | 2.53 | 0.42 |
| 1:H:365:HIS:O | 1:H:367:ILE:N | 2.53 | 0.42 |
| 1:I:718:GLN:HE22 | 1:I:829:GLN:H | 1.68 | 0.42 |
| 2:T:172:PHE:HE2 | 2:T:179:ILE:HG13 | 1.84 | 0.42 |
| 2:T:220:ARG:HA | 2:T:258:ASN:HD21 | 1.83 | 0.42 |
| 2:T:278:PHE:HE2 | 2:T:303:PHE:HB3 | 1.85 | 0.42 |
| 1:A:142:LEU:HD21 | 1:A:241:MET:HG2 | 2.02 | 0.42 |
| 1:C:17:GLU:HB2 | 4:N:9:SER:HB3 | 2.02 | 0.42 |
| 1:F:244:ASP:OD2 | 1:E:798:TYR:OH | 2.36 | 0.42 |
| 1:H:167:ASN:HD21 | 1:G:773:MET:HB2 | 1.85 | 0.42 |
| 1:H:278:ASP:OD2 | 1:H:327:ARG:NH2 | 2.52 | 0.42 |
| 1:L:714:PHE:HA | 1:L:717:VAL:HG12 | 2.02 | 0.42 |
| 1:C:467:ILE:HG13 | 1:C:467:ILE:H | 1.65 | 0.42 |
| 1:I:367:ILE:HG13 | 1:I:367:ILE:H | 1.42 | 0.42 |
| 1:J:421:TYR:HA | 1:J:425:ALA:HB3 | 2.02 | 0.42 |
| 5:M:115:ASP:HA | 5:M:160:LYS:HG2 | 2.02 | 0.42 |
| 3:P:74:TYR:HB3 | 3:P:225:THR:HG22 | 2.01 | 0.42 |

*Continued on next page...*

CONFIDENTIAL VALIDATION REPORT

*Continued from previous page...*

| Atom-1 | Atom-2 | Interatomic distance (Å) | Clash overlap (Å) |
|---|---|---|---|
| 1:C:102:LEU:HD11 | 1:C:540:LEU:HD21 | 2.02 | 0.41 |
| 1:F:537:ASN:HD21 | 1:F:548:LEU:H | 1.68 | 0.41 |
| 1:I:773:MET:HB2 | 1:G:167:ASN:HD21 | 1.85 | 0.41 |
| 1:H:482:PHE:HZ | 1:H:825:ARG:HD2 | 1.85 | 0.41 |
| 1:H:228:ILE:HG21 | 1:I:399:ILE:HD13 | 2.02 | 0.41 |
| 1:K:52:THR:HG21 | 1:J:834:SER:HB3 | 2.02 | 0.41 |
| 1:K:736:THR:HB | 1:K:743:ASN:HD21 | 1.85 | 0.41 |
| 1:L:169:GLN:HB3 | 1:K:779:CYS:HB2 | 2.01 | 0.41 |
| 5:M:170:LYS:NZ | 5:M:172:ASP:OD1 | 2.50 | 0.41 |
| 1:A:417:ARG:NH2 | 1:A:782:GLN:OE1 | 2.53 | 0.41 |
| 1:C:191:ARG:NH2 | 1:C:244:ASP:OD2 | 2.52 | 0.41 |
| 1:D:139:VAL:HB | 1:D:262:ARG:HH12 | 1.84 | 0.41 |
| 1:E:744:ASN:ND2 | 1:E:824:ASP:O | 2.52 | 0.41 |
| 1:H:724:ASN:HD22 | 1:H:837:LEU:HD22 | 1.84 | 0.41 |
| 1:J:178:PRO:HG3 | 1:J:365:HIS:CE1 | 2.56 | 0.41 |
| 1:J:377:GLY:HA2 | 1:J:397:ALA:H | 1.85 | 0.41 |
| 1:J:96:PHE:HB2 | 1:J:509:VAL:HG22 | 2.02 | 0.41 |
| 1:L:483:ASN:ND2 | 1:L:648:TYR:OH | 2.40 | 0.41 |
| 5:M:143:ASP:OD1 | 5:M:386:GLN:NE2 | 2.53 | 0.41 |
| 2:S:90:ARG:NH1 | 2:S:153:GLU:OE2 | 2.39 | 0.41 |
| 2:S:200:ILE:HG12 | 2:S:221:CYS:HB3 | 2.03 | 0.41 |
| 2:S:214:ASN:N | 2:S:248:ASN:OD1 | 2.53 | 0.41 |
| 1:B:365:HIS:CD2 | 1:B:368:SER:HB3 | 2.55 | 0.41 |
| 1:B:668:CYS:HB3 | 1:B:690:PHE:HB2 | 2.01 | 0.41 |
| 1:B:751:GLN:HG3 | 1:B:817:THR:HB | 2.02 | 0.41 |
| 1:C:725:GLN:NE2 | 1:C:731:ASN:OD1 | 2.53 | 0.41 |
| 1:D:71:ASN:ND2 | 1:D:73:ASN:OD1 | 2.53 | 0.41 |
| 1:G:107:SER:O | 1:G:272:ASN:ND2 | 2.53 | 0.41 |
| 1:G:71:ASN:HD22 | 1:G:71:ASN:HA | 1.65 | 0.41 |
| 1:I:161:VAL:HG12 | 1:I:166:PRO:HD3 | 2.03 | 0.41 |
| 1:J:630:LEU:O | 1:J:872:TYR:N | 2.43 | 0.41 |
| 1:A:735:CYS:SG | 1:A:737:LYS:NZ | 2.88 | 0.41 |
| 1:B:714:PHE:HA | 1:B:717:VAL:HG12 | 2.01 | 0.41 |
| 1:D:678:TRP:CD1 | 1:D:679:PRO:HA | 2.56 | 0.41 |
| 1:F:578:GLU:OE2 | 1:F:582:ARG:NE | 2.45 | 0.41 |
| 1:F:722:SER:OG | 1:F:829:GLN:NE2 | 2.53 | 0.41 |
| 1:I:428:LEU:HD13 | 1:I:460:ILE:HD11 | 2.01 | 0.41 |
| 1:J:450:TYR:CZ | 1:J:454:ARG:HD2 | 2.55 | 0.41 |
| 1:J:102:LEU:HD11 | 1:J:540:LEU:HD21 | 2.02 | 0.41 |
| 1:L:327:ARG:HH12 | 1:L:336:GLN:HB2 | 1.85 | 0.41 |
| 2:Q:214:ASN:N | 2:Q:248:ASN:OD1 | 2.54 | 0.41 |

*Continued on next page...*

CONFIDENTIAL VALIDATION REPORT

*Continued from previous page...*

| Atom-1 | Atom-2 | Interatomic distance (Å) | Clash overlap (Å) |
|---|---|---|---|
| 2:R:214:ASN:N | 2:R:248:ASN:OD1 | 2.53 | 0.41 |
| 2:S:251:ASN:HD22 | 2:R:187:TYR:HE1 | 1.69 | 0.41 |
| 1:A:500:GLY:H | 1:C:751:GLN:HE21 | 1.68 | 0.41 |
| 1:D:638:ILE:HD11 | 1:D:655:LEU:HD11 | 2.02 | 0.41 |
| 1:F:89:VAL:HB | 1:F:521:LEU:HB3 | 2.01 | 0.41 |
| 1:G:630:LEU:O | 1:G:872:TYR:N | 2.49 | 0.41 |
| 1:H:796:HIS:HB2 | 1:I:127:ASN:HD21 | 1.85 | 0.41 |
| 1:J:167:ASN:HB3 | 1:J:170:VAL:HG22 | 2.03 | 0.41 |
| 5:M:314:GLN:NE2 | 5:M:316:TYR:OH | 2.42 | 0.41 |
| 2:Q:278:PHE:HE2 | 2:Q:303:PHE:HB3 | 1.85 | 0.41 |
| 1:D:295:PHE:HD2 | 1:D:304:ILE:HD12 | 1.86 | 0.41 |
| 1:D:520:LEU:HB3 | 1:D:588:GLN:HE22 | 1.84 | 0.41 |
| 1:E:668:CYS:HB3 | 1:E:690:PHE:HB2 | 2.01 | 0.41 |
| 1:F:380:MET:HA | 1:F:392:THR:HG22 | 2.03 | 0.41 |
| 1:L:611:VAL:HG11 | 2:Q:11:ASN:HD22 | 1.84 | 0.41 |
| 5:M:337:SER:HB2 | 5:M:427:ARG:HG3 | 2.02 | 0.41 |
| 1:B:126:ILE:HD11 | 1:B:248:TRP:HH2 | 1.86 | 0.41 |
| 1:E:563:TYR:HE2 | 1:D:712:LYS:HE2 | 1.85 | 0.41 |
| 1:F:670:ILE:HG12 | 1:F:860:MET:HG2 | 2.03 | 0.41 |
| 1:G:843:THR:HG23 | 1:G:846:GLY:H | 1.85 | 0.41 |
| 1:G:296:SER:HA | 1:G:893:ILE:HD11 | 2.02 | 0.41 |
| 1:J:662:SER:O | 1:J:828:TRP:NE1 | 2.48 | 0.41 |
| 1:L:167:ASN:HD22 | 1:L:168:PRO:HD2 | 1.86 | 0.41 |
| 1:K:13:ARG:NH1 | 3:P:222:ILE:O | 2.45 | 0.41 |
| 2:R:200:ILE:HG12 | 2:R:221:CYS:HB3 | 2.03 | 0.41 |
| 2:S:278:PHE:HE2 | 2:S:303:PHE:HB3 | 1.85 | 0.41 |
| 1:F:408:GLU:HB2 | 1:E:364:PRO:HG3 | 2.01 | 0.41 |
| 1:G:126:ILE:HD11 | 1:G:248:TRP:CH2 | 2.55 | 0.41 |
| 1:G:656:ASP:N | 1:G:656:ASP:OD1 | 2.50 | 0.41 |
| 1:I:377:GLY:N | 1:I:397:ALA:O | 2.52 | 0.41 |
| 2:S:95:ASP:OD2 | 2:Q:140:ASN:ND2 | 2.47 | 0.41 |
| 2:T:156:TYR:HE1 | 2:T:184:ARG:HB3 | 1.86 | 0.41 |
| 1:B:668:CYS:O | 1:B:690:PHE:N | 2.53 | 0.41 |
| 1:C:216:GLY:HA3 | 1:B:774:PRO:HB2 | 2.02 | 0.41 |
| 1:D:272:ASN:HA | 1:D:543:THR:HB | 2.03 | 0.41 |
| 1:F:428:LEU:HD11 | 1:F:460:ILE:HD11 | 2.02 | 0.41 |
| 1:F:774:PRO:HB2 | 1:D:216:GLY:HA3 | 2.02 | 0.41 |
| 1:I:294:SER:HA | 1:I:303:ASN:HD21 | 1.85 | 0.41 |
| 1:J:629:ARG:HD3 | 1:J:664:THR:HG23 | 2.02 | 0.41 |
| 1:J:751:GLN:HG2 | 1:J:817:THR:HG22 | 2.03 | 0.41 |
| 1:K:69:GLU:HG2 | 1:K:76:LYS:HB3 | 2.02 | 0.41 |

*Continued on next page...*

CONFIDENTIAL VALIDATION REPORT

*Continued from previous page...*

| Atom-1 | Atom-2 | Interatomic distance (Å) | Clash overlap (Å) |
|---|---|---|---|
| 1:A:616:ASP:OD1 | 1:A:616:ASP:N | 2.53 | 0.41 |
| 1:A:733:PRO:HG2 | 1:A:736:THR:HG22 | 2.03 | 0.41 |
| 1:B:290:SER:HA | 5:M:57:TYR:HE2 | 1.86 | 0.41 |
| 1:B:626:SER:HB2 | 1:B:876:LEU:HG | 2.03 | 0.41 |
| 1:F:292:THR:HG22 | 1:F:307:ASP:HB2 | 2.02 | 0.41 |
| 1:H:367:ILE:HD13 | 1:G:139:VAL:HG13 | 2.03 | 0.41 |
| 1:H:148:ALA:HB2 | 1:H:155:GLU:HB2 | 2.02 | 0.41 |
| 1:I:141:GLN:NE2 | 1:G:367:ILE:HA | 2.36 | 0.41 |
| 1:J:297:SER:HA | 1:J:526:TYR:HA | 2.03 | 0.41 |
| 1:K:347:ILE:HG21 | 1:K:470:ARG:HE | 1.85 | 0.41 |
| 1:L:151:THR:HA | 1:L:236:ARG:HA | 2.02 | 0.41 |
| 5:M:244:ILE:HD11 | 5:M:285:SER:HB3 | 2.02 | 0.41 |
| 2:T:200:ILE:HD13 | 2:T:224:LEU:HD22 | 2.03 | 0.41 |
| 2:T:200:ILE:HG12 | 2:T:221:CYS:HB3 | 2.03 | 0.41 |
| 1:B:291:ASN:HB3 | 1:B:531:TYR:HD2 | 1.86 | 0.41 |
| 1:A:403:ASN:HD21 | 1:B:794:CYS:HA | 1.86 | 0.41 |
| 1:D:269:ASN:ND2 | 1:D:452:ASN:HD21 | 2.19 | 0.41 |
| 1:D:753:PRO:HD3 | 1:D:806:LEU:HD21 | 2.03 | 0.41 |
| 1:K:460:ILE:H | 1:K:460:ILE:HG13 | 1.75 | 0.41 |
| 3:P:79:GLN:NE2 | 3:P:224:ARG:H | 2.19 | 0.41 |
| 1:L:134:ASN:HD22 | 2:R:118:GLU:HG2 | 1.85 | 0.41 |
| 1:D:208:TYR:HB3 | 1:D:251:PRO:HD2 | 2.02 | 0.40 |
| 1:D:363:PRO:HA | 1:D:364:PRO:HD3 | 1.81 | 0.40 |
| 1:E:291:ASN:N | 1:E:291:ASN:OD1 | 2.54 | 0.40 |
| 1:E:428:LEU:HD21 | 1:E:460:ILE:HD13 | 2.03 | 0.40 |
| 1:F:279:ASN:ND2 | 1:F:318:GLN:OE1 | 2.47 | 0.40 |
| 1:I:773:MET:HB3 | 1:G:169:GLN:HE21 | 1.86 | 0.40 |
| 1:G:254:HIS:ND1 | 1:G:269:ASN:OD1 | 2.49 | 0.40 |
| 1:H:169:GLN:NE2 | 1:G:775:ILE:O | 2.54 | 0.40 |
| 1:H:293:GLY:HA3 | 1:H:530:TRP:CE3 | 2.56 | 0.40 |
| 1:H:783:GLN:HG2 | 1:H:788:ASN:O | 2.21 | 0.40 |
| 1:I:589:ASN:HB3 | 1:I:884:VAL:HG12 | 2.03 | 0.40 |
| 1:L:403:ASN:HD21 | 1:K:172:GLN:HB2 | 1.85 | 0.40 |
| 5:M:286:TRP:CD1 | 5:M:297:LEU:HD21 | 2.56 | 0.40 |
| 2:R:278:PHE:HE2 | 2:R:303:PHE:HB3 | 1.85 | 0.40 |
| 2:S:141:PHE:HD2 | 2:S:148:ALA:HB1 | 1.86 | 0.40 |
| 1:D:126:ILE:HD11 | 1:D:248:TRP:CH2 | 2.57 | 0.40 |
| 1:E:284:MET:HB3 | 1:E:530:TRP:HE1 | 1.85 | 0.40 |
| 1:G:668:CYS:HB3 | 1:G:690:PHE:HB2 | 2.02 | 0.40 |
| 5:M:66:VAL:HG22 | 5:M:70:ASP:HB3 | 2.03 | 0.40 |
| 4:N:212:SER:HA | 4:N:215:LEU:HD23 | 2.02 | 0.40 |

*Continued on next page...*

CONFIDENTIAL VALIDATION REPORT

*Continued from previous page...*

| Atom-1 | Atom-2 | Interatomic distance (Å) | Clash overlap (Å) |
|---|---|---|---|
| 2:S:156:TYR:HE1 | 2:S:184:ARG:HB3 | 1.86 | 0.40 |
| 2:T:214:ASN:N | 2:T:248:ASN:OD1 | 2.54 | 0.40 |
| 1:C:699:ASP:OD1 | 1:C:702:GLY:N | 2.51 | 0.40 |
| 1:H:784:LYS:HD2 | 1:I:170:VAL:HA | 2.02 | 0.40 |
| 1:L:773:MET:HG3 | 1:J:167:ASN:HD21 | 1.85 | 0.40 |
| 1:J:906:SER:OG | 1:J:907:ALA:N | 2.53 | 0.40 |
| 1:L:117:ASN:HA | 1:K:778:ASN:HB3 | 2.04 | 0.40 |
| 5:M:215:ARG:HG3 | 5:M:218:ASN:HD22 | 1.86 | 0.40 |
| 1:D:611:VAL:HG11 | 2:R:11:ASN:HD22 | 1.87 | 0.40 |
| 2:R:200:ILE:HD13 | 2:R:224:LEU:HD22 | 2.03 | 0.40 |
| 1:C:345:VAL:HG11 | 1:C:484:HIS:NE2 | 2.37 | 0.40 |
| 1:C:610:VAL:HG11 | 1:C:873:LEU:HD22 | 2.04 | 0.40 |
| 1:D:532:PHE:HE2 | 1:D:560:ILE:HG21 | 1.85 | 0.40 |
| 1:F:595:GLY:O | 1:F:623:ARG:NH2 | 2.47 | 0.40 |
| 1:H:432:TYR:HE2 | 1:H:475:VAL:HG11 | 1.86 | 0.40 |
| 1:L:300:GLN:O | 1:L:302:LEU:N | 2.54 | 0.40 |
| 5:M:111:LEU:HD13 | 5:M:126:TRP:CD2 | 2.56 | 0.40 |
| 1:B:167:ASN:HD22 | 1:B:168:PRO:HD2 | 1.87 | 0.40 |
| 1:B:98:ILE:HD11 | 1:B:507:ILE:HD11 | 2.03 | 0.40 |
| 1:A:170:VAL:HG22 | 1:C:784:LYS:HD3 | 2.03 | 0.40 |
| 1:F:796:HIS:HB2 | 1:D:127:ASN:HD21 | 1.87 | 0.40 |
| 1:E:379:GLY:HA2 | 1:D:228:ILE:HA | 2.02 | 0.40 |
| 1:H:277:ARG:HD3 | 1:H:541:GLN:HB2 | 2.03 | 0.40 |
| 1:I:159:GLN:HB2 | 1:I:225:LYS:HZ2 | 1.87 | 0.40 |
| 1:I:714:PHE:HA | 1:I:717:VAL:HG12 | 2.04 | 0.40 |
| 1:J:205:TYR:HB3 | 1:J:266:ALA:HB3 | 2.03 | 0.40 |
| 1:L:98:ILE:HD11 | 1:L:507:ILE:HD11 | 2.03 | 0.40 |
| 1:L:724:ASN:HD21 | 1:L:838:ASN:H | 1.69 | 0.40 |
| 2:Q:156:TYR:HE1 | 2:Q:184:ARG:HB3 | 1.86 | 0.40 |
| 2:Q:229:MET:HA | 2:Q:266:LEU:HD22 | 2.04 | 0.40 |
| 2:R:141:PHE:HD2 | 2:R:148:ALA:HB1 | 1.86 | 0.40 |

There are no symmetry-related clashes.

#### 5.3 Torsion angles (i)

##### 5.3.1 Protein backbone (i)

In the following table, the Percentiles column shows the percent Ramachandran outliers of the chain as a percentile score with respect to all PDB entries followed by that with respect to all EM entries.

CONFIDENTIAL VALIDATION REPORT

The Analysed column shows the number of residues for which the backbone conformation was analysed, and the total number of residues.

| Mol | Chain | Analysed | Favoured | Allowed | Outliers | Percentiles | |
|---|---|---|---|---|---|---|---|
| 1 | A | 903/909 (99%) | 825 (91%) | 77 (8%) | 1 (0%) | 53 | 85 |
| 1 | B | 900/909 (99%) | 816 (91%) | 82 (9%) | 2 (0%) | 49 | 82 |
| 1 | C | 905/909 (100%) | 816 (90%) | 87 (10%) | 2 (0%) | 49 | 82 |
| 1 | D | 904/909 (99%) | 838 (93%) | 65 (7%) | 1 (0%) | 53 | 85 |
| 1 | E | 904/909 (99%) | 835 (92%) | 68 (8%) | 1 (0%) | 53 | 85 |
| 1 | F | 902/909 (99%) | 829 (92%) | 73 (8%) | 0 | 100 | 100 |
| 1 | G | 904/909 (99%) | 834 (92%) | 68 (8%) | 2 (0%) | 49 | 82 |
| 1 | H | 903/909 (99%) | 822 (91%) | 79 (9%) | 2 (0%) | 49 | 82 |
| 1 | I | 904/909 (99%) | 829 (92%) | 73 (8%) | 2 (0%) | 49 | 82 |
| 1 | J | 904/909 (99%) | 841 (93%) | 61 (7%) | 2 (0%) | 49 | 82 |
| 1 | K | 906/909 (100%) | 822 (91%) | 82 (9%) | 2 (0%) | 49 | 82 |
| 1 | L | 904/909 (99%) | 833 (92%) | 67 (7%) | 4 (0%) | 36 | 73 |
| 2 | Q | 365/370 (99%) | 325 (89%) | 39 (11%) | 1 (0%) | 43 | 77 |
| 2 | R | 365/370 (99%) | 325 (89%) | 39 (11%) | 1 (0%) | 43 | 77 |
| 2 | S | 365/370 (99%) | 325 (89%) | 39 (11%) | 1 (0%) | 43 | 77 |
| 2 | T | 365/370 (99%) | 326 (89%) | 38 (10%) | 1 (0%) | 43 | 77 |
| 3 | O | 182/278 (66%) | 161 (88%) | 21 (12%) | 0 | 100 | 100 |
| 3 | P | 182/278 (66%) | 160 (88%) | 22 (12%) | 0 | 100 | 100 |
| 4 | N | 253/609 (42%) | 230 (91%) | 23 (9%) | 0 | 100 | 100 |
| 5 | M | 449/451 (100%) | 391 (87%) | 55 (12%) | 3 (1%) | 24 | 63 |
| All | All | 13369/14004 (96%) | 12183 (91%) | 1158 (9%) | 28 (0%) | 53 | 82 |

All (28) Ramachandran outliers are listed below:

| Mol | Chain | Res | Type |
|---|---|---|---|
| 1 | K | 789 | VAL |
| 1 | L | 173 | PRO |
| 1 | I | 347 | ILE |
| 1 | G | 347 | ILE |
| 1 | L | 347 | ILE |
| 1 | J | 347 | ILE |
| 1 | H | 261 | ASP |
| 1 | C | 31 | GLN |

*Continued on next page...*

CONFIDENTIAL VALIDATION REPORT

*Continued from previous page...*

| Mol | Chain | Res | Type |
|---|---|---|---|
| 1 | B | 673 | ASP |
| 5 | M | 359 | ALA |
| 1 | A | 658 | THR |
| 1 | L | 172 | GLN |
| 1 | K | 658 | THR |
| 1 | J | 300 | GLN |
| 1 | I | 254 | HIS |
| 2 | S | 163 | MET |
| 2 | R | 163 | MET |
| 1 | B | 300 | GLN |
| 2 | T | 163 | MET |
| 2 | Q | 163 | MET |
| 5 | M | 360 | VAL |
| 1 | L | 885 | ILE |
| 1 | C | 458 | PRO |
| 1 | H | 366 | VAL |
| 1 | D | 347 | ILE |
| 1 | G | 366 | VAL |
| 5 | M | 119 | PRO |
| 1 | E | 366 | VAL |

##### 5.3.2 Protein sidechains (i)

In the following table, the Percentiles column shows the percent sidechain outliers of the chain as a percentile score with respect to all PDB entries followed by that with respect to all EM entries.

The Analysed column shows the number of residues for which the sidechain conformation was analysed, and the total number of residues.

| Mol | Chain | Analysed | Rotameric | Outliers | Percentiles | |
|---|---|---|---|---|---|---|
| 1 | A | 775/777 (100%) | 763 (98%) | 12 (2%) | 67 | 86 |
| 1 | B | 774/777 (100%) | 762 (98%) | 12 (2%) | 65 | 85 |
| 1 | C | 776/777 (100%) | 766 (99%) | 10 (1%) | 71 | 87 |
| 1 | D | 776/777 (100%) | 767 (99%) | 9 (1%) | 74 | 88 |
| 1 | E | 775/777 (100%) | 764 (99%) | 11 (1%) | 69 | 87 |
| 1 | F | 775/777 (100%) | 764 (99%) | 11 (1%) | 69 | 87 |
| 1 | G | 775/777 (100%) | 763 (98%) | 12 (2%) | 67 | 86 |
| 1 | H | 775/777 (100%) | 763 (98%) | 12 (2%) | 67 | 86 |

*Continued on next page...*

CONFIDENTIAL VALIDATION REPORT

*Continued from previous page...*

| Mol | Chain | Analysed | Rotameric | Outliers | Percentiles | |
|---|---|---|---|---|---|---|
| 1 | I | 775/777 (100%) | 762 (98%) | 13 (2%) | 63 | 84 |
| 1 | J | 775/777 (100%) | 763 (98%) | 12 (2%) | 67 | 86 |
| 1 | K | 776/777 (100%) | 767 (99%) | 9 (1%) | 74 | 88 |
| 1 | L | 776/777 (100%) | 762 (98%) | 14 (2%) | 62 | 83 |
| 2 | Q | 301/304 (99%) | 300 (100%) | 1 (0%) | 93 | 97 |
| 2 | R | 301/304 (99%) | 300 (100%) | 1 (0%) | 93 | 97 |
| 2 | S | 301/304 (99%) | 300 (100%) | 1 (0%) | 93 | 97 |
| 2 | T | 301/304 (99%) | 300 (100%) | 1 (0%) | 93 | 97 |
| 3 | O | 158/234 (68%) | 155 (98%) | 3 (2%) | 60 | 83 |
| 3 | P | 158/234 (68%) | 155 (98%) | 3 (2%) | 60 | 83 |
| 4 | N | 226/520 (44%) | 221 (98%) | 5 (2%) | 55 | 81 |
| 5 | M | 406/406 (100%) | 399 (98%) | 7 (2%) | 63 | 84 |
| All | All | 11455/11934 (96%) | 11296 (99%) | 159 (1%) | 71 | 87 |

All (159) residues with a non-rotameric sidechain are listed below:

| Mol | Chain | Res | Type |
|---|---|---|---|
| 1 | A | 62 | ARG |
| 1 | A | 234 | ASN |
| 1 | A | 263 | ARG |
| 1 | A | 303 | ASN |
| 1 | A | 335 | ASN |
| 1 | A | 383 | ASN |
| 1 | A | 452 | ASN |
| 1 | A | 470 | ARG |
| 1 | A | 527 | ASN |
| 1 | A | 724 | ASN |
| 1 | A | 793 | ARG |
| 1 | A | 838 | ASN |
| 1 | L | 35 | ASN |
| 1 | L | 82 | ASN |
| 1 | L | 167 | ASN |
| 1 | L | 262 | ARG |
| 1 | L | 263 | ARG |
| 1 | L | 335 | ASN |
| 1 | L | 350 | ASN |
| 1 | L | 380 | MET |
| 1 | L | 410 | ASN |

*Continued on next page...*

CONFIDENTIAL VALIDATION REPORT

*Continued from previous page...*

| Mol | Chain | Res | Type |
|---|---|---|---|
| 1 | L | 655 | LEU |
| 1 | L | 684 | MET |
| 1 | L | 750 | ARG |
| 1 | L | 769 | ASN |
| 1 | L | 793 | ARG |
| 1 | K | 78 | ARG |
| 1 | K | 82 | ASN |
| 1 | K | 167 | ASN |
| 1 | K | 234 | ASN |
| 1 | K | 383 | ASN |
| 1 | K | 470 | ARG |
| 1 | K | 724 | ASN |
| 1 | K | 793 | ARG |
| 1 | K | 838 | ASN |
| 1 | J | 22 | ASN |
| 1 | J | 35 | ASN |
| 1 | J | 82 | ASN |
| 1 | J | 174 | ASN |
| 1 | J | 262 | ARG |
| 1 | J | 335 | ASN |
| 1 | J | 350 | ASN |
| 1 | J | 452 | ASN |
| 1 | J | 750 | ARG |
| 1 | J | 777 | ASN |
| 1 | J | 793 | ARG |
| 1 | J | 838 | ASN |
| 1 | H | 22 | ASN |
| 1 | H | 35 | ASN |
| 1 | H | 71 | ASN |
| 1 | H | 272 | ASN |
| 1 | H | 335 | ASN |
| 1 | H | 383 | ASN |
| 1 | H | 487 | ASN |
| 1 | H | 724 | ASN |
| 1 | H | 758 | ASN |
| 1 | H | 761 | ASN |
| 1 | H | 793 | ARG |
| 1 | H | 838 | ASN |
| 1 | I | 71 | ASN |
| 1 | I | 262 | ARG |
| 1 | I | 263 | ARG |
| 1 | I | 303 | ASN |

*Continued on next page...*

CONFIDENTIAL VALIDATION REPORT

*Continued from previous page...*

| Mol | Chain | Res | Type |
|---|---|---|---|
| 1 | I | 350 | ASN |
| 1 | I | 452 | ASN |
| 1 | I | 487 | ASN |
| 1 | I | 598 | ASN |
| 1 | I | 617 | ARG |
| 1 | I | 750 | ARG |
| 1 | I | 769 | ASN |
| 1 | I | 788 | ASN |
| 1 | I | 793 | ARG |
| 1 | G | 22 | ASN |
| 1 | G | 35 | ASN |
| 1 | G | 71 | ASN |
| 1 | G | 82 | ASN |
| 1 | G | 335 | ASN |
| 1 | G | 350 | ASN |
| 1 | G | 470 | ARG |
| 1 | G | 734 | ASN |
| 1 | G | 750 | ARG |
| 1 | G | 793 | ARG |
| 1 | G | 838 | ASN |
| 1 | G | 890 | ARG |
| 1 | F | 263 | ARG |
| 1 | F | 350 | ASN |
| 1 | F | 380 | MET |
| 1 | F | 383 | ASN |
| 1 | F | 527 | ASN |
| 1 | F | 583 | ASN |
| 1 | F | 684 | MET |
| 1 | F | 750 | ARG |
| 1 | F | 769 | ASN |
| 1 | F | 793 | ARG |
| 1 | F | 899 | ARG |
| 1 | E | 82 | ASN |
| 1 | E | 167 | ASN |
| 1 | E | 185 | ASN |
| 1 | E | 234 | ASN |
| 1 | E | 303 | ASN |
| 1 | E | 383 | ASN |
| 1 | E | 470 | ARG |
| 1 | E | 724 | ASN |
| 1 | E | 777 | ASN |
| 1 | E | 793 | ARG |

*Continued on next page...*

CONFIDENTIAL VALIDATION REPORT

*Continued from previous page...*

| Mol | Chain | Res | Type |
|---|---|---|---|
| 1 | E | 838 | ASN |
| 1 | D | 335 | ASN |
| 1 | D | 470 | ARG |
| 1 | D | 487 | ASN |
| 1 | D | 750 | ARG |
| 1 | D | 777 | ASN |
| 1 | D | 793 | ARG |
| 1 | D | 838 | ASN |
| 1 | D | 890 | ARG |
| 1 | D | 899 | ARG |
| 1 | C | 22 | ASN |
| 1 | C | 167 | ASN |
| 1 | C | 174 | ASN |
| 1 | C | 229 | ASN |
| 1 | C | 452 | ASN |
| 1 | C | 750 | ARG |
| 1 | C | 777 | ASN |
| 1 | C | 793 | ARG |
| 1 | C | 853 | ASN |
| 1 | C | 899 | ARG |
| 1 | B | 35 | ASN |
| 1 | B | 82 | ASN |
| 1 | B | 167 | ASN |
| 1 | B | 262 | ARG |
| 1 | B | 263 | ARG |
| 1 | B | 350 | ASN |
| 1 | B | 583 | ASN |
| 1 | B | 655 | LEU |
| 1 | B | 664 | THR |
| 1 | B | 750 | ARG |
| 1 | B | 769 | ASN |
| 1 | B | 793 | ARG |
| 2 | S | 194 | ASN |
| 2 | T | 194 | ASN |
| 2 | R | 194 | ASN |
| 2 | Q | 194 | ASN |
| 3 | P | 50 | ARG |
| 3 | P | 62 | ARG |
| 3 | P | 235 | ASN |
| 3 | O | 50 | ARG |
| 3 | O | 62 | ARG |
| 3 | O | 235 | ASN |

*Continued on next page...*

CONFIDENTIAL VALIDATION REPORT

*Continued from previous page...*

| Mol | Chain | Res | Type |
|---|---|---|---|
| 4 | N | 99 | ASN |
| 4 | N | 116 | ARG |
| 4 | N | 135 | ARG |
| 4 | N | 180 | ARG |
| 4 | N | 193 | ASN |
| 5 | M | 23 | ASN |
| 5 | M | 152 | ASN |
| 5 | M | 174 | ARG |
| 5 | M | 175 | ASN |
| 5 | M | 252 | LYS |
| 5 | M | 277 | LYS |
| 5 | M | 427 | ARG |

Some sidechains can be flipped to improve hydrogen bonding and reduce clashes. All (281) such sidechains are listed below:

| Mol | Chain | Res | Type |
|---|---|---|---|
| 1 | A | 67 | GLN |
| 1 | A | 86 | ASN |
| 1 | A | 117 | ASN |
| 1 | A | 234 | ASN |
| 1 | A | 303 | ASN |
| 1 | A | 335 | ASN |
| 1 | A | 336 | GLN |
| 1 | A | 369 | ASN |
| 1 | A | 423 | ASN |
| 1 | A | 511 | GLN |
| 1 | A | 537 | ASN |
| 1 | A | 724 | ASN |
| 1 | A | 725 | GLN |
| 1 | A | 829 | GLN |
| 1 | A | 838 | ASN |
| 1 | L | 35 | ASN |
| 1 | L | 82 | ASN |
| 1 | L | 86 | ASN |
| 1 | L | 137 | HIS |
| 1 | L | 167 | ASN |
| 1 | L | 301 | GLN |
| 1 | L | 335 | ASN |
| 1 | L | 350 | ASN |
| 1 | L | 359 | ASN |
| 1 | L | 410 | ASN |
| 1 | L | 414 | ASN |

*Continued on next page...*

CONFIDENTIAL VALIDATION REPORT

*Continued from previous page...*

| Mol | Chain | Res | Type |
|---|---|---|---|
| 1 | L | 423 | ASN |
| 1 | L | 445 | HIS |
| 1 | L | 508 | GLN |
| 1 | L | 527 | ASN |
| 1 | L | 537 | ASN |
| 1 | L | 574 | GLN |
| 1 | L | 645 | ASN |
| 1 | L | 724 | ASN |
| 1 | L | 728 | GLN |
| 1 | L | 769 | ASN |
| 1 | L | 783 | GLN |
| 1 | L | 829 | GLN |
| 1 | L | 856 | HIS |
| 1 | K | 9 | HIS |
| 1 | K | 82 | ASN |
| 1 | K | 117 | ASN |
| 1 | K | 167 | ASN |
| 1 | K | 234 | ASN |
| 1 | K | 359 | ASN |
| 1 | K | 369 | ASN |
| 1 | K | 423 | ASN |
| 1 | K | 559 | GLN |
| 1 | K | 561 | ASN |
| 1 | K | 724 | ASN |
| 1 | K | 725 | GLN |
| 1 | K | 743 | ASN |
| 1 | K | 783 | GLN |
| 1 | K | 815 | GLN |
| 1 | K | 829 | GLN |
| 1 | K | 838 | ASN |
| 1 | J | 22 | ASN |
| 1 | J | 35 | ASN |
| 1 | J | 82 | ASN |
| 1 | J | 117 | ASN |
| 1 | J | 159 | GLN |
| 1 | J | 186 | ASN |
| 1 | J | 201 | GLN |
| 1 | J | 287 | ASN |
| 1 | J | 291 | ASN |
| 1 | J | 335 | ASN |
| 1 | J | 350 | ASN |
| 1 | J | 359 | ASN |

*Continued on next page...*

CONFIDENTIAL VALIDATION REPORT

*Continued from previous page...*

| Mol | Chain | Res | Type |
|---|---|---|---|
| 1 | J | 495 | GLN |
| 1 | J | 537 | ASN |
| 1 | J | 613 | ASN |
| 1 | J | 777 | ASN |
| 1 | J | 788 | ASN |
| 1 | J | 829 | GLN |
| 1 | J | 838 | ASN |
| 1 | J | 859 | ASN |
| 1 | H | 9 | HIS |
| 1 | H | 22 | ASN |
| 1 | H | 35 | ASN |
| 1 | H | 71 | ASN |
| 1 | H | 86 | ASN |
| 1 | H | 117 | ASN |
| 1 | H | 272 | ASN |
| 1 | H | 335 | ASN |
| 1 | H | 369 | ASN |
| 1 | H | 383 | ASN |
| 1 | H | 487 | ASN |
| 1 | H | 511 | GLN |
| 1 | H | 561 | ASN |
| 1 | H | 574 | GLN |
| 1 | H | 588 | GLN |
| 1 | H | 589 | ASN |
| 1 | H | 598 | ASN |
| 1 | H | 724 | ASN |
| 1 | H | 758 | ASN |
| 1 | H | 761 | ASN |
| 1 | H | 783 | GLN |
| 1 | H | 815 | GLN |
| 1 | H | 829 | GLN |
| 1 | H | 838 | ASN |
| 1 | I | 71 | ASN |
| 1 | I | 86 | ASN |
| 1 | I | 117 | ASN |
| 1 | I | 137 | HIS |
| 1 | I | 141 | GLN |
| 1 | I | 172 | GLN |
| 1 | I | 182 | ASN |
| 1 | I | 185 | ASN |
| 1 | I | 303 | ASN |
| 1 | I | 350 | ASN |

*Continued on next page...*

CONFIDENTIAL VALIDATION REPORT

*Continued from previous page...*

| Mol | Chain | Res | Type |
|---|---|---|---|
| 1 | I | 445 | HIS |
| 1 | I | 452 | ASN |
| 1 | I | 487 | ASN |
| 1 | I | 574 | GLN |
| 1 | I | 588 | GLN |
| 1 | I | 598 | ASN |
| 1 | I | 724 | ASN |
| 1 | I | 769 | ASN |
| 1 | I | 771 | GLN |
| 1 | I | 788 | ASN |
| 1 | I | 829 | GLN |
| 1 | I | 856 | HIS |
| 1 | G | 22 | ASN |
| 1 | G | 28 | GLN |
| 1 | G | 35 | ASN |
| 1 | G | 71 | ASN |
| 1 | G | 82 | ASN |
| 1 | G | 86 | ASN |
| 1 | G | 117 | ASN |
| 1 | G | 141 | GLN |
| 1 | G | 159 | GLN |
| 1 | G | 172 | GLN |
| 1 | G | 182 | ASN |
| 1 | G | 185 | ASN |
| 1 | G | 201 | GLN |
| 1 | G | 291 | ASN |
| 1 | G | 335 | ASN |
| 1 | G | 350 | ASN |
| 1 | G | 359 | ASN |
| 1 | G | 369 | ASN |
| 1 | G | 414 | ASN |
| 1 | G | 574 | GLN |
| 1 | G | 725 | GLN |
| 1 | G | 734 | ASN |
| 1 | G | 751 | GLN |
| 1 | G | 783 | GLN |
| 1 | G | 829 | GLN |
| 1 | G | 838 | ASN |
| 1 | F | 67 | GLN |
| 1 | F | 86 | ASN |
| 1 | F | 287 | ASN |
| 1 | F | 350 | ASN |

*Continued on next page...*

CONFIDENTIAL VALIDATION REPORT

*Continued from previous page...*

| Mol | Chain | Res | Type |
|---|---|---|---|
| 1 | F | 383 | ASN |
| 1 | F | 387 | GLN |
| 1 | F | 445 | HIS |
| 1 | F | 527 | ASN |
| 1 | F | 537 | ASN |
| 1 | F | 559 | GLN |
| 1 | F | 561 | ASN |
| 1 | F | 574 | GLN |
| 1 | F | 583 | ASN |
| 1 | F | 598 | ASN |
| 1 | F | 698 | GLN |
| 1 | F | 769 | ASN |
| 1 | F | 829 | GLN |
| 1 | F | 887 | GLN |
| 1 | E | 9 | HIS |
| 1 | E | 31 | GLN |
| 1 | E | 82 | ASN |
| 1 | E | 117 | ASN |
| 1 | E | 167 | ASN |
| 1 | E | 185 | ASN |
| 1 | E | 186 | ASN |
| 1 | E | 234 | ASN |
| 1 | E | 303 | ASN |
| 1 | E | 309 | ASN |
| 1 | E | 369 | ASN |
| 1 | E | 383 | ASN |
| 1 | E | 423 | ASN |
| 1 | E | 511 | GLN |
| 1 | E | 724 | ASN |
| 1 | E | 777 | ASN |
| 1 | E | 838 | ASN |
| 1 | E | 856 | HIS |
| 1 | D | 86 | ASN |
| 1 | D | 117 | ASN |
| 1 | D | 141 | GLN |
| 1 | D | 172 | GLN |
| 1 | D | 201 | GLN |
| 1 | D | 269 | ASN |
| 1 | D | 287 | ASN |
| 1 | D | 301 | GLN |
| 1 | D | 335 | ASN |
| 1 | D | 383 | ASN |

*Continued on next page...*

CONFIDENTIAL VALIDATION REPORT

*Continued from previous page...*

| Mol | Chain | Res | Type |
|---|---|---|---|
| 1 | D | 438 | ASN |
| 1 | D | 445 | HIS |
| 1 | D | 487 | ASN |
| 1 | D | 645 | ASN |
| 1 | D | 725 | GLN |
| 1 | D | 751 | GLN |
| 1 | D | 777 | ASN |
| 1 | D | 783 | GLN |
| 1 | D | 829 | GLN |
| 1 | D | 838 | ASN |
| 1 | C | 22 | ASN |
| 1 | C | 117 | ASN |
| 1 | C | 167 | ASN |
| 1 | C | 201 | GLN |
| 1 | C | 229 | ASN |
| 1 | C | 287 | ASN |
| 1 | C | 369 | ASN |
| 1 | C | 445 | HIS |
| 1 | C | 495 | GLN |
| 1 | C | 537 | ASN |
| 1 | C | 574 | GLN |
| 1 | C | 688 | ASN |
| 1 | C | 725 | GLN |
| 1 | C | 751 | GLN |
| 1 | C | 777 | ASN |
| 1 | C | 783 | GLN |
| 1 | C | 829 | GLN |
| 1 | C | 853 | ASN |
| 1 | B | 35 | ASN |
| 1 | B | 67 | GLN |
| 1 | B | 82 | ASN |
| 1 | B | 141 | GLN |
| 1 | B | 167 | ASN |
| 1 | B | 287 | ASN |
| 1 | B | 350 | ASN |
| 1 | B | 387 | GLN |
| 1 | B | 527 | ASN |
| 1 | B | 537 | ASN |
| 1 | B | 724 | ASN |
| 1 | B | 725 | GLN |
| 1 | B | 728 | GLN |
| 1 | B | 769 | ASN |

*Continued on next page...*

CONFIDENTIAL VALIDATION REPORT

*Continued from previous page...*

| Mol | Chain | Res | Type |
|---|---|---|---|
| 1 | B | 783 | GLN |
| 1 | B | 829 | GLN |
| 2 | S | 14 | ASN |
| 2 | S | 194 | ASN |
| 2 | S | 208 | ASN |
| 2 | S | 218 | ASN |
| 2 | S | 272 | GLN |
| 2 | T | 194 | ASN |
| 2 | T | 208 | ASN |
| 2 | T | 252 | HIS |
| 2 | T | 272 | GLN |
| 2 | R | 11 | ASN |
| 2 | R | 194 | ASN |
| 2 | R | 208 | ASN |
| 2 | R | 218 | ASN |
| 2 | R | 272 | GLN |
| 2 | Q | 11 | ASN |
| 2 | Q | 194 | ASN |
| 2 | Q | 208 | ASN |
| 2 | Q | 218 | ASN |
| 2 | Q | 272 | GLN |
| 3 | P | 13 | GLN |
| 3 | P | 44 | GLN |
| 3 | P | 79 | GLN |
| 3 | P | 235 | ASN |
| 3 | O | 13 | GLN |
| 3 | O | 44 | GLN |
| 3 | O | 79 | GLN |
| 3 | O | 109 | GLN |
| 3 | O | 235 | ASN |
| 4 | N | 99 | ASN |
| 4 | N | 144 | GLN |
| 4 | N | 145 | HIS |
| 4 | N | 218 | ASN |
| 5 | M | 23 | ASN |
| 5 | M | 29 | GLN |
| 5 | M | 46 | ASN |
| 5 | M | 152 | ASN |
| 5 | M | 175 | ASN |
| 5 | M | 218 | ASN |
| 5 | M | 294 | GLN |
| 5 | M | 345 | HIS |

*Continued on next page...*

CONFIDENTIAL VALIDATION REPORT

*Continued from previous page...*

| Mol | Chain | Res | Type |
|---|---|---|---|
| 5 | M | 351 | GLN |
| 5 | M | 367 | GLN |
| 5 | M | 386 | GLN |

##### 5.3.3 RNA (i)

There are no RNA molecules in this entry.

#### 5.4 Non-standard residues in protein, DNA, RNA chains (i)

There are no non-standard protein/DNA/RNA residues in this entry.

#### 5.5 Carbohydrates (i)

There are no carbohydrates in this entry.

#### 5.6 Ligand geometry (i)

There are no ligands in this entry.

#### 5.7 Other polymers (i)

There are no such residues in this entry.

#### 5.8 Polymer linkage issues (i)

There are no chain breaks in this entry.

CONFIDENTIAL VALIDATION REPORT
